# Examining Discrimination Performance and Likelihood Ratio Values for Two Different Likelihood Ratio Systems Using the Provedit Dataset

**DOI:** 10.1101/2021.05.26.445891

**Authors:** Sarah Riman, Hari Iyer, Peter M. Vallone

## Abstract

The conventional capillary electrophoresis (CE) genotyping workflow used in forensic DNA laboratories is composed of two processes: measurement and interpretation. The outcome of the measurement process is an electropherogram (EPG). The outcome of the interpretation process is a strength of evidence statement often reported in the form of a likelihood ratio (LR) which typically requires probabilistic genotyping software (PGS). An LR system is defined as the entire pipeline of the measurement and interpretation processes where PGS is a piece of the whole LR system. To gain understanding on how two LR systems perform, a total of 154 two-person mixture, 147 three-person mixture, and 127 four-person mixture profiles of varying DNA quality, DNA quantity, and mixture ratios were obtained from the filtered (.CSV) files of the GlobalFiler 29 cycles 15s PROVEDIt dataset and deconvolved in two independently developed fully continuous programs, STRmix v2.6 and EuroForMix v2.1.0. Various parameters were set in each software and LR computations obtained from the two software were based on same/fixed EPG features, same pair of propositions, number of contributors, theta, and population allele frequencies. The ability of each LR system to discriminate between contributor (H1-true) and non-contributor (H2-true) scenarios was evaluated qualitatively and quantitatively. Differences in the numeric LR values and their corresponding verbal classifications between the two LR systems were compared. The magnitude of the differences in the assigned LRs and the potential explanations for the observed differences greater than or equal to 3 on the log_10_ scale were described. Cases of LR < 1 for H1-true tests and LR > 1 for H2-true tests were also discussed. Our intent is to demonstrate the value of using a publicly available ground truth known mixture dataset to assess discrimination performance of any LR system and show the steps used to investigate and understand similarities and differences between different LR systems. We share our observations with the forensic community and describe how examining more than one PGS with similar discrimination power can be beneficial, help analysts compare interpretation especially with low-template profiles or minor contributor cases, and be a potential additional diagnostic check even if software in use does contain certain diagnostic statistics as part of the output.

**Highlights:**

- The use of two different Likelihood Ratio (LR) systems to assign LRs is discussed.
- H1-true and H2-true tests are performed using STRmix and EuroForMix and a large set of PROVEDIt mixture profiles.
- Assessment of discrimination performance of two LR systems using ROC plots, scatter plots, and relative frequency histograms.
- The ability of the two LR systems to discriminate between contributors and non-contributors are statistically indistinguishable for the data that we considered.
- Potential reasons for the differences in LR values between the two LR systems that are ≥ 3 on the log_10_ scale are investigated and discussed.
- Contributors with LRs < 1 and non-contributors with LRs > 1 generated from each LR system are discussed.

## 1. Introduction

In recent years, the forensic DNA field has moved away from threshold-based interpretation approaches [1–5] and towards the implementation of fully continuous probabilistic genotyping software (PGS) [3, 5, 6]. PGS uses computer algorithms and complex calculations to apply biological, statistical, and mathematical models to resolve genotypes of contributors and/or assign evidential weight for the DNA typing results [6–9].

Fully continuous models, unlike binary and semi-continuous models, have caused a “paradigm shift in DNA profile interpretation” [9]. The models use quantitative information contained within a profile (e.g. allelic designations, peak heights, molecular weights/fragment length), take into account stochastic effects, model peak height variability, and allow interpretation of low-level and complex DNA mixtures, thus reducing subjective judgement and promoting objectivity during interpretation [2, 3, 9–13]. Numerous commercial and open-source software and freeware packages implementing fully continuous models have been developed [11, 14]. At the time of writing the available commercial software are: STRmix [2, 15], TrueAllele [16, 17], GenoProof Mixture 3 [18, 19], MaSTR [20], DNA- View Mixture Solution [21], and LiRa-HT [22, 23]. The available open-source software are EuroForMix [24, 25], CEESIt [26, 27], LikeLTD [28, 29], Kongoh [30, 31], and DNAmixtures [32, 33]. Other free fully-continuous models such as DNAxs 2.0/DNAStatistX [34, 35] and BulletProof [36] are available for users upon request or subscription. Differences exist among the programs in the way they model the distribution of allelic peak heights, stutter artifacts, mixture ratios, degradation, and stochastic events [2, 3, 37–41].

Most PGS require the assignment of two propositions, the prosecution proposition (H1) and defense proposition (H2) that include the specification of the number of contributors. Other parameters specific for each PGS are also required to deliver a key output, a Bayes factor, commonly referred to as the likelihood ratio (LR) [42, 43]. LR is the strength of the evidence in favor of H1 relative to H2. It is expressed as the ratio of two probabilities:

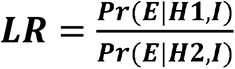

where *E* is the findings or evidence and *I* is the relevant background information. The numerator is the probability of the findings given that H1 and background information are true and the denominator is the probability of the findings given that H2 and background information are true [44, 45].

Several guidelines, recommendations, and reports [46–49] have emphasized the evaluation of PG models for use in DNA forensic casework not only by software developers but also by the end-users to test the validity, accuracy, and reliability, as well as identify and share the limitations of the software. Numerous scientific papers using empirically obtained data have described through intra- and inter-laboratory studies the performance and limitation of PGS in analyzing complex DNA profiles [3, 8, 12, 16, 50–56]. The behavior of LR was also studied under different conditions such as increasing profile complexity, using Polymerase Chain Reaction (PCR) replicates, assuming contributors, and adding incorrect or irrelevant information [57]. Several studies assessed the effect of assumptions leading to uncertainty on LR calculations such as number of contributors, propositions, population genetic models, allele frequencies, and co-ancestry coefficient (*F*_ST_ or θ) [6, 57–60]. Sources of variability pertaining to generating and interpreting DNA mixture profiles were also measured (e.g. PCR amplification, capillary electrophoresis load and injection, Markov chain Monte Carlo (MCMC) resampling method, and make up of allele frequency database) [58].

So far there is no consensus within the forensic DNA community on implementing a standardized continuous PGS [61, 62]. As a result, depending on the software being used, the analysis of the same DNA profile could yield different numeric LR values and, if used, different verbal characterizations [62, 63]. Even if the same PGS is used, the overall LR system could be different and hence will lead to different LRs [50, 55, 64]. Laboratories transitioning to PGS have compared the LR output from different statistical models: binary, semi-continuous, and fully continuous. The studies agree on the advantage of adopting fully continuous models that use peak height information since they have a better ability to distinguish between contributor (donor) and non-contributor (non-donor) and give larger LR values for the true contributors [1, 30, 52, 54, 65–70]. Using small datasets, a few select studies compared LR calculations across various fully continuous PGS and concluded that the models yielded similar LRs despite the differences among the PG modelling assumptions [29, 30, 52, 67]. However, the authors of a recent work demonstrated the impact of inter-model variability on numerical values and verbal expression of the LR when four variant models of the same continuous software, CEESIt, were compared [62].

In this study, we examine the discrimination performance and the assigned LR values for two different LR systems using two independently developed fully continuous PG models, STRmix v2.6 and EuroForMix (EFM) v2.1.0 [15, 25], and a large dataset composed of a range of complex two-person, three-person and four-person mixture profiles selected from PROVEDIt [71, 72]. STRmix and EFM were chosen out of the several mentioned software because they were readily available, and the authors were trained on how to use them. The focus of this study is not to suggest that any one of the software is based on a true or best model. Our intent is to:

- demonstrate the value of using publicly available ground truth known mixture data [72] to assess the performance of any LR system
- illustrate the steps used to investigate and understand any noteworthy discrepancies between different LR systems
- share our observations with the forensic community that can lead to improving one or both models
- describe how examining more than one PGS with similar discrimination power can be beneficial, help analysts compare interpretation especially with low-template profiles or minor contributor cases, and be a potential additional diagnostic check even if software in use does contain certain diagnostic statistics as part of the output
- address “Under what circumstances—and why—does the method produce results (random inclusion probabilities) that differ substantially from those produced by other methods?”, as recommended by the President’s Council of Advisors on Science and Technology (PCAST) report [49]

In the first part of this paper, we describe (i) the PROVEDIt dataset from which the mixture profiles were selected, (ii) the definition of an LR system, (iii) features and modeling assumptions of STRmix and EFM, and (iv) methods used to determine the parameters in each software. The remainder of the work focuses on the software discrimination performance, sample to sample profile LR comparisons, instances that resulted in LR < 1 for known contributors and LR > 1 for known non-contributors, and verbal classifications (as per Scientific Working Group on DNA Analysis Methods (SWGDAM) guidelines) of the reported numeric LR values.

## 2. Methods

### 2.1. PROVEDIt database description

The PROVEDIt (Project Research Openness for Validation with Empirical Data) is a comprehensive database openly available at lftdi.com and contains single source and complex mixture profiles generated from varying DNA quality, quantity, and mixture proportions. Different Short Tandem Repeat (STR) multiplexes, capillary electrophoresis (CE) instrument models, and injection times were used to generate the dataset which is available in three different data formats (Raw, Unfiltered, and Filtered) [71, 72].

In this study, the STR profiles used to set the PGS parameters and calculate the LRs were selected from the PROVEDIt dataset that was amplified with GlobalFiler (GF) kit (29 cycles) and analyzed on 3500 Genetic Analyzer with an injection time of 15 seconds (s). The profiles included one to four person mixtures (discussed in detail below), and were obtained from samples that were either left untreated (pristine DNA) or treated with different laboratory conditions (enzymatic degradation, UV irradiation, sonication-induced degradation, and PCR inhibition with humic acid). Both raw (.hid) and filtered (.CSV) files were used in the analysis. The filtered files present in the PROVEDIt database consist of the exported genotype tables containing allele designation, base pair (bp) size, and peak heights information for each sample profile analyzed in GeneMapper ID-X at an analytical threshold (AT) of one Relative Fluorescent Unit (RFU). Also, these filtered files did not contain artefacts such as pull-up, minus A, and – 2 bp in the SE33 locus as they were removed according to a defined criteria set by Alfonse et al. [72].

### 2.2. The LR system

The conventional CE genotyping workflow used in forensic DNA laboratories is composed of several steps that can be grouped into two processes: measurement and interpretation (Fig. 1A) [73]. The measurement process involves genomic DNA extraction, quantification, amplification using commercial multiplex STR kits (herein GF 29 cycles), and electrophoretic separation (herein 3500 at 15s injection time). The outcome of the measurement process is an electropherogram (EPG) composed of the length variants, heights, and sizes of the allelic and non-allelic peaks. The interpretation process involves data analysis. The outcome of the interpretation process is a strength of evidence statement often reported in the form of a LR and typically requires PGS.

**Fig. 1.**
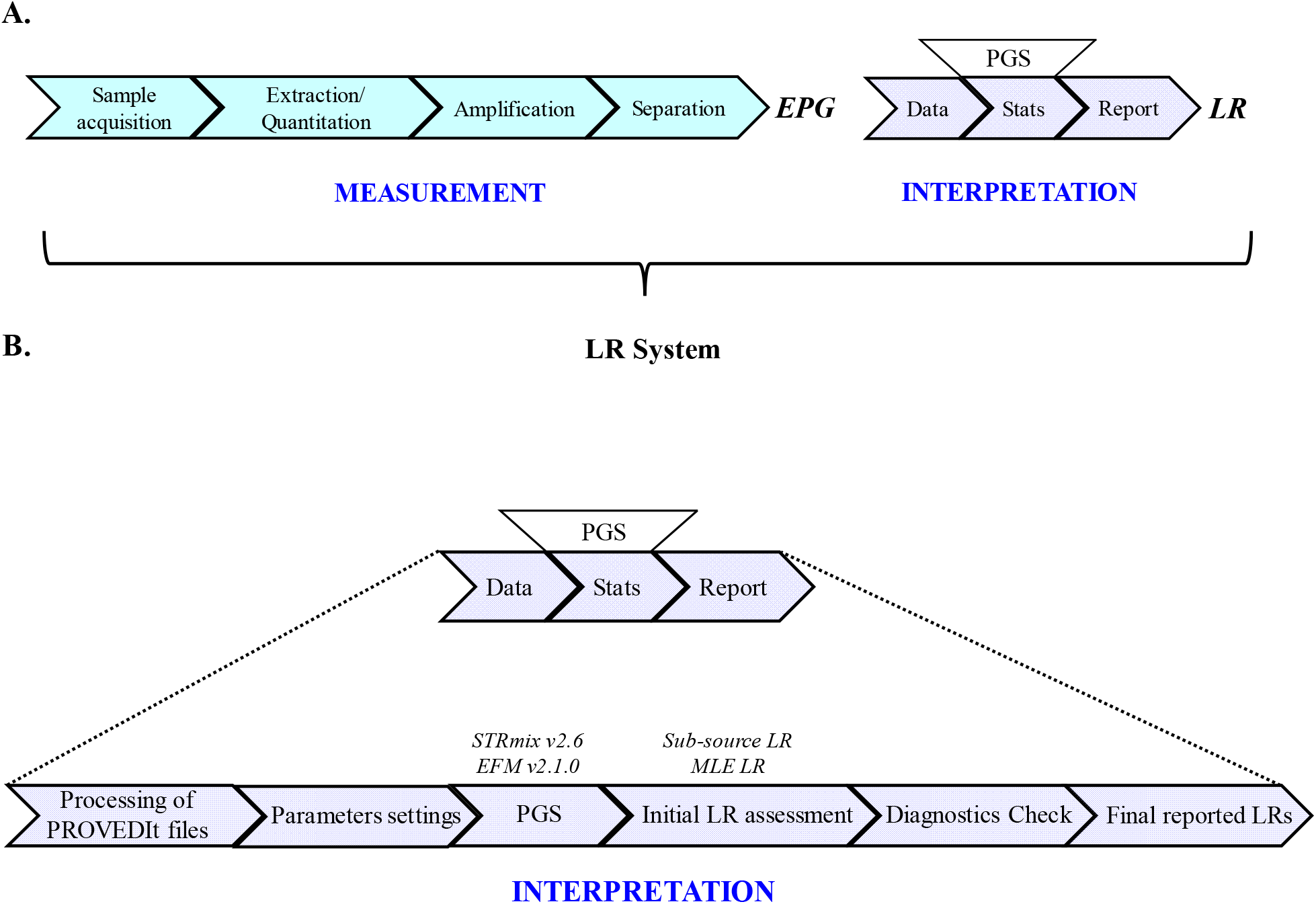
Schematic overview of the Likelihood Ratio (LR) system, adapted from [73, 74]. (A) The LR system is the entire pipeline of the two processes of the Capillary Electrophoresis (CE) genotyping workflow, measurement and interpretation. The Probabilistic Genotypic Software (PGS) is a piece of the whole LR system. (B) The performance assessment of the LR system in this study applies mainly to the interpretation process as the measurement process did not contain any variables in the explored dataset and was fixed throughout the study.

Our definition of the LR system is the entire pipeline starting from sample acquisition all the way to LR calculation. The PGS is a piece of the whole LR system. Therefore, performance assessment of the LR system is not only an assessment of the software but an assessment of the entire process.

Herein the measurement process was established by Alfonse et al. [72] as mentioned previously in Section 2.1 and therefore was fixed for both LR systems. Thus, the performance assessment in this study encompasses the interpretation process as shown in Fig. 1B that includes:

- our decision of using the filtered PROVEDIt files (files created according to the criteria in [72])
- our decision in processing the input filtered files (e.g., establishing and setting the AT values, removing Off-Ladder (OL) peaks, applying certain stutter models (discussed in Section 2.6), and importing same input files into both software though some stutter types were not modeled in EFM v2.1.0)
- the choice of setting the following input parameters:

- parameter values determined according to the chosen software (e.g. for STRmix: values determined for the variances, drop-in, stutter, and AT; for EFM: values determined for the AT and drop-in parameters)
- true number of contributors under the specified propositions
- NIST 1036-Caucasian allele frequencies and a θ correction of 0.01
- the choice of probabilistic genotyping software (herein STRmix v2.6 and EFM v2.1.0)
- the use and initial assessment of the sub-source LR values (labeled as sub-source LR in STRmix reports and maximum likelihood estimation (MLE) based LR in EFM reports)
- the check of diagnostics (review of per locus LR, deconvolution, genotypic weights, Gelman-Rubin statistics, log likelihood, and model selection)
- the reporting of the LRs

### 2.3. STRmix and EuroForMix (EFM)

STRmix [2, 15] and EFM [24, 25] implement fully continuous approaches and use graphical user interface for DNA profiles interpretation. Both carry out deconvolution of a DNA profile into sets of weighted genotypes and LR calculations.

In this study, the mixtures were interpreted using STRmix v2.6 [2, 15] and EFM v2.1.0 [24, 25]. In these versions, STRmix accounts for all the known stutter type artifacts into its modeling framework while EFM incorporates only back stutter model parameters. A summary of the different features and modeling assumptions of both software are shown in Table 1.

**Table 1.** Summary of different features and modelling assumptions of both software are related to versions used in this study [24, 69, 75, 76]. New features added to the updated versions are not covered here.

| Characteristics | *STRmix v2.6* | *EuroForMix v2.1.0* |
| --- | --- | --- |
| **Model** | Continuous | Continuous |
| **Software code** | Commercial written in Java | Open source written in R |
| **Inference approach** | Bayesian | MLE or Bayesian |
| **Graphical User Interface (GUI)** | ✓ | ✓ |
| **Calibration of single source samples** | ✓ | Possible |
| **Peak height distribution** | Log-normal | Gamma |
| **Stutter models** | Included for back, forward, double-back, and -2bp stutter types | Included for back stutter |
| **Drop-in model** | ✓ | ✓ |
| **Degradation model** | ✓ | ✓ |
| **Coancestry coefficient ($F_{ST}$ or $\theta$)** | ✓ | ✓ |
| **Allow multiple replicates** | ✓ | ✓ |
| **Different profiling kits and populations** | ✓ | ✓ |
| **Detection or analytical threshold** | Dye- or locus-specific | Global (overall) |
| **Relatedness model** | ✓ | ✓ |
| **Batch mode** | ✓ | Not available in the GUI |

### 2.4. Analytical thresholds

The analytical thresholds (ATs) are established to help discriminate between allelic and non-allelic signal (noise) [77]. AT values are also involved in setting drop-in parameters and modelling the drop-out probabilities [78, 79].

To determine the AT, 41 pristine single source DNA profiles with varying amounts of DNA template 0 ng – 0.5 ng were obtained from the filtered version (.CSV) of PROVEDIt files (GF 29 cycles/3500/15s injection time) [55, 71]. The list of the 41 samples selected for AT determination are detailed in Supplementary Table 1. Allelic, stutter, and other artifactual peaks were discarded from these profiles. The mean (µ) and standard deviation (σ) of the remaining peaks (noise observations) were estimated per dye-color channel. Then, AT was determined by substituting the values of µ and σ in the following equation: AT = μ + k* σ, where k was set to 10 [55, 77, 80–82]. The AT values were rounded up to the nearest multiple of 5 (Table 2) [55]. All peaks in the input profiles explored in this study with peak heights below the determined dye-specific AT values (shown in Table 2) were filtered out before importing the data into STRmix and EFM.

**Table 2.**
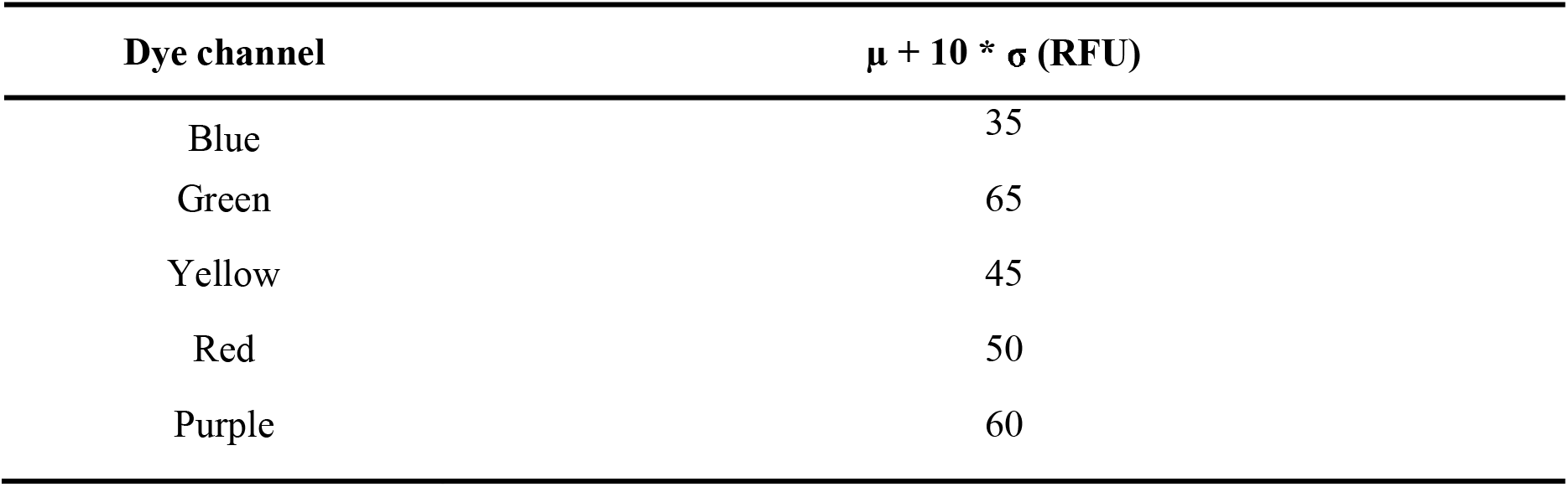
Optimized dye-specific Analytical Thresholds (ATs) used to filter out peaks with heights below these values prior to importing the DNA mixture profiles into each software. These per dye ATs were also set within STRmix v2.6 kit settings. EFM v2.1.0 implements an overall single AT value; hence, the lowest RFU value (35 RFU) was set within EFM.

| Dye channel | $\mu + 10 * \sigma$ (RFU) |
| --- | --- |
| Blue | 35 |
| Green | 65 |
| Yellow | 45 |
| Red | 50 |
| Purple | 60 |

STRmix and EFM require users to set the determined AT values in the detection threshold settings within each software before proceeding with LR calculations. STRmix v2.6 allows the user to specify either dye- or locus-specific analytical thresholds within the software. We chose to set the dye-specific ATs in STRmix as determined empirically and shown in Table 2. EFM v2.1.0 allows the user to set an overall single AT value [79]. The lowest RFU value (35 RFU) was used as the AT parameter in the EFM software.

### 2.5. Drop-in

Raw data (.hid files) of a set of 189 negative control profiles (listed in Supplementary Table 2) from PROVEDIt database (GF 29 cycles/3500/15s injection time) were analyzed in GeneMapper ID-X v1.5 with a 35 RFU for all the dye channels. The selected profiles that were amplified with no DNA (0 ng) resulted in 7 drop-in events that were observed and recorded with their corresponding occurrences and heights.

For STRmix, the drop-in frequency and drop-in cap parameters were determined and entered in the software. The frequency of the observed drop-in events was determined by using the drop-in worksheet available on STRmix support website [78]. Per instruction, the information entered into the worksheet was: the AT value used to analyze the negative controls, the total number of observed drop-in events with their corresponding occurrences and peak height information, the number of negative controls analyzed, and the number of loci in the GlobalFiler multiplex. The drop-in cap (in RFU) is based on the maximum observed drop-in height in the analyzed data and is advised to be set at a value that is greater than the maximum peak height of the observed drop-in peak [55, 78]. The highest drop-in peak observed in this study was 101 RFU. To set the drop-in cap at a value that is greater than the 101 RFU, we calculated the µ and σ of the peak heights of the observed drop-in events and substituted these values in the following equation: Drop-in cap = µ + k* σ, where k was set to ≈ 5. The drop-in cap was then rounded up to the nearest multiple of 5. The determined values of drop-in frequency and drop-in cap (shown in Table 3) were both set into the STRmix v2.6 software. Also, within STRmix v2.6, drop-in is modelled using either a uniform or a gamma distribution depending on the total number of observed drop-in peaks [78, 83]. Due to the few drop-in events (only 7) observed within our analysis from these PROVEDIt data, a uniform distribution was selected in the software.

**Table 3.** The drop-in parameters determined for STRmix v2.6 and EFM v2.1.0.

| Software | Drop-in parameters |
| --- | --- |
| *STRmix v2.6* | Drop-in cap = 180 RFU<br>Drop-in frequency = 0.0015 |
| *EFM v2.1.0* | Drop-in probability = 0.0015<br>Drop-in hyper-parameter ($\lambda$) = 0.032 |

EFM requires the setting of the drop-in parameters, the drop-in probability (C), and the hyper- parameter (l), that are also obtained from the analysis of the negative control profiles. Here, C was determined using C = n/N*L, where C is the drop-in probability per marker, n is the number of drop-in events, N is the number of samples used to count the number of drop-ins, and L is the number of markers in each sample used to count number of drop-ins. The estimated hyper-parameter (l) was determined using l = n/S_i_(x_i_ -T), where T is the analytical threshold used for analyzing drop-in, x_i_ is the peak height of each drop-in observed, and n is the number of drop-in events [25, 66, 79]. The determined values of C and l (shown in Table 3) were set in EFMv2.1.0 to model the drop-in peak heights.

### 2.6. Stutter

Both STRmix and EFM model stutter peaks. STRmix v2.6 models the following types of stutter: back/N-1 (B1), forward/N+1 (F1), double-back/N-2 (B2), and - 2bp stutter at SE33 and D1S1656, and implements a ‘per allele’ stutter ratio [37, 84]. STRmix estimates expected peak heights during interpretation by using the stutter exceptions and regression files. Typically, stutter files are prepared from empirical data of the STR kit in use [78, 83]. In this study, stutter models in STRmix (for B2, B1, and F1) were applied when assessing LR. All the mixture profiles analyzed herein did not contain stutter peaks at the – 2bp position. We used the stutter files that already exist within the software from a previously validated GF 29 cycle kit [55, 85]. EFM v2.1.0 models only back stutter by including an expected stutter proportion parameter [24, 66, 79]. Stutter types chosen to be modelled in STRmix (B1, F1, and B2) were retained in the input files after applying the AT values, and imported in both software even though F1 and B2 were not modelled in EFMv2.1.0. Any unmodelled stutter can also be explained as drop-in allelic events [29].

### 2.7. Saturation threshold

STRmix implements a saturation setting that is specific for the model of CE instrument in use. The setting accounts for camera saturation caused by excess amount of input DNA and high-level fluorescence. It is defined as the maximum RFU allowed in STRmix above which modeling of peak heights and DNA amounts will no longer be accurate. STRmix recommends determining the threshold from single source samples amplified with excess amount of DNA template (≥ 2 ng) [55, 78, 83].

In this study, the point of saturation was not determined because all the PROVEDIt profiles analyzed herein were amplified with DNA amount of < 1 ng and none of the observed data appeared to be approaching saturation levels. As per Kelly et al. [55], a saturation value of 30,000 RFU (commonly observed with ABI 3500 data) was set in the software kit settings.

### 2.8. Variance parameters

Model maker, a function in STRmix, models and determines the stutter and allele peak height variance constants (*k*^2^ for stutter peaks and *c*^2^ for allelic peaks) and the locus-specific amplification efficiency variance (LSAE) from single source profiles with known genotypes [8, 55, 78, 83].

Single source profiles (n = 333) obtained from the GF 29 cycles/3500/15s PROVEDIt database (filtered CSV files) were analyzed at an analytical threshold of 10 RFU at all the dye channels to maximize stutter observations of all the stutter types being modelled. To capture the range of peak height variability, the calibration set contained profiles amplified with varying DNA amount and quality [8, 55, 78, 83]. A detailed description of the quality and quantity of the samples used in the calibration set is listed in Supplementary Table 3. Prior to running the Model Maker function, an AT of 10 RFU was set in the software. The drop-in, stutter, and saturation parameters were set to the optimized values as discussed in Sections 2.5, 2.6, and 2.7, respectively.

The α, β parameters and mode describing the gamma distribution (Gamma (α, β)) of the allele peak height variance (*c*^2^) and stutter peak height variances (*k*^2^), and the mean of the LSAE variance derived from the Model Maker analysis are shown in Table 4 and were set into the software prior to the interpretation of the DNA mixture profiles.

**Table 4.**
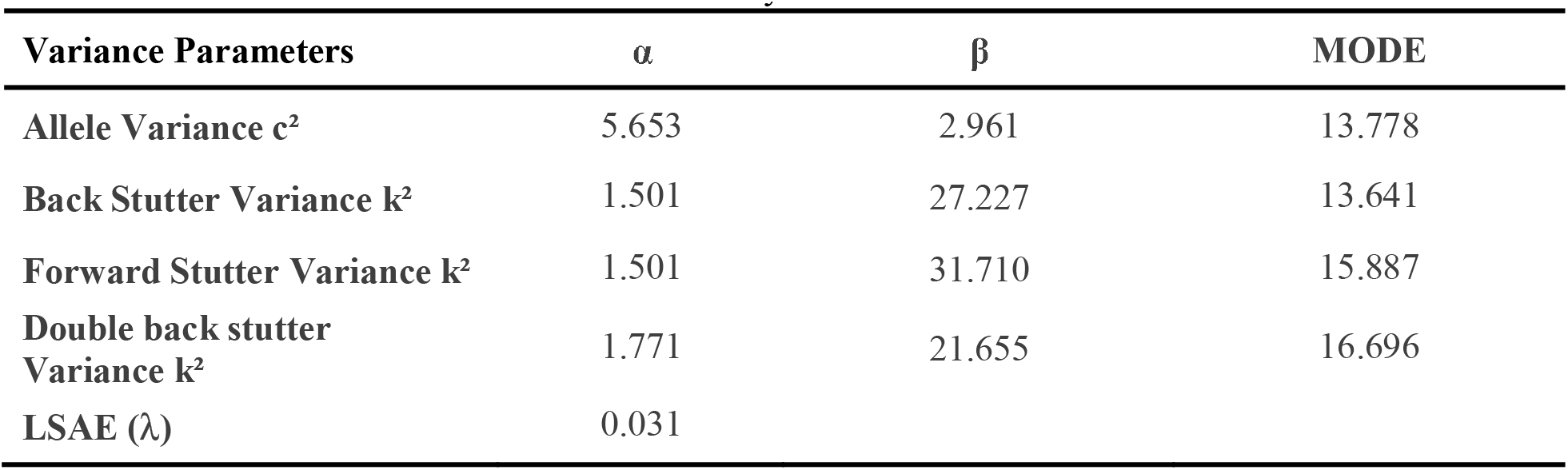
Peak height variance parameters for the allelic and different stutter peak types modelled by gamma distributions, along with the LSAE variance parameter used within STRmix were determined via Model Maker function for the GlobalFiler 29 cycles 15s PROVEDIt dataset.

### 2.9. PROVEDIt DNA mixture profiles

A total of 154 two-person (2P), 147 three-person (3P), and 127 four-person (4P) mixture profiles (Table 5) were obtained from the filtered (.CSV) files of the GlobalFiler 29 cycles 15s PROVEDIt dataset and used for LR calculations. The 2P, 3P, and 4P testing sets were prepared using DNA from 22 individuals for whom reference profiles were also available. The profiles used had varying: (1) minor contributor template amounts, (2) total input template amounts, (3) contributor ratios, and (4) DNA quality. A detailed description of the 2P, 3P, and 4P profiles that were used in the study is shown in Supplementary Table 4.

**Table 5.** Summary of the total number of PROVEDIt mixture profiles and H1-true and H2-true propositions analyzed in both STRmix v2.6 and EFM v2.1.0 for 2P, 3P, and 4P mixtures. POI indicates the person of interest that can be either known contributor or known non-contributor. U1, U2, U3, and U4 indicate one, two, three, or four unknown, unrelated individual(s) to the mixtures. For each mixture, we performed as many known contributor LR analysis (H1-true tests) as there are contributors to each mixture. For each contributor analysis, a non-contributor LR analysis (i.e. single H2-true test) was also performed using real (true-genotype) profiles randomly chosen from NIST 1036 US population dataset [86].

| Number of contributors | Number of mixtures | Propositions | Number of H1-true tests | Number of H2-true tests |
| --- | --- | --- | --- | --- |
| 2P | 154 | H1: POI + U1<br>H2: U1 + U2 | 308 | 308 |
| 3P | 147 | H1: POI + U1 + U2<br>H2: U1 + U2 + U3 | 441 | 441 |
| 4P | 127 | H1: POI + U1 + U2 + U3<br>H2: U1 + U2 + U3 + U4 | 508 | 508 |

The mixture input files were analyzed using the per dye specific ATs shown in Table 2 and converted along with person of interest (POI) files into a format specific to each software [79, 83]. Non- numeric values, “OL” peaks, were eliminated from all the analysis [83].

### 2.10. LR calculations and data analysis

The strength of evidence was assessed in both STRmix v2.6 [2, 15] and EFM v2.1.0 [24, 25] after setting parameters specific to each software as summarized in Table 6. STRmix interpretations were undertaken using the recommended MCMC parameters (default settings) of eight chains of 100,000 burn-in and 50,000 post burn-in accepts per chain [83]. In follow up analyses two interpretations were repeated with an increase in the number of accepts (1,000,000 burn-in and 500,000 post burn-in accepts per chain) to allow each of the chains to explore more possibilities in the probability space [87]. The reported sub-source LRs within the STRmix reports were considered for the analysis in this study.

**Table 6.**
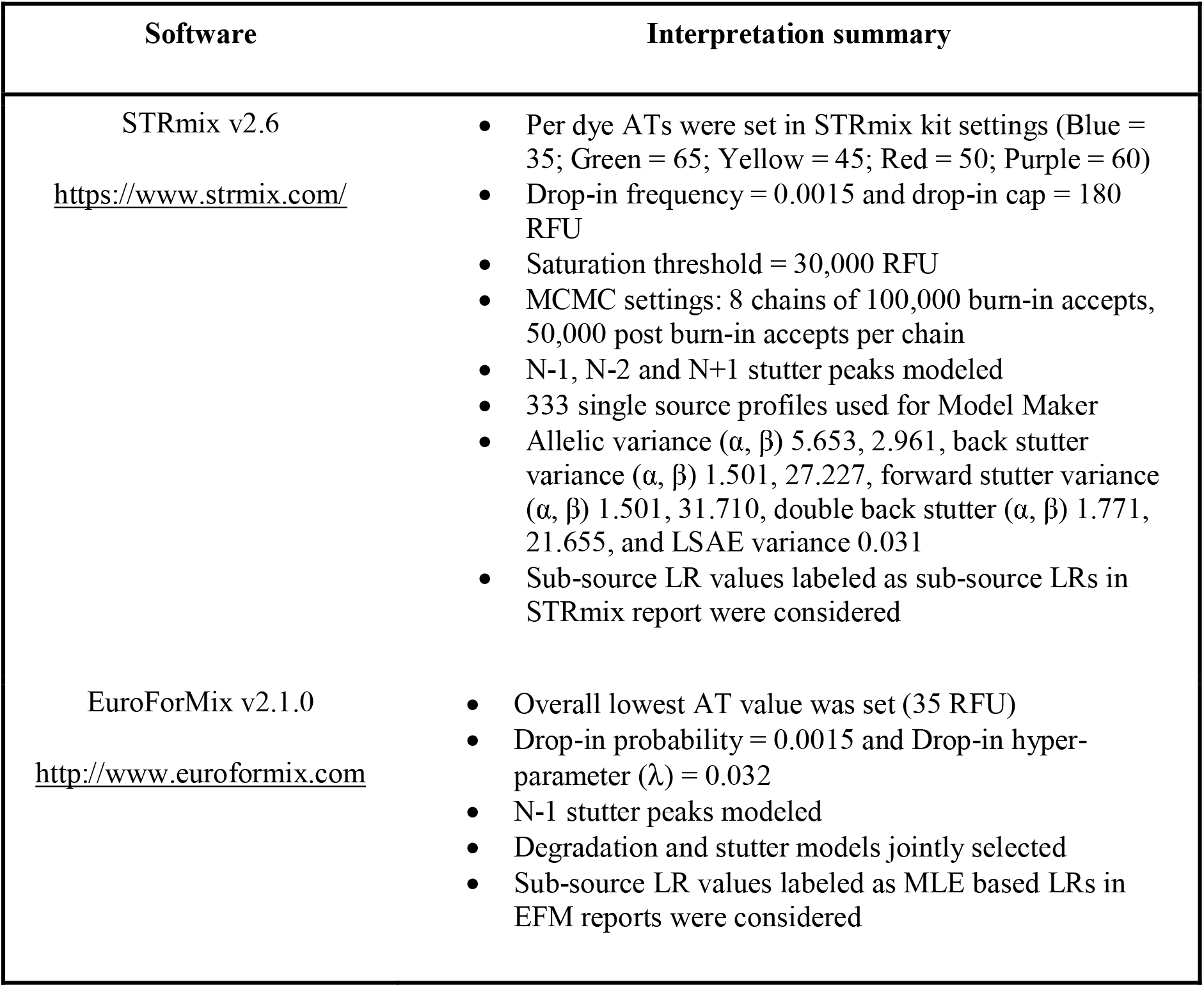
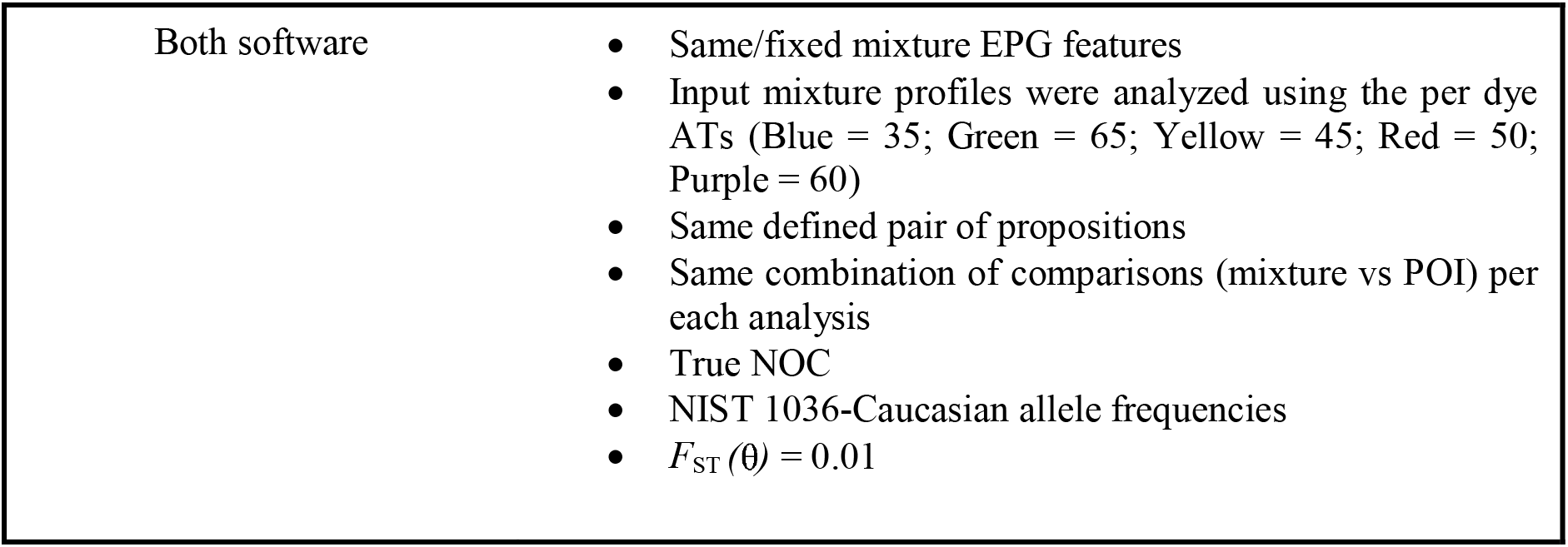
Summary of STRmix v2.6 and EFM v2.1.0 interpretation parameters and reported LR values

LR calculations in EFM were performed using the maximum likelihood estimate (MLE) method with both the degradation and stutter statistical models jointly turned on and included in all the EFM analysis. The reported sub-source LRs within the EFM labeled as MLE based LRs were used in the data analysis.

Both STRmix v2.6 and EFM v2.1.0 require the number of contributors (NOC) to be assigned to a profile prior to interpretation. The true NOC (ground truth) was specified in the settings of the software for each mixture profile that was interpreted. Each of the PROVEDIt mixture profile was compared to the appropriate known contributors (Supplementary Table 4) and known non-contributors (Supplementary Table 5). The known non-contributors were real (true-genotype) profiles randomly selected from the NIST 1036 US population dataset [86].

The propositions (Table 5) considered were:

H1: the DNA originated from the POI (either known contributor or known non-contributor) and N-1 unknown, unrelated individual(s).

H2: the DNA originated from N unknown, unrelated individuals; where N represented the true number of contributors (NOC).

All LRs were calculated using the NIST 1036-Caucasian allele frequencies [86]. To adjust for sub-population structuring, a *F*_ST_ of 0.01 was set in both software [41, 88]. Each software resulted in 308 LRs for H1-true tests and 308 LRs for H2-true tests for the 2P analysis, 441 LRs for H1-true tests and 441 LRs for H2-true tests for the 3P analysis, and 508 LRs for H1-true tests and 508 LRs for H2-true tests for the 4P analysis (outlined in Table 5). All the LR values yielded from both software are reported in log_10_ scale in Supplementary Table 4 (log_10_(LRs) for H1-true tests) and Supplementary Table 5 (log_10_(LRs) for H2-true tests) with the corresponding combination of comparisons (mixture vs POI). The profile LRs and the per-locus LRs assigned by STRmix and EFM were for the 21 autosomal STR markers only. LR assessment for the gender and Y-STR markers, Amelogenin, Y-indel, and DYS391, were not considered by either software.

### 2.11. Empirical Receiver Operating Characteristic (ROC) plots

An empirical ROC plot associated with a metric that is used to discriminate between two groups (for instance true contributors/true non-contributors) is a two-dimensional graph generated by plotting the rate of true positives (TP) in the empirical dataset versus the rate of false positives (FP) in the empirical dataset at each given threshold value for decision-making [89]. An illustration of an ROC plot is shown in Fig. 2.

**Fig. 2.**
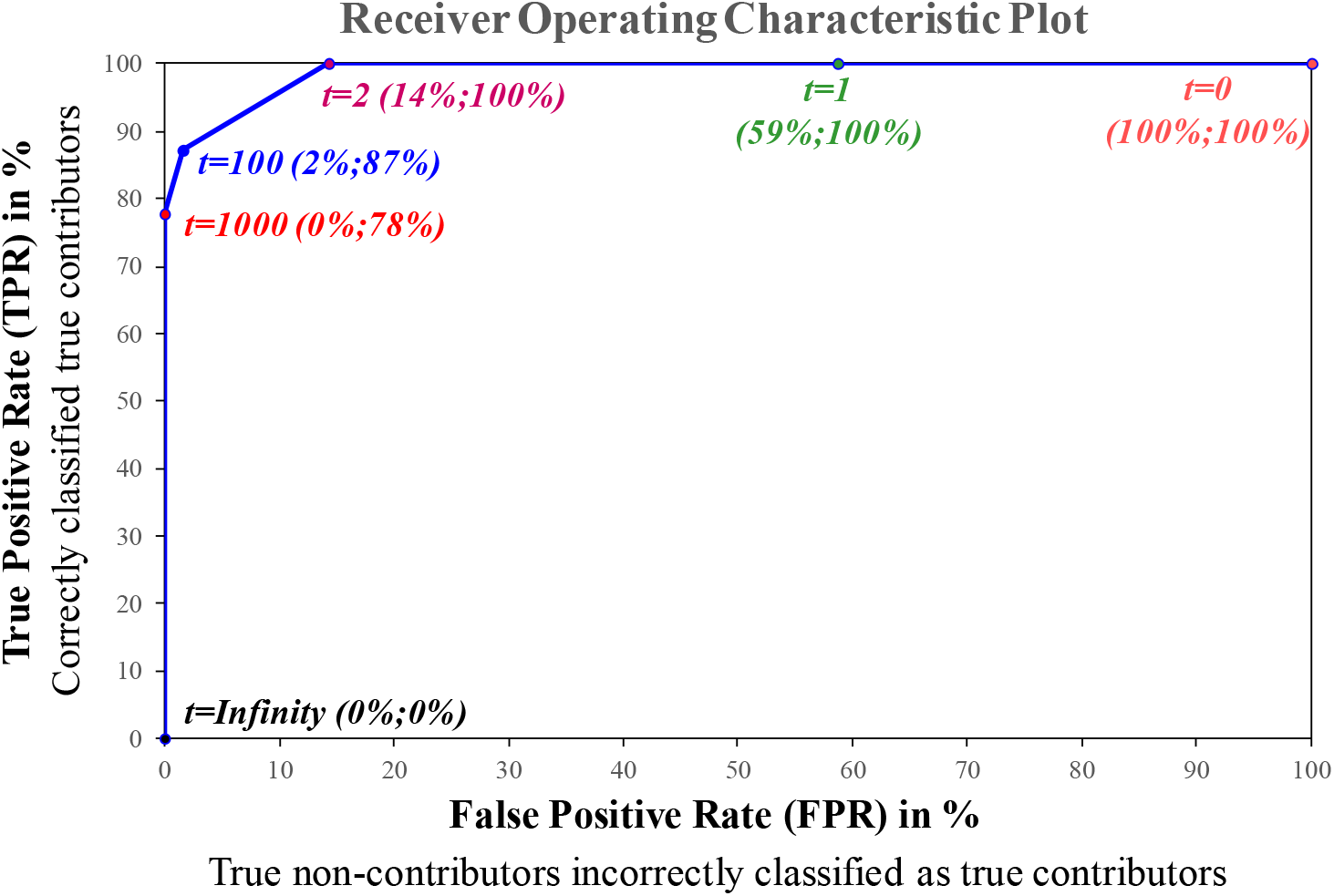
Receiver Operating Characteristic plot or (ROC plot). An illustration of an ROC plot used to evaluate the discrimination performance of an LR system. At a given LR threshold value *(t)*, the rates of the correctly classified true contributors (TPR) and the rates of true non-contributors incorrectly classified as true contributors (FPR) are plotted on the y-axis and x-axis, respectively. The ROC captures all the LR thresholds (*t*) simultaneously. Specific (*t)* values of 0, 1, 2, 100, 1000, and infinity are highlighted in this illustration. For every possible (*t)* value, a (FPR; TPR) value is recorded. As (*t)* value increases, the rate of both TP and FP decreases.

LR values from H1-true tests and H2-true tests were combined across each NOC level (2P, 3P, and 4P) generated from each software (STRmix and EFM), thus creating six datasets: STRmix 2P, EFM 2P, STRmix 3P, EFM 3P, STRmix 4P, and EFM 4P. To construct the ROCs shown in Section 3.3, a series of thresholds were applied to each of the six datasets. These thresholds corresponded to all the actual consecutive LR values observed in each dataset. For every possible threshold, two outcome measures or coordinates (FPR, TPR) were created. Herein, known contributors were considered as the reference standard. Hence, an event was classified and counted as a TP, when the LR values of the known contributors were greater than (>) a given threshold value. Therefore, TPR represented the counts of the true contributors of which LR values were > a given threshold value divided by the total counts of the known contributors in the considered dataset. FPR represented the counts of the known non-contributors with LR values > a given threshold value divided by the total counts of known non-contributors in the considered dataset. The TPR represented the rates of the correctly classified true contributors at a given threshold. FPR represented the rates of true non-contributors incorrectly classified as true contributors at the given threshold. TPR and FPR were plotted on the y-axis and x-axis, respectively. Plotting all these coordinates generated the ROCs that captured all the thresholds simultaneously. The points in the ROC plot were connected by line segments for visualization purposes. A point that is closest to the top left corner on the ROC plot has a high number of true positives, and low number of false positives [90]. True negative rates (TNR) and false negative rates (FNR) can also be deduced from the plotted ROCs. TNR are the rates of the correctly classified non-contributors. FNR are the rates of true contributors falsely labeled as non-contributors.

All data analysis and visualization discussed were conducted using the open source software **R** [91].

## 3. Results and discussion

### 3.1. Empirical assessment of LR systems using discrimination performance of H1-true and H2-true LR distributions

The ability of the assigned LR values to discriminate between H1-true scenarios and H2-true scenarios have been studied within STRmix and EFM by several authors using different experimental data. Various graphical tools were used to display and visualize these LR values such as scatter plots, violin plots, and density plots [8, 9, 51, 54, 66, 68, 92, 93].

LRs obtained from H1-true tests and H2-true tests are known to be affected by the information content in DNA mixture profiles. For example, the NOC, profile quality, minor template amount and/or total template amount, assumption of known contributors under both propositions, and replicate amplifications are factors that influence LR values and ultimately the discrimination performance of LR systems. These observations have been explored and documented by Taylor [57] and demonstrated by numerous publications [8, 9, 51, 54, 66, 68, 92, 93].

Although performance-based studies have been conducted for both STRmix and EFM in [8, 9, 51, 54, 66, 68, 92, 93] as mentioned above, we wanted to perform a diagnostic check to examine if the empirical LR results obtained in our study are consistent with expectations observed with previous work [8, 9, 51, 54, 66, 68, 92, 93], examine the overall performance of the two methods, and also conduct a case-by-case assessment of the assigned LR values between the two LR systems. As described in Sections 2.9 and 2.10 and shown in Supplementary Tables 4 and 5, mixtures from PROVEDIt database [71, 72] constructed from two, three, and four contributors at varying mixture proportions, DNA quality, and DNA template amounts were used to assess LR values within STRmix and EFM using both known contributors (H1-true or sensitivity tests) and randomly chosen known non-contributors (H2-true or specificity tests). The distributions of the assigned log_10_(LR) values were plotted as function of NOC (2P, 3P, and 4P), propositions (H1 and H2), and software (STRmix and EFM) (Fig. 3). The overall distribution plot shown in Fig. 3 was further broken down by varying mixture ratios (Additional file 1) and different DNA treatments used to compromise the DNA quality of the samples (DNA damage, DNA degradation, and PCR inhibition) (Additional file 2).

**Fig. 3.**
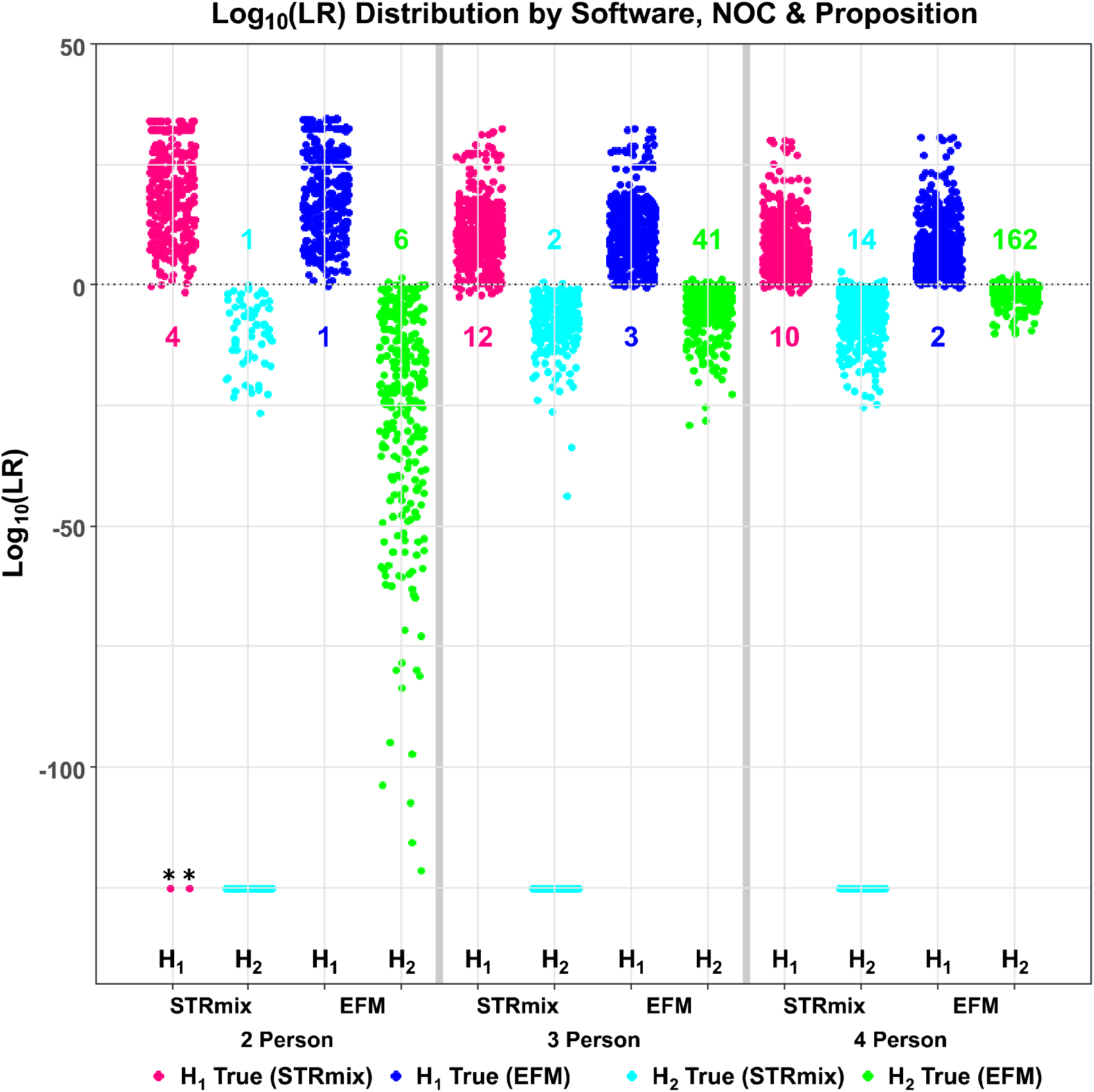
Distribution of log_10_(LR) values for H1-true and H2-true tests assessed by STRmix and EFM for two, three, and four person mixtures. The x-axis shows the labels of propositions (H1 and H2), software (STRmix and EFM), and the NOC = 2 Person, 3 Person, and 4 Person. LRs are plotted on the y-axis as log_10_(LR) values. Log_10_(LR) values obtained in STRmix (shown in magenta) and EFM (shown in blue) when two, three, and four person mixtures were compared to known contributors (H1-true or sensitivity test). Log_10_(LR) values obtained in STRmix (shown in cyan) and EFM (shown in green) when two, three, and four person mixtures were compared to known non-donors (H2-true or specificity test). All samples from different mixture ratios, total DNA template amounts, and DNA treatments are built into this global/overall distribution plot. The plot contains a total of 308 H1-true tests and 308 H2-true tests for the 2P analysis, 441 H1-true and 441 H2-true calculations for the 3P analysis, and 508 H1-true and 508 H2-true tests for the 4P mixtures. The central horizontal line plotted at log_10_(LR) = 0 represents neutrality. Results above and below the zero-line support inclusion and exclusion, respectively. The number of known contributor analyses that yielded log_10_(LRs) < 0 within STRmix and EFM are shown in magenta and blue for the 2P, 3P, and 4P and are also indicated with their corresponding log_10_(LRs) in Fig. 4 and Supplementary Table 6 (discussed in Section 3.2). The number of known non-contributor analyses that yielded log_10_(LRs) > 0 within STRmix and EFM are shown in cyan and green for the 2P, 3P, and 4P and are also indicated with their corresponding log_10_(LRs) in Fig. 4 and Supplementary Table 7 (discussed in Section 3.2). STRmix provides an LR value of 0 for excluded loci resulting in profile LR of 0, while EFM gives a non-zero LR value (generally very close to zero). Profiles with LR results of 0 from STRmix are plotted at −125 on the log_10_ scale. ** Two H1-true test interpretations of 2P mixtures for which STRmix assigned profile LRs of 0 (plotted at H1 true STRmix NOC = 2 Person in magenta at −125 on the log_10_ scale and discussed in detail in Section 3.6).

The magenta and blue data points are the log_10_(LRs) of the H1-true tests generated in STRmix and EFM, respectively. Log_10_(LRs) of the H2-true tests assigned by STRmix and EFM are shown in cyan and green, respectively (Fig. 3, and Additional files 1 and 2). The distribution of log_10_(LRs) from the H1- true tests is well separated from the distribution of log_10_(LRs) from the H2-true tests when the quality and DNA template amount of the contributor or total template amount of the samples are sufficiently high and the NOC in a mixture profile is low. As the quality and template amount per contributor of interest or mixture profile decreases and/or the NOC increases, log_10_(LRs) assigned from H1-true tests and H2-true tests become less discriminatory and trend downwards and upwards towards 0 (horizontal line), respectively, (Fig. 3, Additional files 1 and 2). Furthermore, as expected, when the distinction between the major-minor contributions to the same mixture increases so does the LRs of the major contributors as opposed to mixtures with equal contributor proportions (Additional file 1). As expected, the latter have lower LRs since information content associated with peak heights is limited or has no effect on LR calculations [9, 65, 68–70].

The magenta and blue data points below the central dashed horizontal line plotted at log_10_(LR) of zero in Fig. 3 and Additional files 1 and 2, correspond to the analyses of known contributors within STRmix and EFM that yielded log_10_(LRs) < 0 (adventitious exclusionary LRs). Cyan and green points above the horizontal line at log_10_(LR) = 0 in Fig. 3 and Additional files 1 and 2 are instances of H2-true tests that yielded log_10_(LRs) > 0 (adventitious inclusionary LRs). The number of these adventitious inclusionary and exclusionary LR instances are indicated in Fig. 3. These profiles are also presented with their corresponding log_10_(LRs) in Fig. 4 and Supplementary Tables 6 and 7 and are discussed in further details in Section 3.2.

**Fig. 4.**
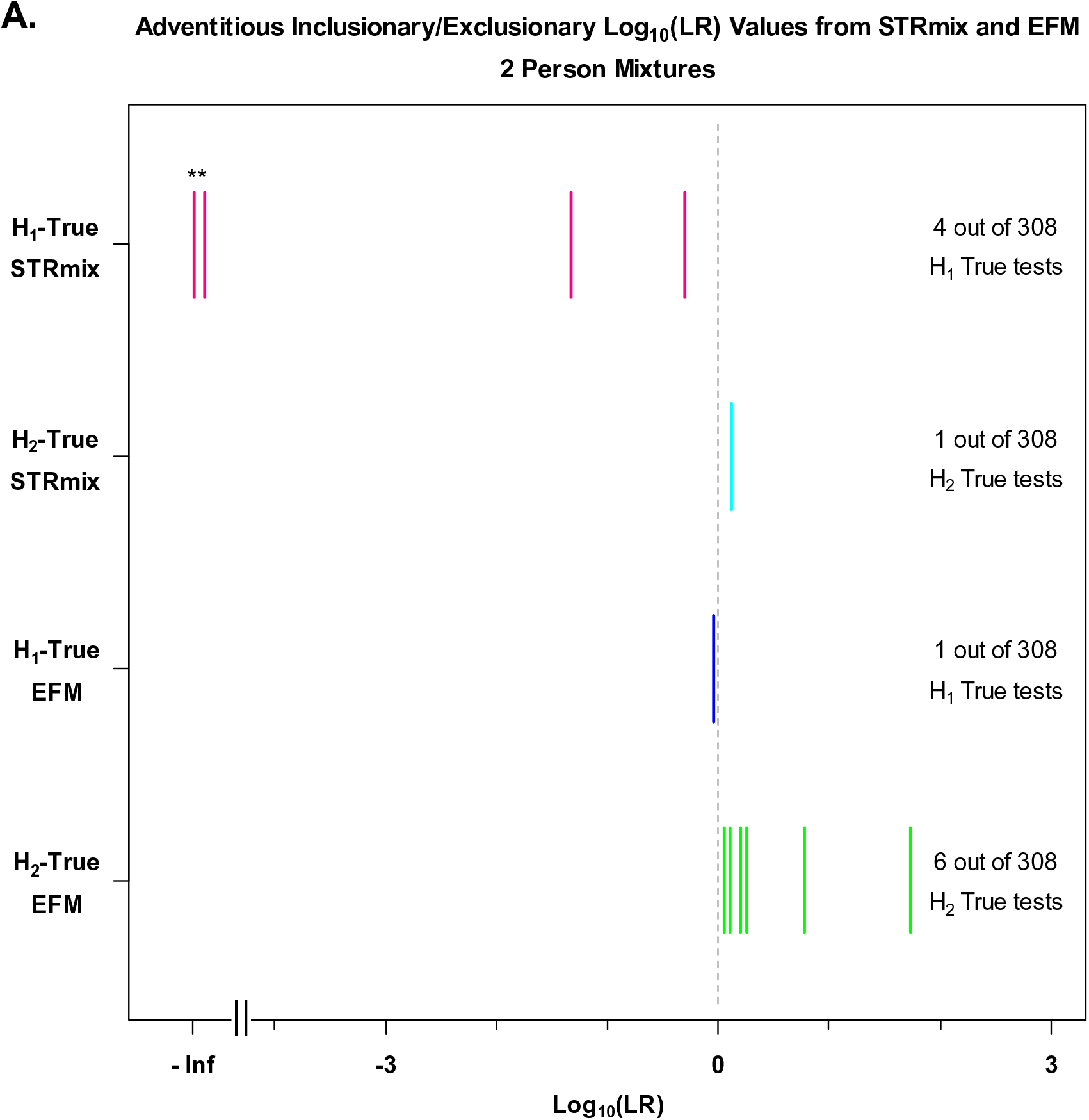
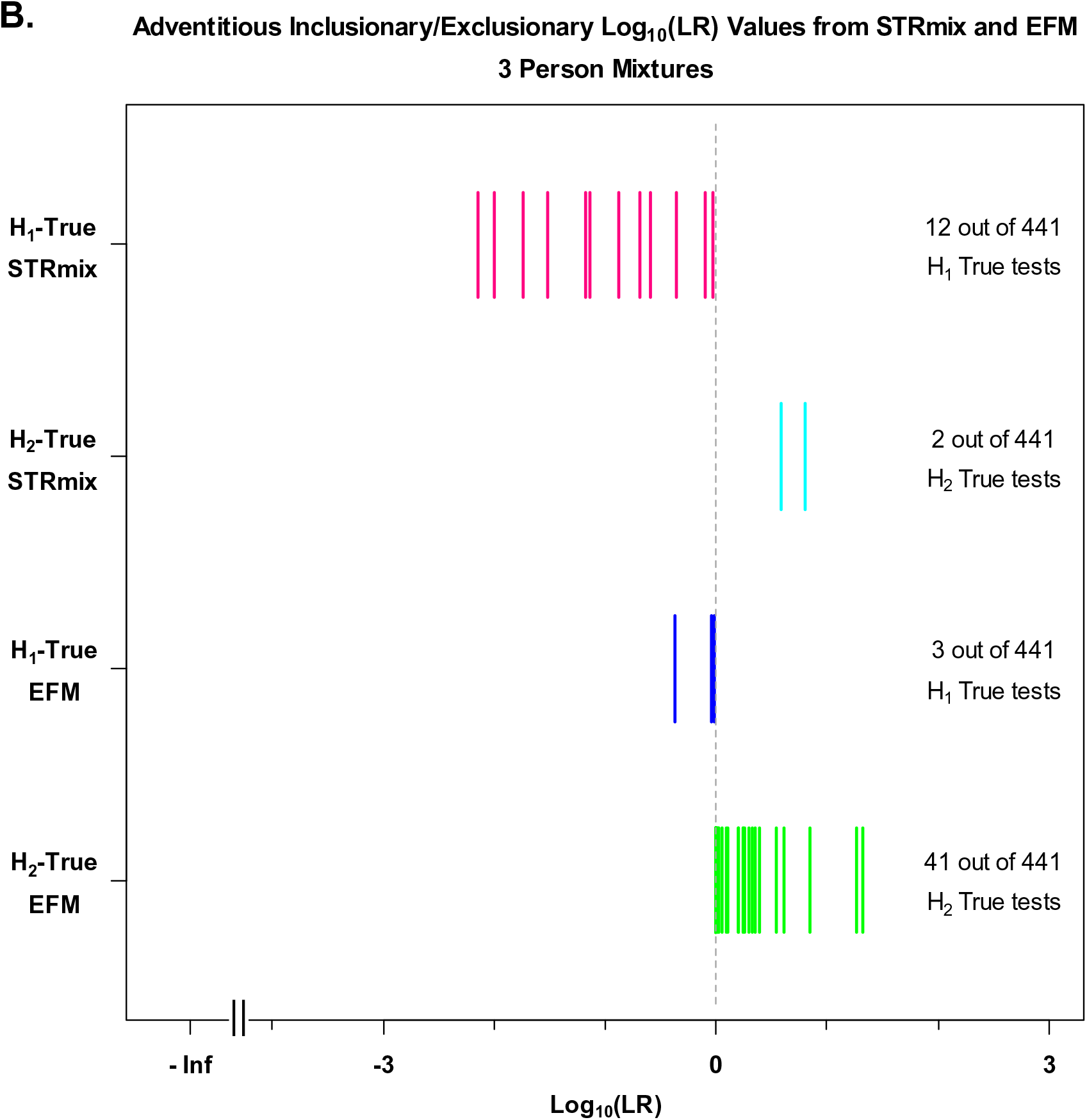
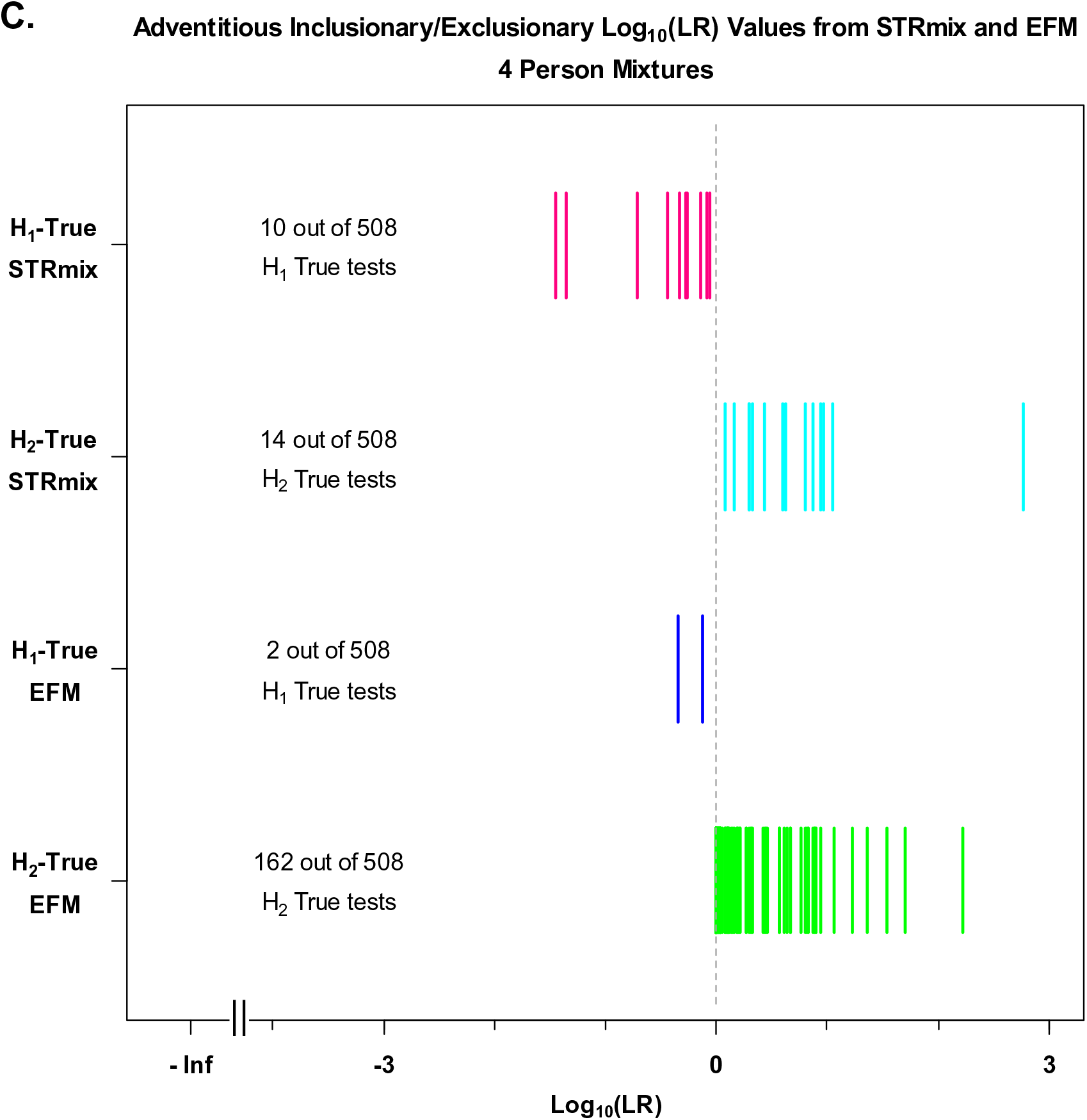
Summary of adventitious exclusionary and inclusionary support from both LR systems. Summary of the number of H1-true tests with their corresponding log_10_(LR) values from STRmix (magenta bars) and EFM (blue bars) that yielded adventitious exclusionary LRs (H2-support) and the number of H2-true tests with their corresponding log_10_(LR) values from STRmix (cyan bars) and EFM (green bars) that yielded adventitious inclusionary LRs (H1-support) for (A) 2P mixture profiles; here ** are the two 2P H1-true test interpretations for which STRmix assigned profile LRs of 0 (plotted in magenta at – Infinity (-Inf) on the log_10_ scale and discussed in detail in Section 3.6), (B) 3P mixture profiles, and (C) 4P mixture profiles. The x-axis shows the log_10_(LR) values for these adventitious exclusionary and inclusionary cases. The y-axis shows the labels of the tested propositions (H1 and H2) from each software (STRmix and EFM).

**Table 7.** Summary of the number of observations and frequency (%) of (A) known contributor analyses (H1-true tests) and (B) known non-contributor analyses (H2-true tests) that yielded log_10_(LR) values > 0 (or LR > 1) and log_10_(LR) values < 0 (or LR <1), respectively, broken down by number of contributors and software (STRmix and EFM). N represents the total number of either H1-true tests or H2-true tests conducted for the different number of contributors.

| **H1-True Tests: LR > 1** |  |  |  |  |
| --- | --- | --- | --- | --- |
| # of contributors | **STRmix** |  | **EFM** |  |
|  | Counts | Frequency % | Counts | Frequency % |
| 2 (N = 308) | 304 | 98.70 | 307 | 99.68 |
| 3 (N = 441) | 429 | 97.28 | 438 | 99.32 |
| 4 (N = 508) | 498 | 98.03 | 506 | 99.61 |
| Total (N=1,257) | 1,231 | 97.93 | 1,251 | 99.52 |

| **H2-True Tests: LR < 1** |  |  |  |  |
| --- | --- | --- | --- | --- |
| # of contributors | **STRmix** |  | **EFM** |  |
|  | Counts | Frequency % | Counts | Frequency % |
| 2 (N = 308) | 307 | 99.68 | 302 | 98.05 |
| 3 (N = 441) | 439 | 99.55 | 400 | 90.70 |
| 4 (N = 508) | 494 | 97.24 | 346 | 68.11 |
| Total (N=1,257) | 1,240 | 98.65 | 1,048 | 83.37 |

The results of the diagnostic checks examined here to check the performance of LR systems were as expected and in-line with previously described behavior of LR [8, 9, 51, 54, 57, 66, 68, 92, 93]. Visual comparisons of the global aggregate of log_10_(LRs) in the distribution plot of Fig.3 indicate qualitatively that STRmix and EFM seem to have equal ability in discriminating between H1-true and H2-true scenarios. Both LR systems indicate better discrimination performance for lower complexity mixtures than for higher complexity mixtures (mixtures characterized by an increase in NOC and/or decrease in DNA quantity and quality). These qualitative observations are substantiated statistically in Section 3.3.

### 3.2. Overall specificity and sensitivity of the two LR systems

An LR > 1 or a positive log_10_(LR) indicates that the evidence supports H1 over H2. An LR < 1 or negative log_10_(LR) indicates that the evidence supports H2 over H1. An LR = 1 doesn’t support one proposition over another; i.e., provides equal support for both propositions and represents neutrality (neutral evidence) [12, 57, 94].

In this section we discuss overall specificity and sensitivity (Table 7) and instances of adventitious exclusionary LRs of which H1-true tests resulted in LR < 1 and cases of adventitious inclusionary LRs of which H2-true tests yielded LR > 1 across both NOC and LR systems (Fig. 4 and Supplementary Tables 6 and 7).

Across all the 2P, 3P, and 4P mixtures, 97.93 % and 99.52 % of H1-true test LRs assigned by STRmix and EFM, respectively, were greater than 1 (or log_10_(LR) > 0) (Table 7A) while 98.65 % and 83.37 % of H2-true test LRs assigned in STRmix and EFM, respectively, resulted in LRs lower than 1 (or log_10_(LR) < 0) (Table 7B). The number of observations and frequency values are broken down by NOC and LR systems as shown in Table 7.

### Adventitious exclusionary support (examples of LR < 1 when H1 is true)

There were instances of adventitious exclusionary LRs for true contributor analyses (H1-true tests) within both LR systems that returned log_10_(LRs) < 0 as illustrated in Fig. 4 and Supplementary Table 6. Across the 1,257 H1-true tests conducted, there were 26 instances of adventitious exclusionary support with STRmix (4 out of 308 with 2P profiles, 12 out of 441 with 3P profiles, and 10 out of 508 with 4P profiles) and 6 instances with EFM (1 out of 308 with 2P profiles, 3 out of 441 with 3P profiles, and 2 out of 508 with 4P profiles) of which log_10_(LR) values for the POI were below 0. These are shown with their corresponding log_10_(LRs) in Fig. 4 and Supplementary Table 6. As expected from the behavior of the LR [57] and as shown in Supplementary Table 6, all the cases of H1-true tests with log_10_(LRs) < 0 from both LR systems mainly occurred when comparing the minor contributors to DNA mixture profiles that contained limited amount of information due to low minor template amount (e.g. ≤ 63 pg), low total template amount, compromised/degraded DNA, loci with allelic dropout, increase in the number of contributors, stochastic variation causing confounding information from the allelic and stutter peaks, and allele sharing between contributors [12].

The number of instances of H1-true tests with log_10_(LRs) < 0 was greater with STRmix than EFM. However, the log_10_(LRs) generated in EFM for these STRmix cases were mostly true inclusions of low-level LR range between (1-1,453) (i.e., uninformative or slightly to moderately supporting H1 over H2) with the exception of three 2P instances that are discussed in Section 3.4. For example, as seen in Supplementary Table 6, when mixture F10_RD14-0003-39_40-1;2-M3c-0.045GF was compared to the minor contributor “39”, STRmix gave a log_10_(LR) of -0.2 while EFM gave a log_10_(LR) of 0.9.

### Adventitious inclusionary support (examples of LR > 1 when H2 is true)

There were also instances of adventitious inclusionary LRs for known non-contributor analyses (H2-true tests) that returned log_10_(LRs) > 0 within both LR systems as illustrated in Fig. 4 and Supplementary Table 7. Out of the 1,257 total H2-true tests performed for 2P, 3P, and 4P, there were 17 log_10_(LRs) greater than zero analyzed with STRmix (1 out of 308 with 2P, 2 out of 441 with 3P profiles, and 14 out of 508 with 4P profiles) and 209 log_10_(LRs) greater than zero with EFM (6 out of 308 with 2P profiles, 41 out of 441 with 3P profiles, and 162 out of 508 with 4P profiles). These cases are presented with their corresponding log_10_(LRs) in Fig. 4 and Supplementary Table 7. The largest observed LR for the known non-contributors assigned by STRmix was 587 (log_10_(LR) = 2.7) and in EFM was 167 (log_10_(LR) = 2.2), when comparing a known non-contributor with the 4P mixture D02_RD14-0003-40_41_42_43- 1;1;1;1-M2e-0.124GF.

As expected, positive log_10_(LRs) obtained from non-donors in both software were attributed to one or more of the following: increased complexity of mixtures, increase in the number of contributors, mixtures generated from low total template and/or compromised low quality DNA, stochastic effects, and chances of allele sharing between the non-contributor profiles and evidence profiles [8, 57, 93, 95].

The number of instances of positive log_10_(LRs) from non-contributors were greater with EFM than STRmix (Fig. 4 and Supplementary Table 7). The LR values assigned by EFM were based on the MLE method, an approach that has elevated rates of LR > 1 for the H2-true tests than the conservative method as stated and observed in [66, 70]. However, these adventitious inclusionary LRs were low-level with range of values between 1 to 53 (i.e., uninformative or slightly supporting H1 over H2) (Supplementary Table 7) [54, 66, 70].

### 3.3. Using empirical Receiver Operating Characteristic (ROC) plots to study discrimination performance of the LR systems

We used Empirical Receiver Operating Characteristic (ROC) plots [89] as statistical tools to quantify the discrimination performance between the H1-true scenarios and H2-true scenarios of the two different LR systems. The discrimination performances were quantified using a numerical metric, the Area Under ROC Curve (AUC). AUC is the area between each ROC plot and the horizonal x-axis (Fig 5). Statistical tests (p-values) for AUC comparisons (i.e., differences between the ROC plots) were calculated and listed in Fig. 5 [96].

**Fig. 5.**
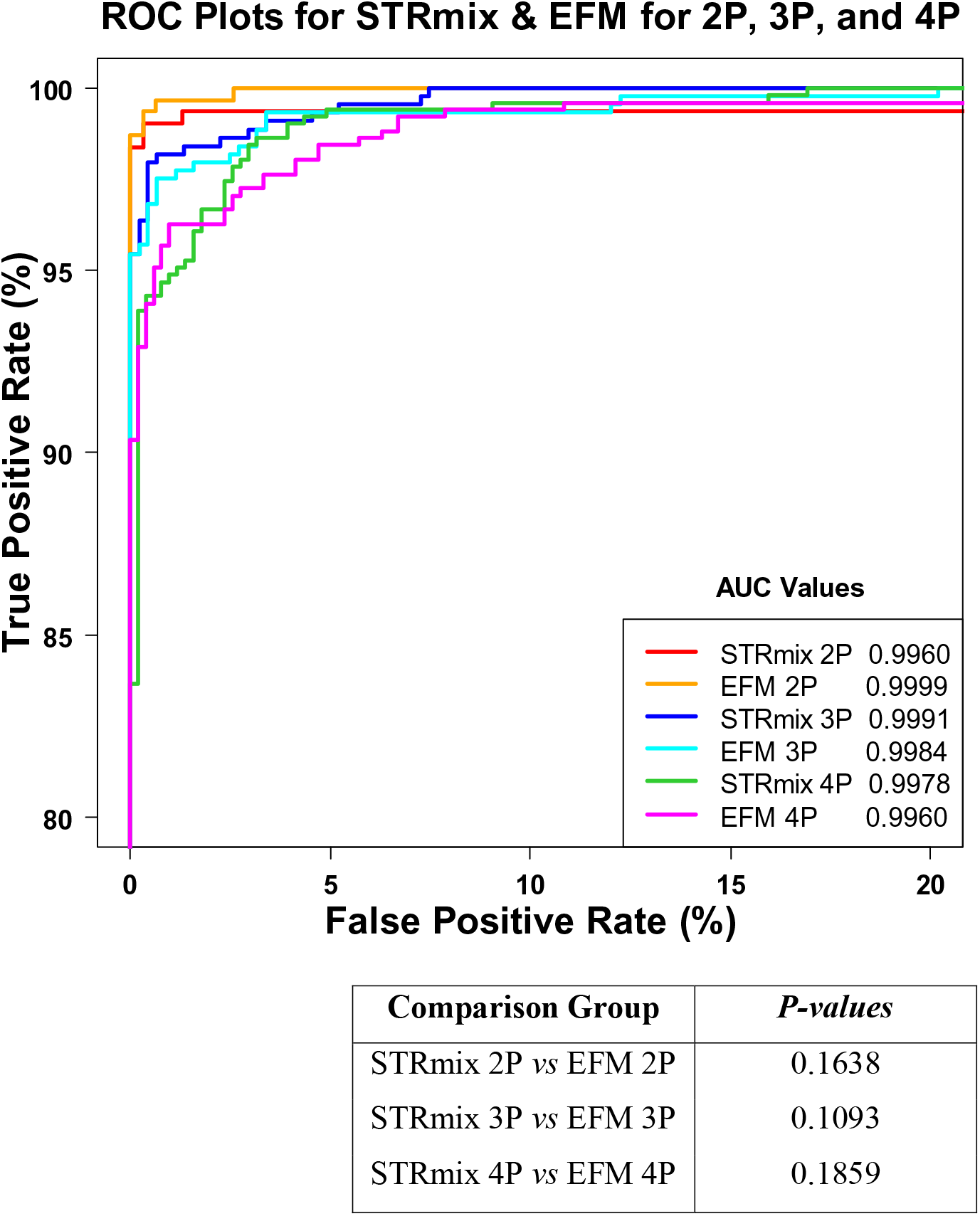
Using empirical ROC plots to study discrimination performance of the LR systems. ROC plots are built per varying NOC and software. Each NOC dataset is composed of profiles of different DNA quality, quantity, and mixture proportions. The red, blue, and green curves are the ROC plots constructed using LR values of known contributors and known non-contributors of 2P, 3P, and 4P mixtures analyzed within STRmix, respectively. ROC plots constructed with LR values assigned by EFM are shown in orange (2P), cyan (3P), and magenta (4P). The plot contains a total of 308 H1- true tests and 308 H2-true tests for the 2P analysis, 441 H1-true and 441 H2-true calculations for the 3P analysis, and 508 H1-true and 508 H2-true tests for the 4P mixtures. Discrimination performances were quantified using the area under the ROC curve (AUC). Statistical tests (p-values) were calculated for the AUCs and are shown in the table. P-values > 0.05 indicate that the differences between the two software in their ability to discriminate between H1-true and H2-true scenarios were found to be statistically indistinguishable.

As detailed in Section 2.11, LR values of the H1-true tests and H2-true tests were combined across each NOC level (2P, 3P, and 4P) generated from each software (STRmix and EFM), thus creating six datasets: STRmix 2P, EFM 2P, STRmix 3P, EFM 3P, STRmix 4P, and EFM 4P. A series of various LR thresholds were applied on each dataset generating true positive rates (TPR) and the corresponding false positive rates (FPR). ROC plots were created by plotting the TPR (along vertical axis) versus the FPR (along horizontal axis). ROC plots constructed using the H1-true test LRs and H2-true test LRs assigned by STRmix are shown in red (2P mixtures), blue (3P mixtures), and green (4P mixtures) (Fig. 5). ROC plots constructed with LR values assigned by EFM are shown in orange (2P mixtures), cyan (3P mixtures), and magenta (4P mixtures). The AUC values quantified for each ROC plot are also shown in Fig. 5.

The p-values of the comparisons of areas under the ROC plots of: STRmix 2P vs EFM 2P, STRmix 3P vs EFM 3P, and STRmix 4P vs EFM 4P were > 0.05 (Fig. 5), indicating that for the considered data the differences between the two software in the ability to discriminate between H1-true and H2-true scenarios were not statistically significant.

The ROC plots shown in Fig. 5 statistically support the qualitative observation visualized in the distribution plots of Fig. 3. Therefore, the ability for the two LR systems to discriminate between known contributors and known non-contributors are statistically indistinguishable for the data that we considered. However, that does not imply that STRmix and EFM are producing equal LR values or agreeing when the same profile is being interpreted within both software. Sample to sample comparisons are discussed in Section 3.4. Rather the plots in Figs. 3 and 5 are considering the data in aggregate.

### 3.4. Global overall profile log_10_(LR) values of H1-true tests and H2-true tests from each LR system

As discussed in Sections 3.1 and 3.3 and considering the data used herein, the discrimination power of both LR systems were shown to be not statistically distinguishable using H1-true and H2-true LR distributions (Fig. 3) and ROC plots (Fig. 5). However, this did not imply that both LR systems assigned the same LR values on a case-by-case basis. In this section, we discuss to what extent actual LR values for the 2P, 3P, and 4P mixture profiles varied across the two LR systems.

Scatter plots (Fig. 6) were produced by plotting the log_10_(LRs) of the H1-true tests (magenta datapoints) and the H2-true tests (blue datapoints) obtained from STRmix on the x-axis against the corresponding log_10_(LRs) assigned using EFM on the y-axis for the 2P (Fig. 6A), 3P (Fig. 6B), and 4P (Fig. 6C) mixture profiles. Identical or near identical log_10_(LR) values assigned by both LR systems fell on the solid black 45^ο^ degree line, X=Y. Datapoints that did not fall on the diagonal line corresponded to instances with varying degrees of difference in the overall LR profile between the two LR systems. For example, datapoints located within the two black dashed lines, two black dash-dotted lines, and two black dotted lines surrounding the line X=Y, corresponded to cases with LR results differing by a factor as high as 10^2^, 10^4^, and 10^6^, respectively (Fig. 6). Datapoints that are outside the pair of black dotted bands represented LRs assigned by the two software that differed by more than a factor of 10^6^. These differences represented instances where either the LRs obtained from STRmix exceeded the ones obtained in EFM or vice versa. It is interesting to note that differences in the assigned LR values were greater with the non-contributor testing profiles than with the H1-true testing cases. Instances that differed by factor of ≥ 10^3^ and the potential explanations for the differences will be discussed in Section 3.6. Impacts of the differences in the inter-software numerical LRs on verbal expression will be discussed in Section 3.7.

**Fig. 6.**
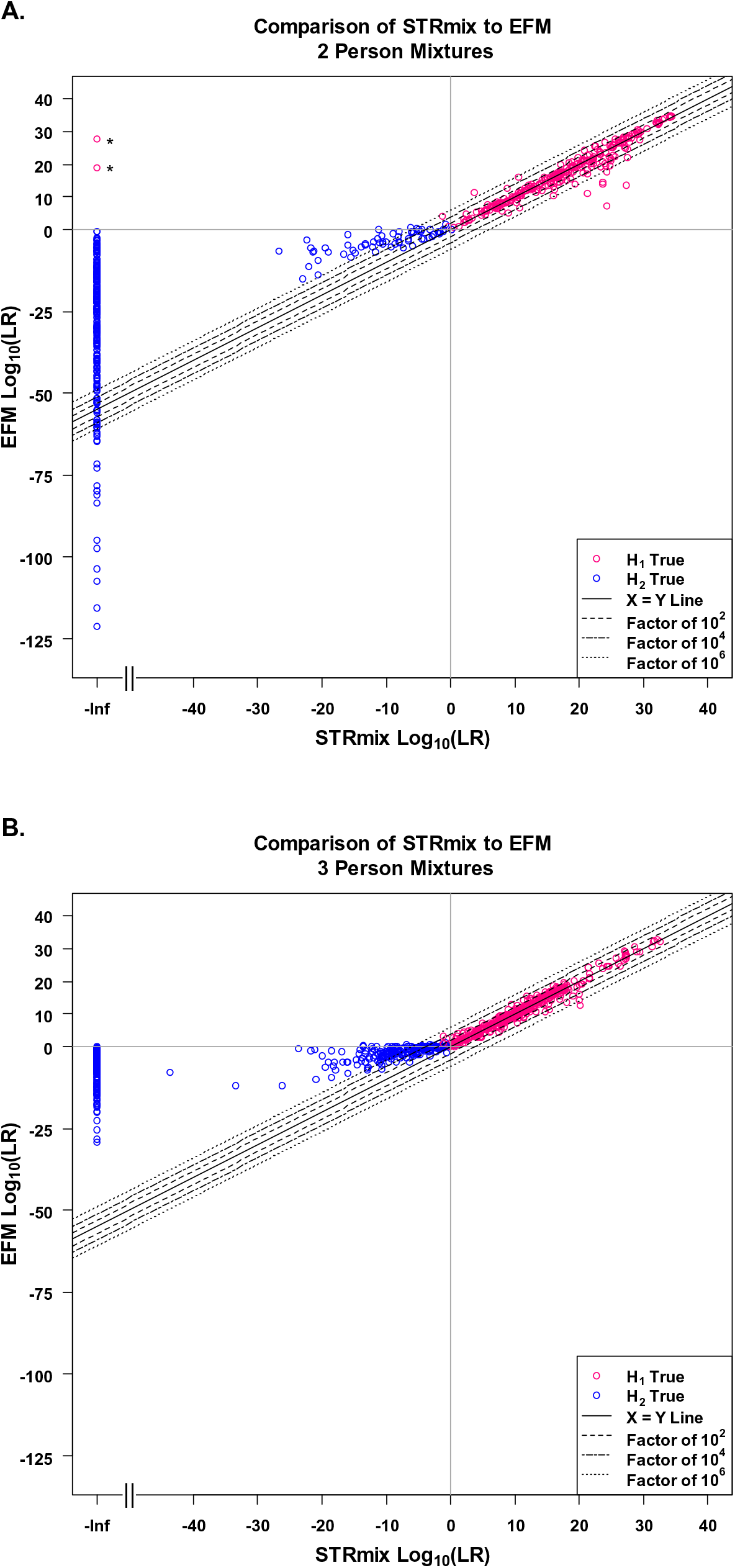
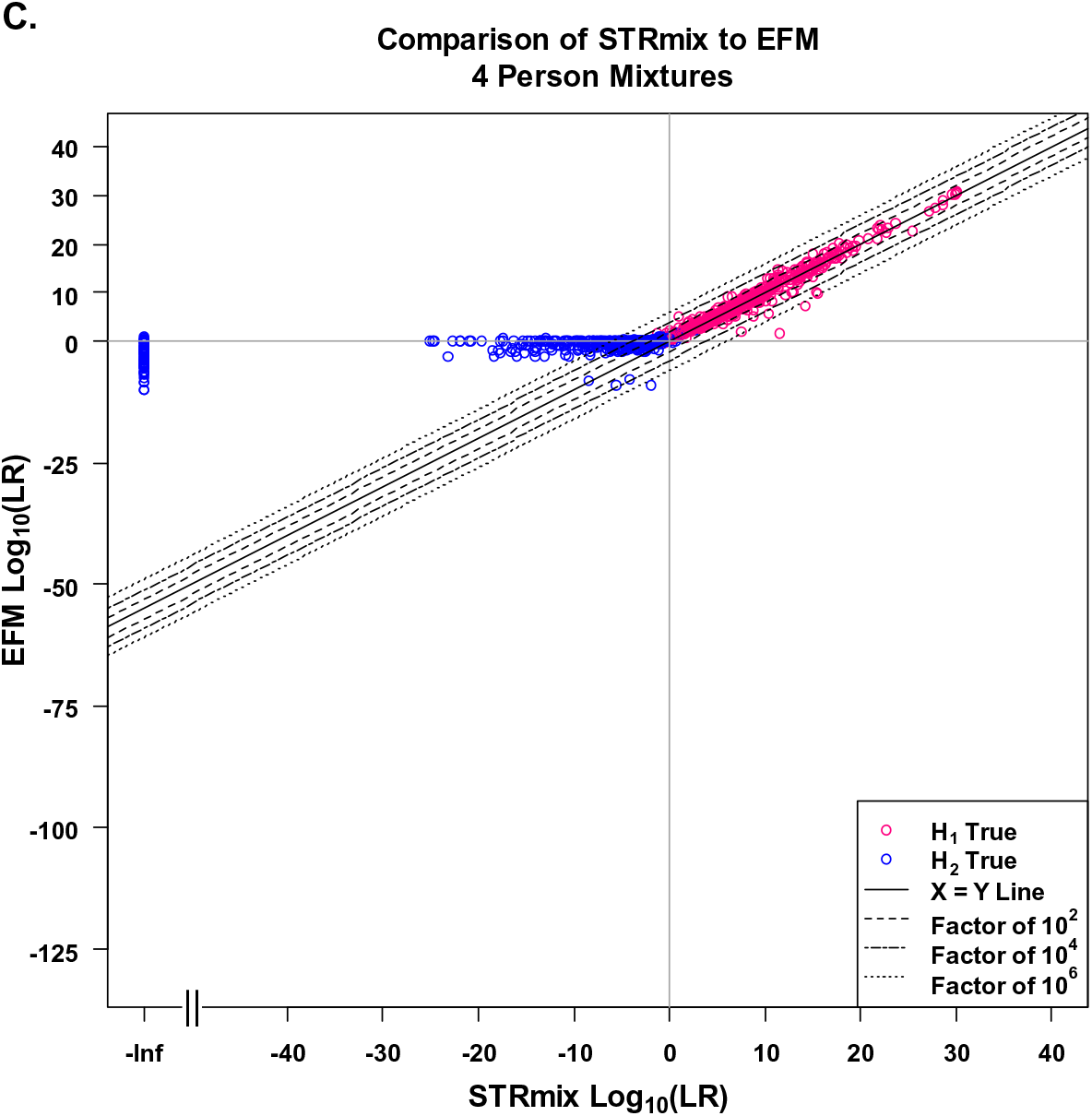
Global overall profile H1-true test and H2-true test log_10_(LR) values assigned by STRmix and EFM. Scatterplots describing the comparisons of the log_10_(LR) values for true contributors and known non-contributors assigned by STRmix (plotted on x-axis) to those assigned by EFM (plotted on y-axis) using (A) 2P, ** are the two 2P H1-true test interpretations for which STRmix assigned profile LRs of 0 (plotted in magenta at – Infinity (-Inf) on the log_10_ scale and discussed in detail in Section 3.6), (B) 3P, and (C) 4P mixture profiles. Log_10_(LRs) of true known contributors are plotted in magenta and log_10_(LRs) from known non-contributors are plotted in blue. The solid black diagonal line represents X=Y. Points that fall on X=Y indicate that the comparison of log_10_(LRs) from both software were similar. Datapoints located within two black dashed lines, two black dash-dotted lines, and two black dotted lines corresponded to calculated LRs that were different between the two software by a factor of 10^2^, 10^4^, and 10^6^, respectively.

To conclude this section, although both LR systems show comparable discrimination performance, differences exist in log_10_(LR) values on a case-by-case basis and that is expected [26]. Differences in log_10_(LR) values assigned by STRmix and EFM at the profile level covered a wide range from zero to over a million (discussed in detail in Sections 3.5 and 3.6) for the same input data (i.e., the same EPG). The differences appear to be greater in the H2-true cases than in the H1-true cases.

### 3.5. Distribution of differences in log_10_(LR) values between the two LR systems

Here, we describe and plot the degree and distribution of the observed differences between the two LR systems. The actual differences in log_10_(LRs) were calculated in both directions (i.e., log_10_(LR)_STRmix_ – log_10_(LR)_EFM_ as well as log_10_(LR)_EFM_ – log_10_(LR)_STRmix_) for the H1-true tests and H2-true tests (histograms shown in Fig. 7). These differences were broken down into factor of 10 bins for the 2P (Fig. 7A), 3P (Fig. 7B), and 4P (Fig. 7C) analysis and the relative frequencies (in %) of these differences are indicated for each bin in Fig. 7. For example in Fig 7A, 21.4% and 47.1% of the differences for the 2P H1-true tests were between 0 to 1 on log_10_ scale for log_10_(LR)_STRmix_ - log_10_(LR)_EFM_ (black histograms) and log_10_(LR)_EFM_ - log_10_(LR)_STRmix_ (grey histograms), respectively. The relative frequency histograms (Fig. 7) indicate that (i) the differences between the two LR systems were smaller with the H1-true testing cases than with the non-contributor tests and (ii) EFM tended to give higher LR values than STRmix for both the H1-true tests and H2-true tests.

**Fig. 7.**
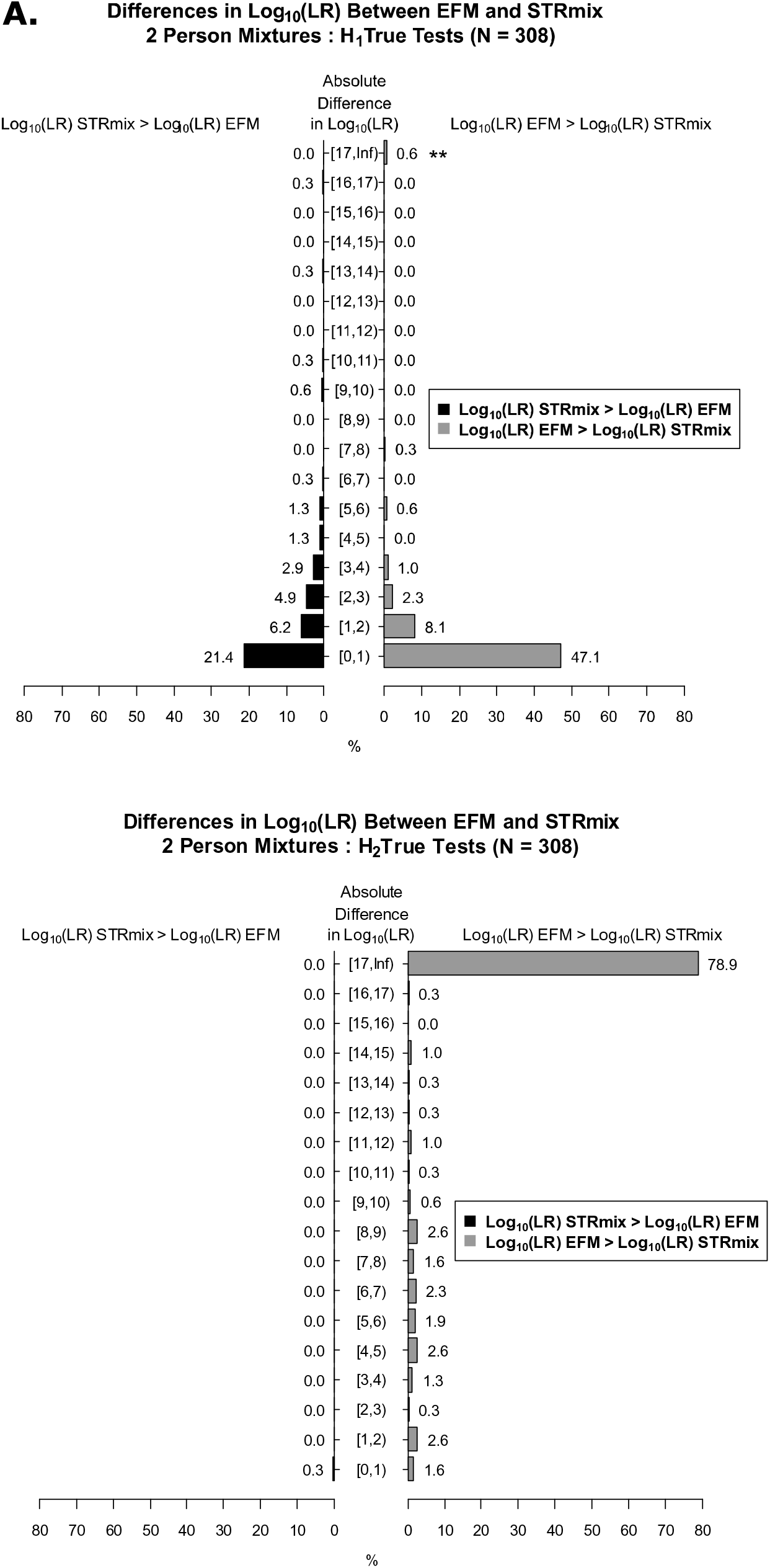
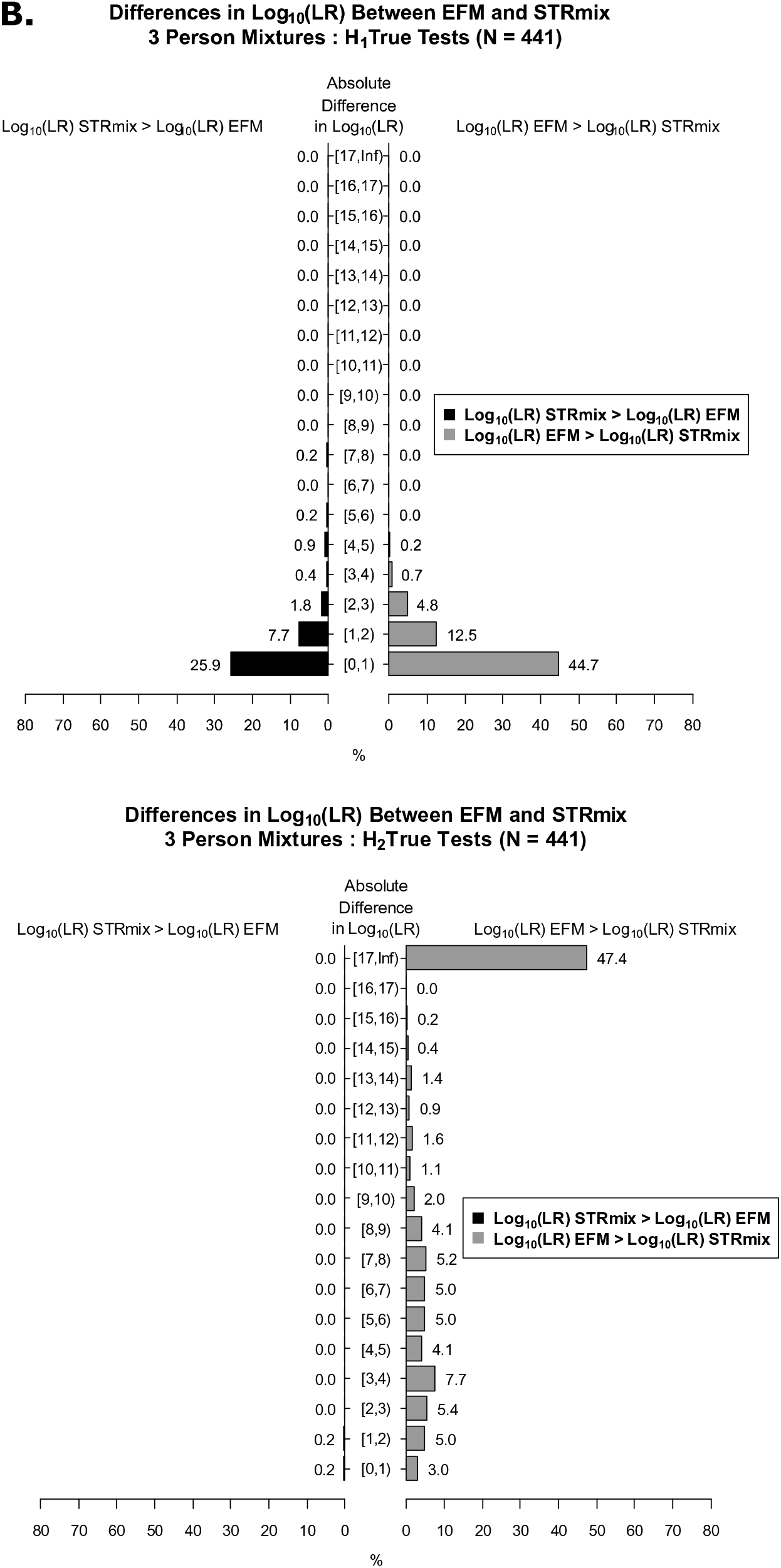
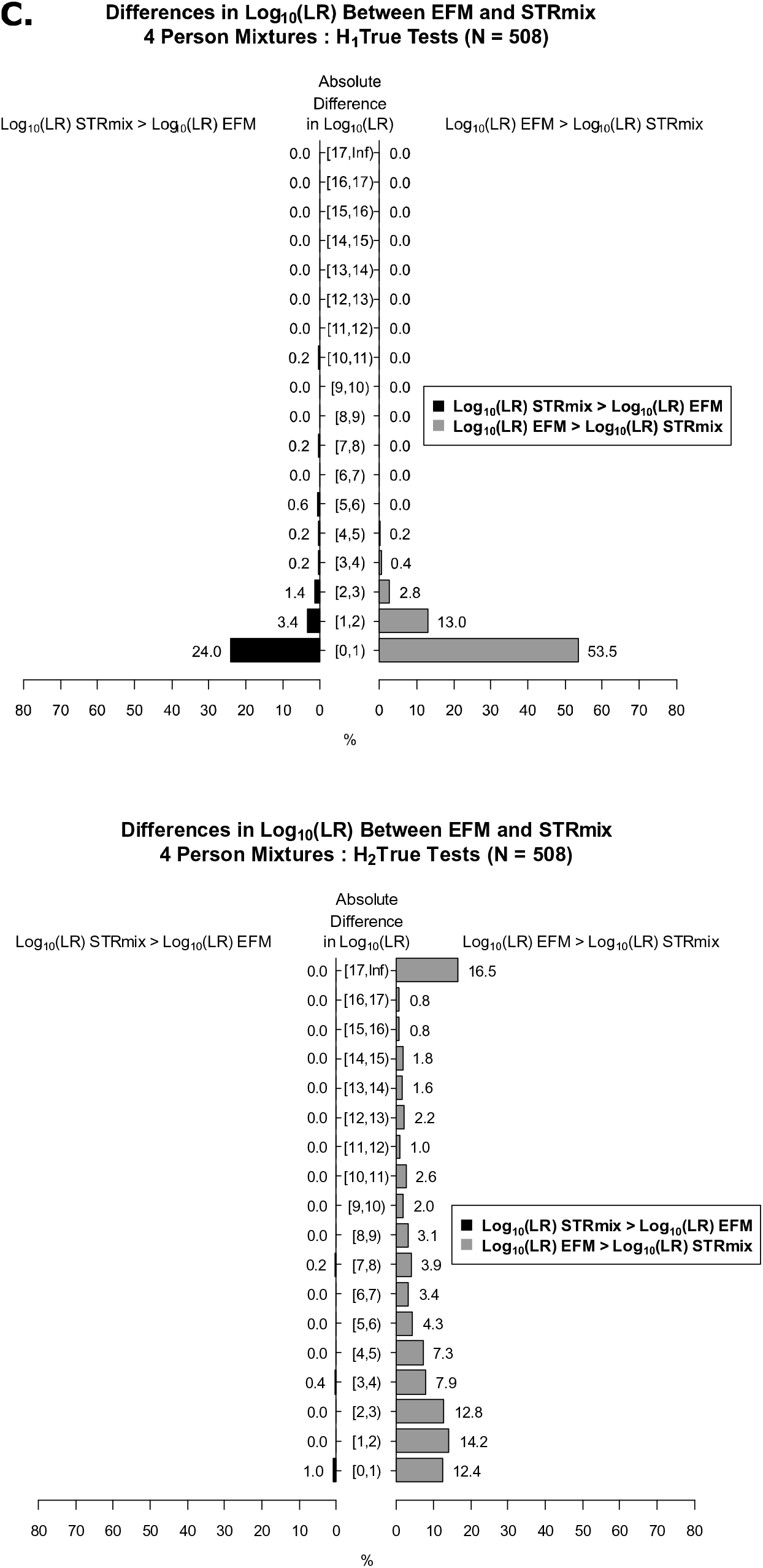
Relative frequency histograms of the degree of differences in log_10_(LR) values between the two LR systems. The histograms illustrating the differences in log_10_(LR) values between the two LR systems that were broken down into factor of 10 bins (shown on the y-axis as absolute difference in log_10_(LR)) for: (A) 308 2P H1-true tests and 308 2P H2-true tests where ** are the two 2P H1-true test interpretations for which STRmix assigned profile LRs of 0 (binned into the [17, Inf) category and discussed in detail in Section 3.6), (B) 441 3P H1-true tests and 441 3P H2-true tests, and (C) 508 4P H1-true tests and 508 4P H2-true tests. The square bracket “[“ in the interval notation ”[)” indicates that the endpoint is included in the interval and the parenthesis “)“ in the interval notation ”[)” indicates that the endpoint is not included. For example, [1,2), is the interval of values between 1 and 2, including 1 and up to but not including 2, i.e., 1 ≤ values < 2. The x-axis shows the relative frequencies (in %) of the differences in log_10_(LR) values between the LR systems occurring within each bin. The relative frequencies are also labeled above each bar of the histogram. The black histograms represent the analysis from log_10_(LR)_STRmix_ - log_10_(LR)_EFM_. The grey histograms represent the analysis from log_10_(LR)_EFM_ - log_10_(LR)_STRmix_.

The actual differences in log_10_(LRs) for the H1-true tests were further stratified by the type of POI (i.e., major, minor, and equal contributors as defined in Supplementary Table 4) constituting the 2P (Fig. 8A), 3P (Fig. 8B), and 4P (Fig. 8C) mixture profiles. As shown from the distribution plots in Fig. 8, the magnitude of the differences for the two LR systems were greater for the minor contributors (shown in magenta) than for the major (shown in blue) and for the equal (shown in green) contributors. LRs assigned by STRmix and EFM agreed more when POI(s) constitute the equal contributors of the mixture (Fig. 8). This is expected because with balanced profiles, peak height information content has less effect on LR calculations than in cases of major: minor profiles [9, 65, 68–70].

**Fig. 8.**
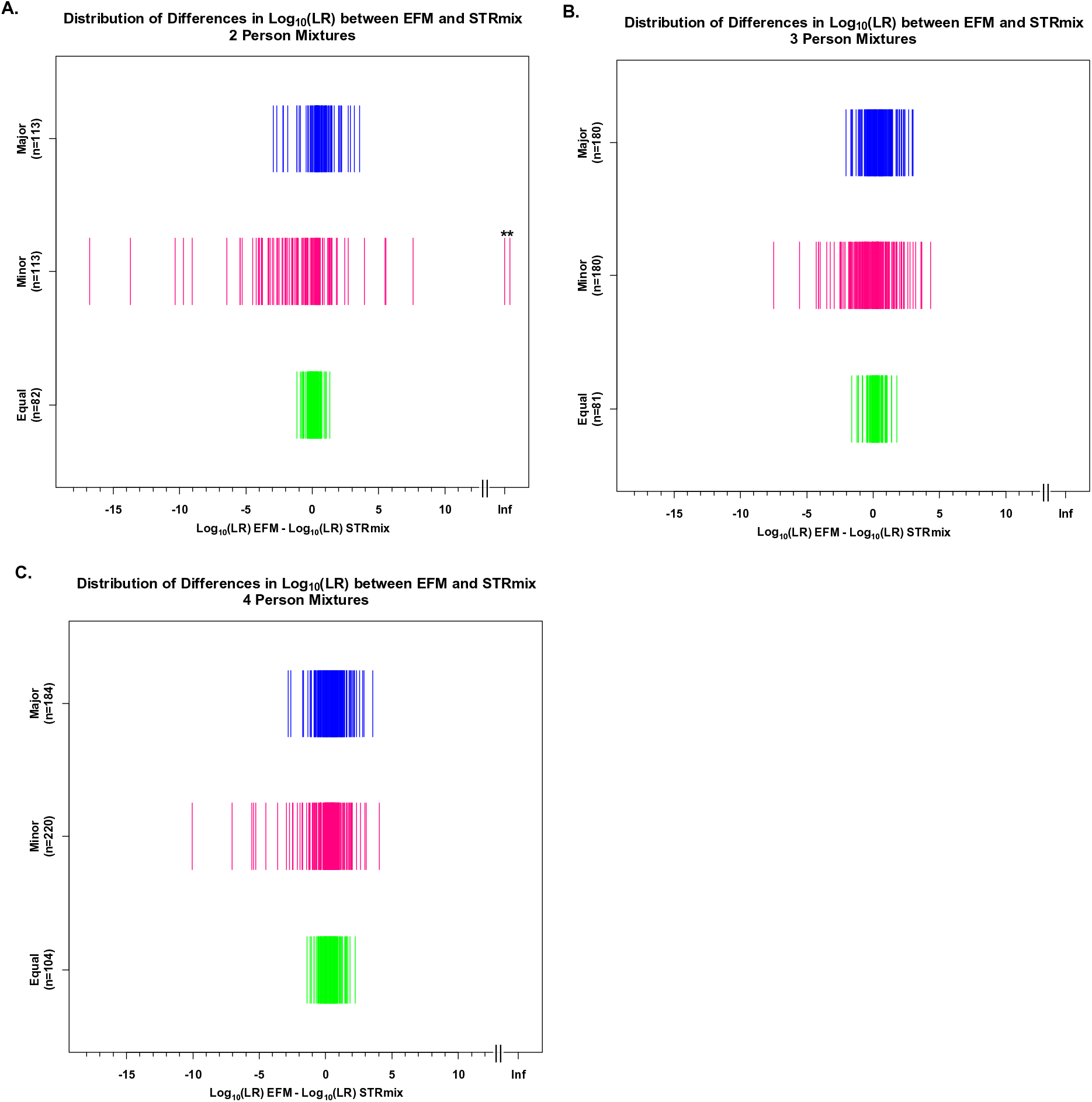
Distribution of differences in log_10_(LRs) across major, minor, and equal contributors. Distribution plots describing the differences in log_10_(LRs) here shown between EFM and STRmix (log_10_(LR)_EFM_ – log_10_(LR)_STRmix_) for the (A) 2P where ** are the two 2P H1-true test interpretations for which STRmix assigned profile LRs of 0 (plotted in magenta at Infinity (Inf) on the log_10_ scale and discussed in detail in Section 3.6), (B) 3P, and (C) 4P analyses of the H1-true tests when POI constitutes the major (shown in blue), minor (shown in magenta), and equal (shown in green) contributors of the mixtures. The differences in log_10_(LRs) are plotted on the x-axis in log_10_ scale. The y-axis shows the labels of the types of POI with their corresponding number of observations.

### 3.6. Evaluation of apparent differences in log_10_(LR) values between the two LR systems

In this section we discuss the steps performed to further investigate differences in the assigned LR values obtained from the two LR systems on a case-by-case basis, where the differences are observed, and the potential explanations for these differences. We restrict our discussion to instances when LR (STRmix) 1000*LR (EFM) that constituted 7.3% of the 2P, 1.7% of the 3P, and 1.4% of the 4P H1-true tests (histograms of Fig. 7 and Supplementary Table 11) and instances when an LR (EFM) ≥ 1000*LR (STRmix) that accounted for 2.5% of the 2P, 0.9% of the 3P, and 0.6% of the 4P H1-true tests (histograms of Fig. 7 and Supplementary Table 12). Only differences in H1-true results (true known contributor samples) are discussed.

LR computations obtained from the two software were based on same/fixed EPG features, same pair of propositions, NOC, theta, and population allele frequency. Therefore, results presented here shows that differences observed in LR values can occur due to one or more of the following reasons:

I. Nonconvergence of the Markov Chain Monte Carlo (MCMC) algorithms and maximum likelihood estimation (MLE)
II. Decision to provide identical EPGs for both LR systems
III. Different modeling assumptions and parameters settings between the two software

We discuss each of the above reasons and provide examples from the data set. The availability of both the mixture and reference profiles was beneficial and helped in the investigation of observed differences of the assigned LR values.

#### I. Non-convergence of the MCMC algorithms and MLE

STRmix pdf reports contain summary statistics for each interpretation conducted in the software and can be used by analysts as diagnostics on the performance of the interpretation according to the specified models. These diagnostics have been classified into primary and secondary categories and are discussed in detail in Russell et al. [87]. In actual casework, every analysis should be subjected to diagnostic checks. But in this study and for practical reasons only cases where STRmix and EFM differed by a factor of ≥ 10^3^ were inspected for genotypic weights, mixture proportions, per-locus LRs, log(likelihood), peak height variance parameters, and Gelman-Rubin (GR) statistics.

Two extreme differences observed between STRmix and EFM were with the 2P mixture profiles, C02_RD14-0003-40_41-1;4-M2U15-0.315GF (herein referred to as “C02”) and H06_RD14-0003-48_49-1;4-M2e-0.315GF (referred to as “H06”) (Supplementary Table 8). C02 and H06 generated profile LR of 0 in STRmix when compared to true known minor contributors, 40 and 48, respectively. A locus LR value of 0 will lead to a profile LR of 0. The log_10_(LR) assessments for these profiles in EFM were 27.6 for C02 and 19.0 for H06. Unlike STRmix, EFM displays low to very low LRs for exclusionary loci but does not provide a zero locus LR. A review of the per locus LRs (Supplementary Table 8) assigned to the evaluation of the POIs in STRmix indicated that almost all loci favor inclusion (LR > 1) except for a single locus displaying an LR of 0 in each interpretation, D1S1656 in C02 and D3S1358 in H06. Instances of single locus LR = 0 have been observed using different data from different studies [8, 51, 55]. In such cases, options are to either ignore that locus during deconvolution, or repeat the deconvolution in STRmix with: a random starting seed for the MCMC different than the one that gave LR = 0, or an increase in number of MCMC accepts, or a larger Random Walk Standard Deviation (RWSD) [8, 12, 51, 55, 87]. Here, we repeated the runs in STRmix with more MCMC accepts (as discussed in Section 2.10) and the repeated interpretations generated non-zero LRs for the affected loci, and profile log_10_(LRs) of 24.8 and 19.6 (Supplementary Table 8). It is to note that these two discussed 2P H1-true test interpretations with profile LRs of 0 assigned by STRmix were plotted: (i) at −125 on the log_10_ scale in Fig. 3 and Additional Files 1 and 2; (ii) at – Infinity (-Inf) in Figs. 4A and 6A; (iii) at Infinity (Inf) in Figs. 7A and 8A; and were binned into the exclusionary verbal category (Table 9A).

Another extreme difference observed between STRmix and EFM was with E04_RD14-0003- 42_43-1;9-M2U105-0.15GF, a 2P mixture profile of which comparison to the minor contributor in STRmix and EFM, yielded log_10_(LRs) of -1.3 and 4.1, respectively (EPG and data shown in Supplementary Table 9). A review of the STRmix output indicated negative log(likelihood), which might be due to several reasons including “flawed input data” [8, 87]. Inspection of the DNA typing results, ground truth genotypes of the POIs, and deconvolution results indicated retained artifact peaks binned into alleles at two loci, D19S433 “18.2” and D5S818 “14” (EPG shown in Supplementary Table 9). The artifacts were each modelled in STRmix as being allelic in origin and were included in the genotypic combinations thus leading to exclusion after comparing the resolved profile to the true contributors [12, 51]. In such cases, mixture samples can be re-injected or reamplified [8]. However, since only the electronic data was accessible for this study, the artifacts were removed and the input file were re- interpreted in both STRmix and EFM, generating profile log_10_(LRs) of 0.8 and 4.6, respectively (Supplementary Table 9).

A GR > 1.2 might be an indication that more MCMC runs may be needed for convergence [12, 87, 97]. Profiles indicating discrepancies of ≥ 10^3^ and with a GR > 1.2 (a total of 6 out of 53) were reinterpreted in STRmix using higher number of burn-in and post burn-in accepts [12]. The repeated LR computations resulted in lower GR. Although the GR decreased, there was either no effect or a slight increase by a factor of 10 in the profile overall LRs (Supplementary Table 10) and did not substantially alter the observed LR differences in these cases.

EFM provides an option for selection of one of four models (turning on either or both of the degradation and stutter models) under H1 and H2 hypothesis and generates a Probability-Probability (PP) plot to examine if the model selected explain the observed data adequately [24, 54, 98]. A linear trend of PP plots within 99% Bonferroni band indicates that the assumed continuous models may be adequate for the data of the observed peak heights above the detection threshold [24, 98]. Herein, we selected the model with both degradation and back-stutter options turned on and cross checked a total of four mixture profiles out of 53 interpretations showing discrepancies (Additional File 3). The PP plots showed that models selected (i.e., degradation ON and stutter ON) appear to adequately explain the data.

These observations indicate that non-convergence of MCMC and the inability of the software to describe the observed profile given the provided information are one of the reasons behind the observed differences.

#### II. Decision to provide identical EPGs for both LR systems

Instances of the underestimation of LR values observed in EFM as compared to STRmix was primarily due to the unmodelled stutter type peaks not filtered from the input files (Supplementary Table 11). Stutter models for B2, B1, and F1 were applied to the mixture deconvolutions performed in STRmix. EFM v2.1.0 used in this work only models stutter peak heights in the -1 repeat unit position [24, 66, 98]. The B2 and F1 peaks retained after applying the analytical thresholds and not pre-filtered from the DNA profiles before analysis in EFMv2.1.0 led to instances of smaller LRs than those assigned by STRmix (Supplementary Table 11). As reflected in (Supplementary Table 11), differences were highest with minor contributors in major/minor mixture profiles where allele peak heights from a minor contributor can have the same size and height as stutter peaks of major contributors [66, 87].

To further examine this hypothesis, we removed the retained (i.e., above AT) unmodelled F1 and B2 stutter peaks from the profiles that showed a difference of factor of ≥ 10^3^ and reinterpreted the analysis in EFM v2.1.0 for the 2P, 3P, and 4P mixtures. The LRs of the minor contributors in the repeated profiles increased substantially, thus decreasing the differences in log_10_(LR) values observed between STRmix and original EFM runs (Supplementary Table 11).

Our intentions of leaving in the unmodelled stutters (F1 and B2) were to have identical EPGs as input files for both software especially since according to certain publications any unmodelled stutter could be explained as drop-in allelic events [29, 66]. For example, according to You et al. [29], “All the alleles explained by the over-stutter (OS) or double-stutter (DS) models could also be explained by the drop-in model, and so it is unclear whether or not there is a material benefit from modelling DS and OS in addition to drop-in, an option that is available in likeLTD”. According to Bleka et al. on the effect of applying the drop-in model to accommodate an extra allele in [66]: “Hence we observe that the implemented drop-in model in EuroForMix accommodates spurious alleles very efficiently - there is a small decrease in the LR. As expected, the larger the peak height, the greater the reduction in LR, because it impacts on heterozygote balance with other alleles.” These unmodelled stutter peaks were considered in certain cases less likely to be drop-ins than alleles and therefore were considered alleles instead as observed from the profile LRs, per locus LRs, and deconvolution (data not shown). A new EFM version 3.2.0 [25, 99] is now available and accounts for forward stutter in LR calculations. This new version was not available during the time of the analysis.

Unmodelled stutter peaks (F1 and B2) can be removed before interpretation to improve the fit of the model to the observed data by using stutter-type specific thresholds [54]. However, there is no guarantee that the stutter thresholds will work all the time across all the cases due to false positives (stutter peaks are left in as alleles) and/or false negatives (removing low-level alleles of the minor contributors).

We discuss an illustration in Additional file 4 on one of the profiles shown in Supplementary Table 11. D05_RD14-0003-48_49-1;4-M3a-0.315GF is a two-person mixed GlobalFiler (GF) DNA profile with major and minor contributors from the PROVEDIt dataset with pristine DNA (a) of total DNA amount of 315pg and mixture ratio of 1:4. When the POI corresponded to the major contributor, STRmix and EFM gave near identical profile log_10_(LR) values of 27.6 and 27.9, respectively. However, for the minor contributor position, STRmix and EFM gave profile log_10_(LR) values of 27.4 and 21.9, respectively, leading to a 5.4 difference in log_10_ scale (Additional file 4A). A further review of the per-Locus LR tables obtained from STRmix and EFM for the minor contributor indicated that all loci had LR values favoring inclusion (i.e., LR > 1), except for the D22S1045 in EFM that has been assigned a locus LR < 1 (i.e., 0.001139) (Additional file 4B). A review of the mixture profile (Additional files 4C and 4D) indicated that the exclusionary LR at D22S1045 generated from EFM is likely due to a peak at “16” at D22S1045 which is likely an F1 of allele “15”. EFMv2.1.0 did not model F1 and had accounted for “16” as being allelic in origin instead of being modeled as “drop-in” (Additional file 4D). We removed the “16” from the input file and reinterpreted in EFMv2.1.0. The rerun gave a D22S1045 locus LR of 16.2 (Additional file 4E) and a profile log_10_(LR) of 26.1 (Additional file 4F), thus decreasing the discrepancy between EFM and STRmix to a factor of approximately 10.

There were cases (e.g. A03-40_41-1;4-M2U105-0.315GF; H03-48_49_50_29-1;4;4;4-M3I22- 0.75GF; E03-48_49_50_29-1;4;4;4-M2I15-0.75GF; D01-50_29_30_31-1;1;2;1-M2a-0.155GF), that did not contain any instances of F1 or B2 and differed by a factor of ≥ 10^3^ when compared to the profile LR generated in EFM (highlighted in red in Supplementary Table 11). A plausible explanation for these differences will be discussed below.

#### III. Different modeling assumptions and parameters settings between the two software

There were instances in which EFM assigned larger LR values than STRmix (Supplementary Table 12) and cases of which STRmix profile LRs were greater than EFM LRs (highlighted in red in Supplementary Table 11 and as mentioned above not due to F1 or B2). Some of these profiles in which EFM assigned larger LR values than STRmix contained instances of F1 and B2. Reinterpreting those profiles in EFMv2.1.0 with F1 and B2 removed resulted in a slight increase or had no effect on the profile LRs (Supplementary Table 12). Larger differences between the two LR systems were observed when comparing minor contributors (in most cases) with mixture profiles composed of low total template amount, low minor template amount, and/or degraded DNA (as reflected in Supplementary Table 12). In these cases, there is increase in stochastic effects, variation in peak heights, and drop-out events.

As an illustration we discuss one of the profiles shown in Supplementary Table 12. B07_RD14- 0003-48_49-1;4-M3e-0.075GF is a two-person mixed GlobalFiler (GF) DNA profile with major and minor contributors from the PROVEDIt dataset with degraded DNA (DNA treated with DNase I) of total DNA amount of 75 pg, minor template amount of 15 pg, and mixture ratio of 1:4. For the minor contributor, EFM and STRmix gave profile log_10_(LR) values of 11.3 and 3.6, respectively, leading to a 7.6 difference in log_10_ scale (Additional file 5A). A further review of the per-Locus LR tables obtained from EFM and STRmix for the minor contributor indicated that the LR of D1S1656 had the largest difference (Additional file 5B). The known genotypes at this locus for major and minor contributors were (12,15) and (13,14), respectively, showing that allele “13” dropped-out. A review of the STRmix deconvolution indicated that the genotype at that locus (Q,14) is accepted with a low assigned weight (Additional file 5C). The weights in STRmix are used for LR assignments [87], hence the low D1S1656 LR value.

Therefore, differences observed in profile LRs between the STRmix and EFM maybe partly influenced by the analyst’s review of data and analyst’s decisions when interpreting DNA typing results, different modeling assumptions and statistical models between the two software (e.g. degradation’s effect on peak height, peak height variability, heterozygote balance, drop-in/drop-out, and different stutter types), parameter values settings, and how each software is implementing deconvolution and LR calculations [95]. Different analysts may make different decisions when interpreting the same EPG, thus leading to different LRs even if when using the same software [64]. Upon changing models (e.g. modeling double-back and forward stutter) and/or changing parameter values (e.g. adding a per-dye detection thresholds in EFM v3.2.0, parameters from model maker and profiling kit in STRmix generated from internal validation studies) the resulting LRs will vary to some degree. Different algorithms will also lead to different deconvolution and LR values for the same DNA profile; EFM uses maximum likelihood approaches and STRmix uses Bayesian or MCMC approaches [24].

### 3.7. The verbal equivalents resulting from the numeric LR values from STRmix and EFM

The numeric LR values can be accompanied by a verbal expression, a qualitative statement used in court to describe the degree of support of the findings for one of the propositions relative to the alternative proposition [100–102]. As an exercise for this study we assessed if differences in the quantitative LRs assigned by the two different LR systems resulted in the same or different verbal expressions for both the H1-true tests and H2-true tests. The LR values assigned by STRmix and EFM were binned into their corresponding verbal categories based on the verbal convention recommendations set by the Scientific Working Group on DNA Analysis Methods (SWGDAM) [103] (shown in Table 8). The SWGDAM verbal scale is composed of 5 verbal categories: ‘uninformative’, ‘limited’, ‘moderate’, ‘strong’, and ‘very strong’ for both H1 and H2 support. Each category is associated with a bracket of numerical range of LR values as shown in Table 8.

**Table 8:** The SWGDAM [103] verbal scale for the expression of the likelihood ratios. The table used to bin the numerical LR values into their corresponding verbal qualifiers was adopted from Buckleton et al. [92].

| LR for H1 Support and 1/LR for H2 Support |  | Verbal Qualifier |
| --- | --- | --- |
| Numerical range of LR values used by SWGDAM | Numerical range of LR values used in this work adopted from [92] |  |
| 1 | $1 \leq LR < 2$ | Uninformative |
| 2 – 99 | $2 \leq LR < 100$ | Limited Support |
| 100 – 9,999 | $100 \leq LR < 10,000$ | Moderate Support |
| 10,000 – 999,999 | $10,000 \leq LR < 1,000,000$ | Strong Support |
| $\geq 1,000,000$ | $1,000,000 \leq LR \leq \infty$ | Very Strong Support |

For the H1-true tests (Tables 9A, 9B, and 9C and Supplementary Table 13), the changes in the verbal statements increased with an increase in the number of contributors. The following analysis were binned into identical verbal categories: 96.42 % (297 out of 308) of the LRs from 2P mixtures, 89.11% (393 out of 441) of the LR values of 3P mixtures, and 86.61% (440 out of 508) of the LRs of the 4P mixtures. Hence, (11 out of 308) of the LRs of 2P samples, (48 out of 441) of the LRs of 3P samples, and (68 out of 508) of the LRs of 4P samples were classified into different categories (Table 9A, 9B, and 9C). For the 11 2P cases that were different verbally, 6 were placed in the neighboring categories (for example, for the same 2P profile, an LR from one software was binned into ‘moderate support’ and the LR from the other software was placed in the ‘strong support’ category). The other 5 cases were located in non-adjacent categories and differed by two or more than two verbal categories (e.g. ‘moderate support’ and ‘very strong support’ or ‘exclusionary’ and ‘limited’ or ‘exclusionary’ and ‘strong support’ or ‘exclusionary’ and ‘very strong support’) (Table 9A). With 3P analysis, (6 cases out of 48) were classified into non-adjacent categories: 4 cases were two categories away (‘exclusion’ and ‘limited support’ or ‘limited support’ and ‘strong support’) and 2 cases were different by three categories (‘exclusion’ and ‘moderate support’) (Table 9B). For the LRs of the 4P (Table 9C) analysis that fell in different categories, only 7 out of 68 cases were different by more than one verbal category: 5 cases were different by two categories (“Exclusion” and “Limited Support” or “Limited Support” and “Strong Support” or “Very Strong Support” and “Moderate Support”), and 2 cases were three categories away: (“Exclusion” and “Moderate Support” or “Very Strong Support” and “Limited Support”). Cases of LRs with more than one category difference corresponded to H1-true tests in which POI was a minor contributor and/or had low template amount (Supplementary Table 13).

**Table 9:** Concordance/discordance tables of the binned LR values assigned by STRmix and EFM into their verbal equivalents.

|  |  |  |  |  |
| --- | --- | --- | --- | --- |
| **1 bin apart** | **2 bins apart** | **3 bins apart** | **4 bins apart** | **5 bins apart** |

| **A. 2 person mixtures (H1-True Tests) N=308** |  |  |  |  |  |  |
| --- | --- | --- | --- | --- | --- | --- |
|  | **EFM** |  |  |  |  |  |
| **STRmix** | Exclusionary | Uninformative | Limited | Moderate | Strong | Very Strong |
| Exclusionary | 0 | 0 | 1 | 0 | 1 | 2** |
| Uninformative | 1 | 0 | 0 | 0 | 0 | 0 |
| Limited | 0 | 0 | 2 | 0 | 0 | 0 |
| Moderate | 0 | 0 | 0 | 7 | 1 | 1 |
| Strong | 0 | 0 | 0 | 0 | 9 | 1 |
| Very Strong | 0 | 0 | 0 | 0 | 3 | 279 |
| **B. 3 person mixtures (H1-True Tests) N=441** |  |  |  |  |  |  |
|  | **EFM** |  |  |  |  |  |
| **STRmix** | Exclusionary | Uninformative | Limited | Moderate | Strong | Very Strong |
| Exclusionary | 3 | 5 | 2 | 2 | 0 | 0 |
| Uninformative | 0 | 0 | 0 | 0 | 0 | 0 |
| Limited | 0 | 1 | 15 | 8 | 2 | 0 |
| Moderate | 0 | 0 | 2 | 18 | 8 | 0 |
| Strong | 0 | 0 | 0 | 6 | 23 | 7 |
| Very Strong | 0 | 0 | 0 | 0 | 5 | 334 |
| **C. 4 person mixtures (H1- True Tests) N=508** |  |  |  |  |  |  |
|  | **EFM** |  |  |  |  |  |
| **STRmix** | Exclusionary | Uninformative | Limited | Moderate | Strong | Very Strong |
| Exclusionary | 2 | 4 | 3 | 1 | 0 | 0 |
| Uninformative | 0 | 0 | 7 | 0 | 0 | 0 |
| Limited | 0 | 1 | 27 | 9 | 1 | 0 |
| Moderate | 0 | 0 | 1 | 64 | 10 | 0 |
| Strong | 0 | 0 | 0 | 4 | 59 | 13 |
| Very Strong | 0 | 0 | 1 | 1 | 12 | 288 |

| **D. 2 person mixtures (H2-True Tests) N=308** |  |  |  |  |  |  |
| --- | --- | --- | --- | --- | --- | --- |
|  | **EFM** |  |  |  |  |  |
| **STRmix** | Very Strong | Strong | Moderate | Limited | Uninformative | Inclusionary |
| Very Strong | **239** | **21** | **17** | **8** | **1** | **1** |
| Strong | **0** | **1** | **2** | **3** | **0** | **3** |
| Moderate | **0** | **0** | **2** | **1** | **1** | **0** |
| Limited | **0** | **0** | **0** | **5** | **1** | **1** |
| Uninformative | **0** | **0** | **0** | **0** | **0** | **0** |
| Inclusionary | **0** | **0** | **0** | **0** | **0** | **1** |

| **E. 3 person mixtures (H2-True Tests) N=441** |  |  |  |  |  |  |
| --- | --- | --- | --- | --- | --- | --- |
|  | **EFM** |  |  |  |  |  |
| **STRmix** | Very Strong | Strong | Moderate | Limited | Uninformative | Inclusionary |
| Very Strong | **107** | **53** | **77** | **74** | **9** | **10** |
| Strong | **0** | **0** | **7** | **21** | **8** | **2** |
| Moderate | **0** | **0** | **1** | **12** | **14** | **14** |
| Limited | **0** | **0** | **0** | **10** | **7** | **13** |
| Uninformative | **0** | **0** | **0** | **0** | **0** | **0** |
| Inclusionary | **0** | **0** | **0** | **0** | **0** | **2** |

| **F. 4 person mixtures (H2-True Tests) N=508** |  |  |  |  |  |  |
| --- | --- | --- | --- | --- | --- | --- |
|  | **EFM** |  |  |  |  |  |
| **STRmix** | Very Strong | Strong | Moderate | Limited | Uninformative | Inclusionary |
| Very Strong | **11** | **15** | **33** | **71** | **32** | **53** |
| Strong | **2** | **0** | **2** | **34** | **15** | **14** |
| Moderate | **0** | **0** | **2** | **41** | **37** | **27** |
| Limited | **1** | **0** | **0** | **16** | **32** | **50** |
| Uninformative | **0** | **0** | **0** | **0** | **0** | **6** |
| Inclusionary | **0** | **0** | **0** | **0** | **2** | **12** |

The categories used for the binning the LRs of the H2-true tests are in favor of H2 over H1 (i.e., mirror image of the verbal scale of the H1-true tests). For the H2-true tests, similarly as for the H1-true tests, as the number of contributors increased the differences in the verbal statements increased as well (Tables 9D, 9E, and 9F and Supplementary Table 14). The following analysis were binned into the same verbal category: 80.51 % (248 out of 308) of the LRs from 2P mixtures, 27.21% (120 out of 441) of the LR values of 3P mixtures, and 8.07% (41 out of 508) of the LRs of the 4P mixtures (Tables 9D, 9E, and 9F). For the 60 2P cases that were different verbally, 35 were placed in non-neighboring categories (Table 9D). With 3P and 4P analysis, (242 cases out of 321) and (367 cases out of 467), respectively, were classified into non-adjacent categories (Tables 9E and 9F).

The tables display the results of the categorization of the LRs for both the H1-true tests of (A) 2P where ** are the two 2P H1-true test interpretations for which STRmix assigned profile LRs of 0 (binned into the Exclusionary category and discussed in detail in Section 3.6), (B) 3P, and (C) 4P and H2-true tests of (D) 2P, (E) 3P, and (F) 4P generated in STRmix and EFM into their corresponding verbal expression. Also, the tables demonstrate the observed differences in the verbal expressions between the two LR systems. The number of cases that resulted in same verbal expression between STRmix and EFM fell inside the diagonal (white cells). All the numbers outside the diagonal (shaded cells) are indication of cases where LRs from both software were classified into different categories and resulted in shifting by one or more than one verbal category (indicated by different shades as shown by the legend). Values in and above the diagonal are the results of the verbal expression of LRs produced in EFM while values in and below the diagonals are the results of the verbal expression of LR values assigned by STRmix. The verbal expressions are shown at the top and left edges of the tables.

## 4. Conclusion

In this independent study, we examined the discrimination performance as well as LR values assigned by two LR systems using two continuous PGS built on different modelling assumptions, STRmix (proprietary) and EFM (open-source) [12, 24]. We use the term LR system deliberately to emphasize that the assigned LR values are a product of the decisions that went into the interpretation process of the LR system and not solely the PGS. For example, our specific choice of the PROVEDIt filtered files, protocols used for the data analysis in both STRmix and EFM, decision to use the known NOC, and to provide similar data (EPGs) into both software are specific to “our” LR system used in this study. We recognize that alternative decisions could have been made, and thus different LR values could have been assigned. We described the degree of differences in the LR values, where the differences occur, and the potential explanations for the observed differences. We analyzed 154 2P, 147 3P, and 127 4P mixture profiles from PROVEDIt database [71, 72] of varying DNA quality, DNA quantity, and mixture ratios (shown in Supplementary Tables 4 and 5). Both H1-true tests (Supplementary Table 4) and H2-true tests (Supplementary Table 5) for the 2P, 3P, and 4P were analyzed in both STRmix and EFM yielding a total of 1,257 of known-contributor LRs and 1,257 of known non-contributor LRs from each software.

The discrimination performance was evaluated qualitatively (Fig. 3) and quantitatively (Fig. 5) by checking the ability of each LR system in discriminating between H1-true and H2-true scenarios. The overall distribution plots (Fig. 3) and ROC plots (Fig. 5) suggest that the ability of the two LR systems to discriminate between known contributors and known non-contributors in aggregate are statistically indistinguishable for the data we considered.

Although both LR systems had similar discrimination performance, that did not imply that STRmix and EFM assigned equal LR values on a case-by-case basis even though LR computations were based on same/fixed EPG features, same pair of propositions, NOC, theta, and population allele frequency (Fig. 6). The magnitude of differences was broken down into factor of 10 bins (Fig. 7) and stratified by the type of POI (Fig. 8). Differences in LR values greater than or equal to 3 on the log_10_ scale (as discussed in Section 3.6) were investigated and could occur due to one or more of the following reasons:

(1) decisions made during parameters settings (e.g. choice of profiles for Model Maker interpretation and choice of settings for analysis such as analytical thresholds and drop-in parameters)
(2) decision to analyze the same input files in both STRmix and EFM of which some of these profiles contained stutter peaks (F1 and B2) that were not modelled by EFM v2.1.0
(3) non-convergence of the MCMC algorithms
(4) differences in modelling assumptions of peak height information and variability, degradation, heterozygote balance, and allelic drop-outs/drop-ins

It is important to note that the apparent differences observed due to mentioned factors (2) and (3) were reduced upon re-interpretation of data both manually and in the software (e.g. re-interpreting profiles in EFM after removing the unmodelled F1 and B2 (Supplementary Table 11) or repeating analysis in STRmix with higher number of accepts (Supplementary Table 8).

Irrespective of the quantitative differences observed in certain cases between the LR systems (Fig. 6), there seems to be a pattern observed in this study. Differences in LR values were observed in both directions (e.g., when LR STRmix ≥ 1000*LR EFM or when LR EFM ≥ 1000*LR STRmix). The magnitude of the differences was greater with minor donors than with equal or major contributors (Fig. 8 and Supplementary Tables 11 and 12). Similar observations were documented in [30, 62, 70] when comparing LRs from various models.

Both LR systems showed adventitious exclusionary LR values (LR < 1) for H1-true tests (mainly with minor contributors) (Fig. 4 and Supplementary Table 6) and adventitious inclusionary LR values (LR > 1) for H2-true tests (Fig. 4 and Supplementary Table 7). The largest LR assigned using our LR systems and dataset was 587 from STRmix and 167 from EFM for a known unrelated non-contributor in the 1,257 H2-true tests (Fig. 4C and Supplementary Table 7).

We observed that in certain cases differences in numerical LR values from both software resulted in differences in one or more than one verbal categories (Table 9). These differences were substantially more with low template minor contributors and higher NOC (Table 9 and Supplementary Tables 13 and 14); observations that have as well been examined in Swaminathan et al. [62]. Also, the cases of differences in the numerical LR values and verbal classification of the H2-true tests between the two models were higher than the ones observed with H1-true tests (Figs. 6 and 7 and Table 9), thus showing the differences in the ability of both models to evaluate/measure the strength of evidence. The comparison of the assigned LR values in the verbal scale framework was included to provide some context to the observed differences. Although interesting, observed differences greater than 10^3^ may have less practical impact for large LRs (e.g. 10^15^ versus 10^18^) as compared to smaller LRs (e.g. 10^1^ versus 10^4^).

The findings of this study are specific to the LR systems (Fig. 1) used in our study: (i) data chosen to generate parameter values and settings for analysis (e.g. Model Maker, analytical thresholds, drop-in, stutter settings), (ii) decisions made prior to the analysis of the mixture profiles in both software, and (iii) mixture profiles used for LR assessments. The profiles used for generating parameter values are shown in Supplementary Tables 1, 2, and 3. We also share with the forensic community the mixture profiles used for H1-true and H2-true tests with their corresponding LR values from both LR systems (Supplementary Tables 4 and 5). The comparisons performed in this study are more extensive than any software comparisons previously reported [1, 29, 30, 52, 54, 62, 65–67, 69, 70]. The included supplementary tables and figures are intended to provide an example of the level of information and transparency we desire to see in similar DNA mixture publications. This provides the opportunity to review a specific mixture profile and further examine the assigned LR value(s). We believe that sharing the assigned LR values correlated with each mixture vs POI comparison complements the global aggregate level ROC and scatter plots used to assess the LR systems. This was further enabled by using the publicly available and consented PROVEDIt mixture profiles (i.e., the sharing of DNA profiles was not an issue). We encourage other investigators to assess the PROVEDIt profiles with *their* LR systems, compare their assigned LR values to those obtained in this study, and/or develop further visualization tools.

To sum up, “there are no true likelihood ratios, just like there are no true models” [104] and “no model perfectly incorporates all sources of uncertainty” [95]. The focus of this study is not to suggest that any one of the software is based on a true or best model. Our intent is to demonstrate the value of using a publicly available ground truth known mixture data [71] to assess performance of any LR system and describe how examining more than one PGS with similar discrimination power can be beneficial and an additional empirical diagnostic check even if software in use does contain certain diagnostic statistics as part of the output.

## Acknowledgements

The authors would like to thank John Butler and Arun Moorthy at NIST for critically reading the manuscript. The authors would also like to thank Øyvind Bleka (Oslo University Hospital), Zane Kerr and Judi Morawitz (Environmental Science and Research), and Steven Myers (CAL DOJ) for their input on using the software and meaningful discussions on data analysis.

## References

[1] J.A. Bright, I.W. Evett, D. Taylor, J.M. Curran, J. Buckleton, A series of recommended tests when validating probabilistic DNA profile interpretation software, Forensic science international. Genetics 14 (2015) 125–31.

[2] D. Taylor, J.A. Bright, J. Buckleton, The interpretation of single source and mixed DNA profiles, Forensic science international. Genetics 7(5) (2013) 516–28.

[3] M.W. Perlin, M.M. Legler, C.E. Spencer, J.L. Smith, W.P. Allan, J.L. Belrose, B.W. Duceman, Validating TrueAllele® DNA mixture interpretation, Journal of forensic sciences 56(6) (2011) 1430–47.

[4] J.A. Bright, J.S. Buckleton, D. Taylor, M.A. Fernando, J.M. Curran, Modeling forward stutter: toward increased objectivity in forensic DNA interpretation, Electrophoresis 35(21-22) (2014) 3152–7.

[5] D. Taylor, J.A. Bright, H. Kelly, M.H. Lin, J. Buckleton, A fully continuous system of DNA profile evidence evaluation that can utilise STR profile data produced under different conditions within a single analysis, Forensic science international. Genetics 31 (2017) 149–154.

[6] J.S. Buckleton, J.A. Bright, S. Gittelson, T.R. Moretti, A.J. Onorato, F.R. Bieber, B. Budowle, D.A. Taylor, The Probabilistic Genotyping Software STRmix: Utility and Evidence for its Validity, Journal of forensic sciences 64(2) (2019) 393–405.

[7] SWGDAM, Guidelines for the validation of probabilistic genotyping systems, (2015).

[8] T.R. Moretti, R.S. Just, S.C. Kehl, L.E. Willis, J.S. Buckleton, J.A. Bright, D.A. Taylor, A.J. Onorato, Internal validation of STRmix™ for the interpretation of single source and mixed DNA profiles, Forensic science international. Genetics 29 (2017) 126–144.

[9] J.A. Bright, D. Taylor, S. Gittelson, J. Buckleton, The paradigm shift in DNA profile interpretation, Forensic science international. Genetics 31 (2017) e24–e32.

[10] H. Kelly, J.A. Bright, J.S. Buckleton, J.M. Curran, A comparison of statistical models for the analysis of complex forensic DNA profiles, Science & justice : journal of the Forensic Science Society 54(1) (2014) 66–70.

[11] M.D. Coble, J.A. Bright, Probabilistic genotyping software: An overview, Forensic science international. Genetics 38 (2019) 219–224.

[12] J.A. Bright, D. Taylor, C. McGovern, S. Cooper, L. Russell, D. Abarno, J. Buckleton, Developmental validation of STRmix™, expert software for the interpretation of forensic DNA profiles, Forensic science international. Genetics 23 (2016) 226–239.

[13] D.J. Balding, J. Buckleton, Interpreting low template DNA profiles, Forensic science international. Genetics 4(1) (2009) 1–10.

[14] J.M. Butler, S. Willis, Interpol review of forensic biology and forensic DNA typing 2016-2019, Forensic science international 2 (2020) 352–367.

[15] STRmix™ forensic software. <https://www.strmix.com/>).

[16] M.W. Perlin, A. Sinelnikov, An information gap in DNA evidence interpretation, PloS one 4(12) (2009) e8327.

[17] TrueAllele ® DNA Interpretation. <https://www.cybgen.com/>).

[18] F.M. Götz, H. Schönborn, V. Borsdorf, A.-M. Pflugbeil, D. Labudde, GenoProof Mixture 3—New software and process to resolve complex DNA mixtures, Forensic Science International: Genetics Supplement Series 6 (2017) e549–e551.

[19] GenoProof Mixture 3. <https://www.qualitype.de/en/solutions/products/evaluation-software/genoproof-mixture/>).

[20] MaSTR™ Software. <https://softgenetics.com/MaSTR.php>).

[21] The DNA·VIEW® Mixture Solution. <http://dna-view.com/>).

[22] R. Puch-Solis, T. Clayton, Evidential evaluation of DNA profiles using a discrete statistical model implemented in the DNA LiRa software, Forensic science international. Genetics 11 (2014) 220–8.

[23] LiRa. <https://cdnmedia.eurofins.com/european-west/media/1418957/lgc_lira_fact_sheet_en_0815_90.pdf>).

[24] Ø. Bleka, G. Storvik, P. Gill, EuroForMix: An open source software based on a continuous model to evaluate STR DNA profiles from a mixture of contributors with artefacts, Forensic science international. Genetics 21 (2016) 35–44.

[25] EuroForMix. <http://www.euroformix.com/>).

[26] H. Swaminathan, A. Garg, C.M. Grgicak, M. Medard, D.S. Lun, CEESIt: A computational tool for the interpretation of STR mixtures, Forensic science international. Genetics 22 (2016) 149–160.

[27] ValiDNA: Forensic DNA Software. <https://lftdi.camden.rutgers.edu/provedit/software/>).

[28] likeLTD (likelihoods for low-template DNA profiles). <https://sites.google.com/site/baldingstatisticalgenetics/software/likeltd-r-forensic-dna-r-code>).

[29] Y. You, D. Balding, A comparison of software for the evaluation of complex DNA profiles, Forensic science international. Genetics 40 (2019) 114–119.

[30] S. Manabe, C. Morimoto, Y. Hamano, S. Fujimoto, K. Tamaki, Development and validation of open-source software for DNA mixture interpretation based on a quantitative continuous model, 12(11) (2017) e0188183.

[31] Kongoh. <https://github.com/manabe0322/Kongoh>).

[32] R.G. Cowell, T. Graversen, S.L. Lauritzen, J. Mortera, Analysis of forensic DNA mixtures with artefacts, Journal of the Royal Statistical Society: Series C (Applied Statistics) 64(1) (2015) 1–48.

[33] DNAmixtures. <http://dnamixtures.r-forge.r-project.org/>).

[34] C.C.G. Benschop, J. Hoogenboom, P. Hovers, M. Slagter, D. Kruise, R. Parag, K. Steensma, K. Slooten, J.H.A. Nagel, P. Dieltjes, V. van Marion, H. van Paassen, J. de Jong, C. Creeten, T. Sijen, A.L.J. Kneppers, DNAxs/DNAStatistX: Development and validation of a software suite for the data management and probabilistic interpretation of DNA profiles, Forensic science international. Genetics 42 (2019) 81–89.

[35] DNAxs/DNAStatistX. <https://www.forensicinstitute.nl/research-and-innovation/international-projects/dnaxs>).

[36] BulletProof probabilistic genotyping software. <http://ednalims.com/probabilistic-genotyping/>).

[37] J.A. Bright, D. Taylor, J.M. Curran, J.S. Buckleton, Developing allelic and stutter peak height models for a continuous method of DNA interpretation, Forensic science international. Genetics 7(2) (2013) 296–304.

[38] R. Puch-Solis, L. Rodgers, A. Mazumder, S. Pope, I. Evett, J. Curran, D. Balding, Evaluating forensic DNA profiles using peak heights, allowing for multiple donors, allelic dropout and stutters, Forensic science international. Genetics 7(5) (2013) 555–63.

[39] R.G. Cowell, S.L. Lauritzen, J. Mortera, A gamma model for DNA mixture analyses, Bayesian Anal. 2(2) (2007) 333–348.

[40] R. Cowell, T. Graversen, S. Lauritzen, J. Mortera, Analysis of forensic DNA mixtures with artefacts, Journal of the Royal Statistical Society: Series C (Applied Statistics) 64(1) (2015) 1–48.

[41] D.J. Balding, R.A. Nichols, DNA profile match probability calculation: how to allow for population stratification, relatedness, database selection and single bands, Forensic science international 64(2-3) (1994) 125–40.

[42] D.V. Lindley, A problem in forensic science, Biometrika 64(2) (1977) 207–213.

[43] I.W. Evett, C. Buffery, G. Willott, D. Stoney, A guide to interpreting single locus profiles of DNA mixtures in forensic cases, Journal - Forensic Science Society 31(1) (1991) 41–7.

[44] P. Gill, C.H. Brenner, J.S. Buckleton, A. Carracedo, M. Krawczak, W. Mayr, N. Morling, M. Prinz, P.M. Schneider, B. Weir, DNA commission of the International Society of Forensic Genetics: recommendations on the interpretation of mixtures, Forensic science international 160(2-3) (2006) 90–101.

[45] G. Jackson, S. Jones, G. Booth, C. Champod, I.W. Evett, The nature of forensic science opinion—a possible framework to guide thinking and practicce in investigation and in court proceedings, Science & Justice 46(1) (2006) 33–44.

[46] SWGDAM, Guidelines for the validation of probabilistic genotyping systems. <(http://media.wix.com/ugd/4344b0_22776006b67c4a32a5ffc04fe3b56515.pdf),>, (2015)).

[47] M.D. Coble, J. Buckleton, J.M. Butler, T. Egeland, R. Fimmers, P. Gill, L. Gusmão, B. Guttman, M. Krawczak, N. Morling, W. Parson, N. Pinto, P.M. Schneider, S.T. Sherry, S. Willuweit, M. Prinz, DNA Commission of the International Society for Forensic Genetics: Recommendations on the validation of software programs performing biostatistical calculations for forensic genetics applications, Forensic science international. Genetics 25 (2016) 191–197.

[48] UK Forensic Science Regulator, Software validation for DNA mixture interpretation, FSR-G-223. <https://assets.publishing.service.gov.uk/government/uploads/system/uploads/attachment_data/file/740877/G223_Mixtures_software_validation_Issue1.pdf>).

[49] President’s Council of Advisors on Science and Technology. Report to the President - Forensic Science in Criminal Courts: Ensuring Scientific Validity of Feature-Comparison Methods <https://obamawhitehouse.archives.gov/sites/default/files/microsites/ostp/PCAST/pcast_forensic_science_report_final.pdf>, (2016)).

[50] J.A. Bright, K. Cheng, Z. Kerr, C. McGovern, H. Kelly, T.R. Moretti, M.A. Smith, F.R. Bieber, B. Budowle, M.D. Coble, R. Alghafri, P.S. Allen, A. Barber, V. Beamer, C. Buettner, M. Russell, C. Gehrig, T. Hicks, J. Charak, K. Cheong-Wing, A. Ciecko, C.T. Davis, M. Donley, N. Pedersen, B. Gartside, D. Granger, M. Greer-Ritzheimer, E. Reisinger, J. Kennedy, E. Grammer, M. Kaplan, D. Hansen, H.J. Larsen, A. Laureano, C. Li, E. Lien, E. Lindberg, C. Kelly, B. Mallinder, S. Malsom, A. Yacovone- Margetts, A. McWhorter, S.M. Prajapati, T. Powell, G. Shutler, K. Stevenson, A.R. Stonehouse, L. Smith, J. Murakami, E. Halsing, D. Wright, L. Clark, D.A. Taylor, J. Buckleton, STRmix™ collaborative exercise on DNA mixture interpretation, Forensic science international. Genetics 40 (2019) 1–8.

[51] J.A. Bright, R. Richards, M. Kruijver, H. Kelly, C. McGovern, A. Magee, A. McWhorter, A. Ciecko, B. Peck, C. Baumgartner, C. Buettner, S. McWilliams, C. McKenna, C. Gallacher, B. Mallinder, D. Wright, D. Johnson, D. Catella, E. Lien, C. O’Connor, G. Duncan, J. Bundy, J. Echard, J. Lowe, J. Stewart, K. Corrado, S. Gentile, M. Kaplan, M. Hassler, N. McDonald, P. Hulme, R.H. Oefelein, S. Montpetit, M. Strong, S. Noël, S. Malsom, S. Myers, S. Welti, T. Moretti, T. McMahon, T. Grill, T. Kalafut, M. Greer-Ritzheimer, V. Beamer, D.A. Taylor, J.S. Buckleton, Internal validation of STRmix™ - A multi laboratory response to PCAST, Forensic science international. Genetics 34 (2018) 11–24.

[52] J.S. Buckleton, J.A. Bright, K. Cheng, B. Budowle, M.D. Coble, NIST interlaboratory studies involving DNA mixtures (MIX13): A modern analysis, Forensic science international. Genetics 37 (2018) 172–179.

[53] D.W. Bauer, N. Butt, J.M. Hornyak, M.W. Perlin, Validating TrueAllele(®) Interpretation of DNA Mixtures Containing up to Ten Unknown Contributors, Journal of forensic sciences 65(2) (2020) 380–398.

[54] C.C.G. Benschop, A. Nijveld, F.E. Duijs, T. Sijen, An assessment of the performance of the probabilistic genotyping software EuroForMix: Trends in likelihood ratios and analysis of Type I & II errors, Forensic science international. Genetics 42 (2019) 31–38.

[55] H. Kelly, J.A. Bright, M. Kruijver, S. Cooper, D. Taylor, K. Duke, M. Strong, V. Beamer, C. Buettner, J. Buckleton, A sensitivity analysis to determine the robustness of STRmix™ with respect to laboratory calibration, Forensic science international. Genetics 35 (2018) 113–122.

[56] M.H. Lin, J.A. Bright, S.N. Pugh, J.S. Buckleton, The interpretation of mixed DNA profiles from a mother, father, and child trio, Forensic science international. Genetics 44 (2020) 102175.

[57] D. Taylor, Using continuous DNA interpretation methods to revisit likelihood ratio behaviour, Forensic science international. Genetics 11 (2014) 144–53.

[58] J.A. Bright, K.E. Stevenson, J.M. Curran, J.S. Buckleton, The variability in likelihood ratios due to different mechanisms, Forensic science international. Genetics 14 (2015) 187–90.

[59] J.S. Buckleton, J.A. Bright, K. Cheng, H. Kelly, D.A. Taylor, The effect of varying the number of contributors in the prosecution and alternate propositions, Forensic science international. Genetics 38 (2019) 225–231.

[60] J.A. Bright, J.M. Curran, J.S. Buckleton, The effect of the uncertainty in the number of contributors to mixed DNA profiles on profile interpretation, Forensic science international. Genetics 12 (2014) 208–14.

[61] S. Cooper, C. McGovern, J.A. Bright, D. Taylor, J. Buckleton, Investigating a common approach to DNA profile interpretation using probabilistic software, Forensic science international. Genetics 16 (2015) 121–131.

[62] H. Swaminathan, M.O. Qureshi, C.M. Grgicak, K. Duffy, D.S. Lun, Four model variants within a continuous forensic DNA mixture interpretation framework: Effects on evidential inference and reporting, 13(11) (2018) e0207599.

[63] J. Hannig, S. Riman, H. Iyer, P.M. Vallone, Are reported likelihood ratios well calibrated?, Forensic Science International: Genetics Supplement Series 7(1) (2019) 572–574.

[64] P.A. Barrio, M. Crespillo, J.A. Luque, M. Aler, C. Baeza-Richer, L. Baldassarri, E. Carnevali, P. Coufalova, I. Flores, O. García, M.A. García, R. González, A. Hernández, V. Inglés, G.M. Luque, A. Mosquera-Miguel, S. Pedrosa, M.L. Pontes, M.J. Porto, Y. Posada, M.I. Ramella, T. Ribeiro, E. Riego, A. Sala, V.G. Saragoni, A. Serrano, S. Vannelli, GHEP-ISFG collaborative exercise on mixture profiles (GHEP-MIX06). Reporting conclusions: Results and evaluation, Forensic science international. Genetics 35 (2018) 156–163.

[65] T.W. Bille, S.M. Weitz, M.D. Coble, J. Buckleton, J.A. Bright, Comparison of the performance of different models for the interpretation of low level mixed DNA profiles, Electrophoresis 35(21-22) (2014) 3125–33.

[66] Ø. Bleka, C.C.G. Benschop, G. Storvik, P. Gill, A comparative study of qualitative and quantitative models used to interpret complex STR DNA profiles, Forensic science international. Genetics 25 (2016) 85–96.

[67] E. Alladio, M. Omedei, S. Cisana, G. D’Amico, D. Caneparo, M. Vincenti, P. Garofano, DNA mixtures interpretation - A proof-of-concept multi-software comparison highlighting different probabilistic methods’ performances on challenging samples, Forensic science international. Genetics 37 (2018) 143–150.

[68] D. Taylor, J. Buckleton, Do low template DNA profiles have useful quantitative data?, Forensic science international. Genetics 16 (2015) 13–16.

[69] C.D. Steele, M. Greenhalgh, D.J. Balding, Evaluation of low-template DNA profiles using peak heights, Statistical applications in genetics and molecular biology 15(5) (2016) 431–445.

[70] K. Slooten, The information gain from peak height data in DNA mixtures, Forensic science international. Genetics 36 (2018) 119–123.

[71] PROVEDIt Database <https://lftdi.camden.rutgers.edu/provedit/files/>).

[72] L.E. Alfonse, A.D. Garrett, D.S. Lun, K.R. Duffy, C.M. Grgicak, A large-scale dataset of single and mixed-source short tandem repeat profiles to inform human identification strategies: PROVEDIt, Forensic science international. Genetics 32 (2018) 62–70.

[73] J.M. Butler, Advanced Topics in Forensic DNA Typing: Interpretation, Elsevier, Amsterdam2015.

[74] J.M. Butler, H.K. Iyer, Validation, Principles, Practices, Parameters, Performance, Evaluations, and Protocols, (ISHI 2020 Validation Workshop).

[75] C.D. Steele, D.J. Balding, Statistical evaluation of forensic DNA profile evidence, Annual Review of Statistics and Its Application 1 (2014) 361–384.

[76] D.A. Taylor, J.A. Bright, J. Buckleton, Commentary: a “source” of error: computer code, criminal defendants, and the constitution, Frontiers in genetics 8 (2017) 33.

[77] J. Bregu, D. Conklin, E. Coronado, M. Terrill, R.W. Cotton, C.M. Grgicak, Analytical thresholds and sensitivity: establishing RFU thresholds for forensic DNA analysis, Journal of forensic sciences 58(1) (2013) 120–9.

[78] STRmix v2.6.0 User Manual and Implementation/Validation Guide. <https://support.strmix.com/>, (2018)).

[79] Manual for EuroForMix v2.1. <http://www.euroformix.com/sites/default/files/euroformixManual_v2_1.pdf>, (2019)).

[80] J.R. Gilder, T.E. Doom, K. Inman, D.E. Krane, Run-specific limits of detection and quantitation for STR-based DNA testing, Journal of forensic sciences 52(1) (2007) 97–101.

[81] D. Taylor, J.A. Bright, C. McGoven, C. Hefford, T. Kalafut, J. Buckleton, Validating multiplexes for use in conjunction with modern interpretation strategies, Forensic science international. Genetics 20 (2016) 6–19.

[82] U.J. Mönich, K. Duffy, M. Médard, V. Cadambe, L.E. Alfonse, C. Grgicak, Probabilistic characterisation of baseline noise in STR profiles, Forensic science international. Genetics 19 (2015) 107–122.

[83] STRmix v2.6.0 Operation Manual. <https://support.strmix.com/>, (2018)).

[84] C. Brookes, J.-A. Bright, S. Harbison, J. Buckleton, Characterising stutter in forensic STR multiplexes, Forensic Science International: Genetics 6(1) (2012) 58–63.

[85] J.A. Bright, J.M. Curran, Investigation into stutter ratio variability between different laboratories, Forensic science international. Genetics 13 (2014) 79–81.

[86] C.R. Steffen, M.D. Coble, K.B. Gettings, P.M. Vallone, Corrigendum to ’U.S. Population Data for 29 Autosomal STR Loci’ [Forensic Sci. Int. Genet. 7 (2013) e82-e83], Forensic science international. Genetics 31 (2017) e36–e40.

[87] L. Russell, S. Cooper, R. Wivell, Z. Kerr, D. Taylor, J. Buckleton, J.-A. Bright, A guide to results and diagnostics within a STRmix™ report, WIREs Forensic Science 1(6) (2019) e1354.

[88] J. Buckleton, J. Curran, J. Goudet, D. Taylor, A. Thiery, B.S. Weir, Population-specific FST values for forensic STR markers: A worldwide survey, Forensic science international. Genetics 23 (2016) 91–100.

[89] D.M. Green, J.A. Swets, Signal detection theory and psychophysics, John Wiley, Oxford, England, 1966.

[90] P. Sonego, A. Kocsor, S. Pongor, ROC analysis: applications to the classification of biological sequences and 3D structures, Briefings in bioinformatics 9(3) (2008) 198–209.

[91] R Core Team. R: A Language and Environment for Statistical Computing. <https://www.r-project.org/>, (R Foundation for Statistical Computing, 2020)).

[92] J. Buckleton, S. Pugh, J.-A. Bright, D. Taylor, J. Curran, M. Kruijver, P. Gill, B. Budowle, K. Cheng, Are low LRs reliable?, Forensic Science International: Genetics (2020) 102350.

[93] S. Noël, J. Noël, D. Granger, J.F. Lefebvre, D. Séguin, STRmix(™) put to the test: 300 000 non-contributor profiles compared to four-contributor DNA mixtures and the impact of replicates, Forensic science international. Genetics 41 (2019) 24–31.

[94] D. Taylor, J. Buckleton, I. Evett, Testing likelihood ratios produced from complex DNA profiles, Forensic science international. Genetics 16 (2015) 165–171.

[95] D. Taylor, D. Balding, How can courts take into account the uncertainty in a likelihood ratio?, Forensic science international. Genetics 48 (2020) 102361.

[96] E.R. DeLong, D.M. DeLong, D.L. Clarke-Pearson, Comparing the areas under two or more correlated receiver operating characteristic curves: a nonparametric approach, Biometrics 44(3) (1988) 837–45.

[97] A. Gelman, D.B. Rubin, Inference from Iterative Simulation Using Multiple Sequences, Statist. Sci. 7(4) (1992) 457–472.

[98] Ø. Bleka, An introduction to EuroForMix. <http://euroformix.com/sites/default/files/EuroForMixTheory_ISFG17.pdf>, 2017).

[99] Ø. Bleka, New update of EuroForMix Version 3.0.0.

[100] P. Gill, T. Hicks, J.M. Butler, E. Connolly, L. Gusmão, B. Kokshoorn, N. Morling, R.A.H. van Oorschot, W. Parson, M. Prinz, P.M. Schneider, T. Sijen, D. Taylor, DNA commission of the International society for forensic genetics: Assessing the value of forensic biological evidence - Guidelines highlighting the importance of propositions. Part II: Evaluation of biological traces considering activity level propositions, Forensic science international. Genetics 44 (2020) 102186.

[101] C. Aitken, F. Taroni, A verbal scale for the interpretation of evidence, Science & Justice 8 (1998) 279–281.

[102] E. Arscott, R. Morgan, G. Meakin, J. French, Understanding forensic expert evaluative evidence: A study of the perception of verbal expressions of the strength of evidence, Science & justice : journal of the Forensic Science Society 57(3) (2017) 221–227.

[103] SWGDAM, Recommendations of the SWGDAM Ad Hoc Working Group on Genotyping Results Reported as Likelihood Ratios. <https://docs.wixstatic.com/ugd/4344b0_dd5221694d1448588dcd0937738c9e46.pdf>, (2018)).

[104] P. Gill, T. Hicks, J.M. Butler, E. Connolly, L. Gusmão, B. Kokshoorn, N. Morling, R.A.H. van Oorschot, W. Parson, M. Prinz, P.M. Schneider, T. Sijen, D. Taylor, DNA commission of the International society for forensic genetics: Assessing the value of forensic biological evidence - Guidelines highlighting the importance of propositions: Part I: evaluation of DNA profiling comparisons given (sub-) source propositions, Forensic science international. Genetics 36 (2018) 189–202.

