## Additional File 3 for "Examining Discrimination Performance and Likelihood Ratio Values for Two Different Likelihood Ratio Systems Using the Provedit Dataset"

### Slide 1
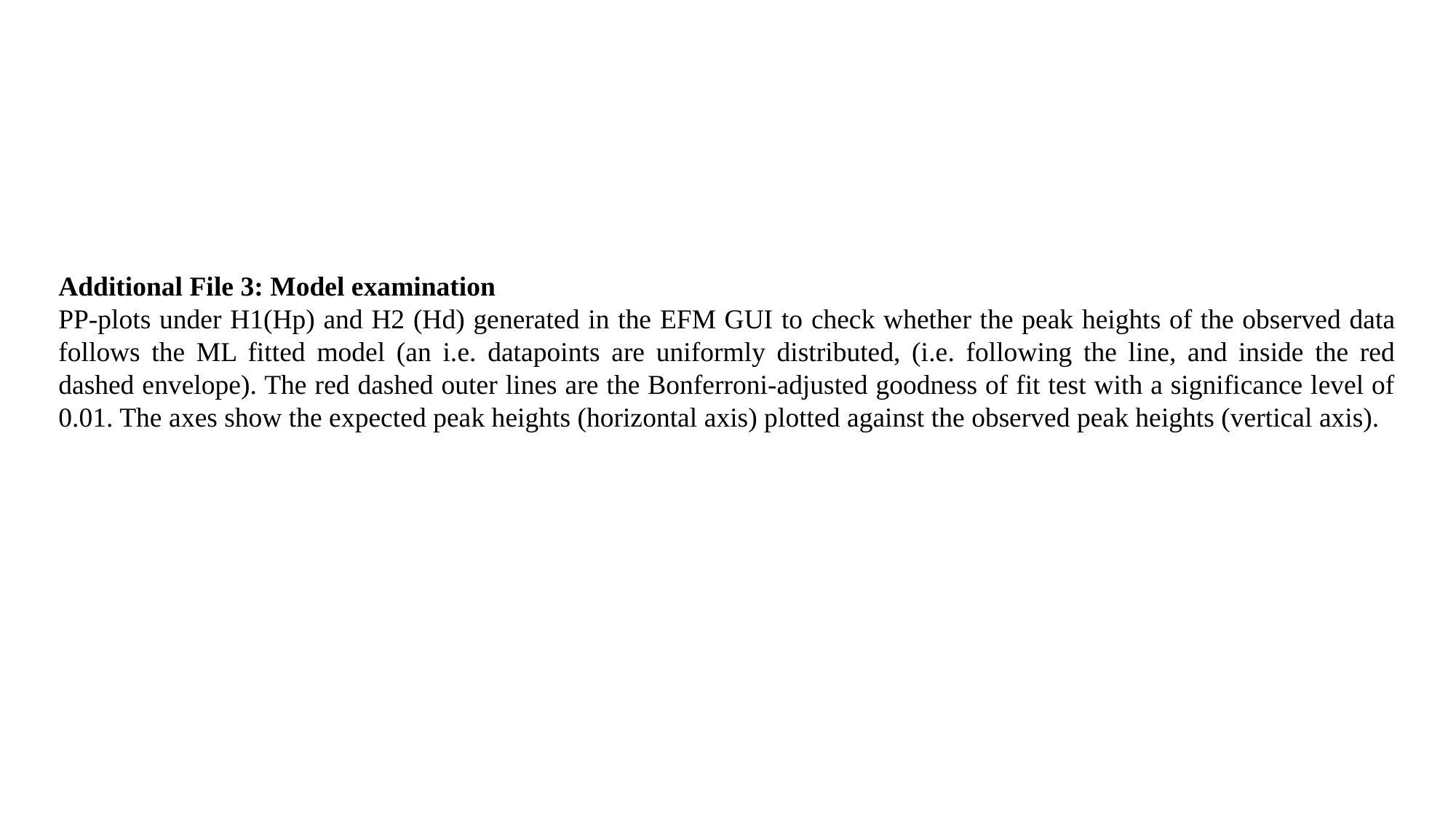

Additional File 3: Model examination
PP-plots under H1(Hp) and H2 (Hd) generated in the EFM GUI to check whether the peak heights of the observed data follows the ML fitted model (an i.e. datapoints are uniformly distributed, (i.e. following the line, and inside the red dashed envelope). The red dashed outer lines are the Bonferroni-adjusted goodness of fit test with a significance level of 0.01. The axes show the expected peak heights (horizontal axis) plotted against the observed peak heights (vertical axis).

### Slide 2
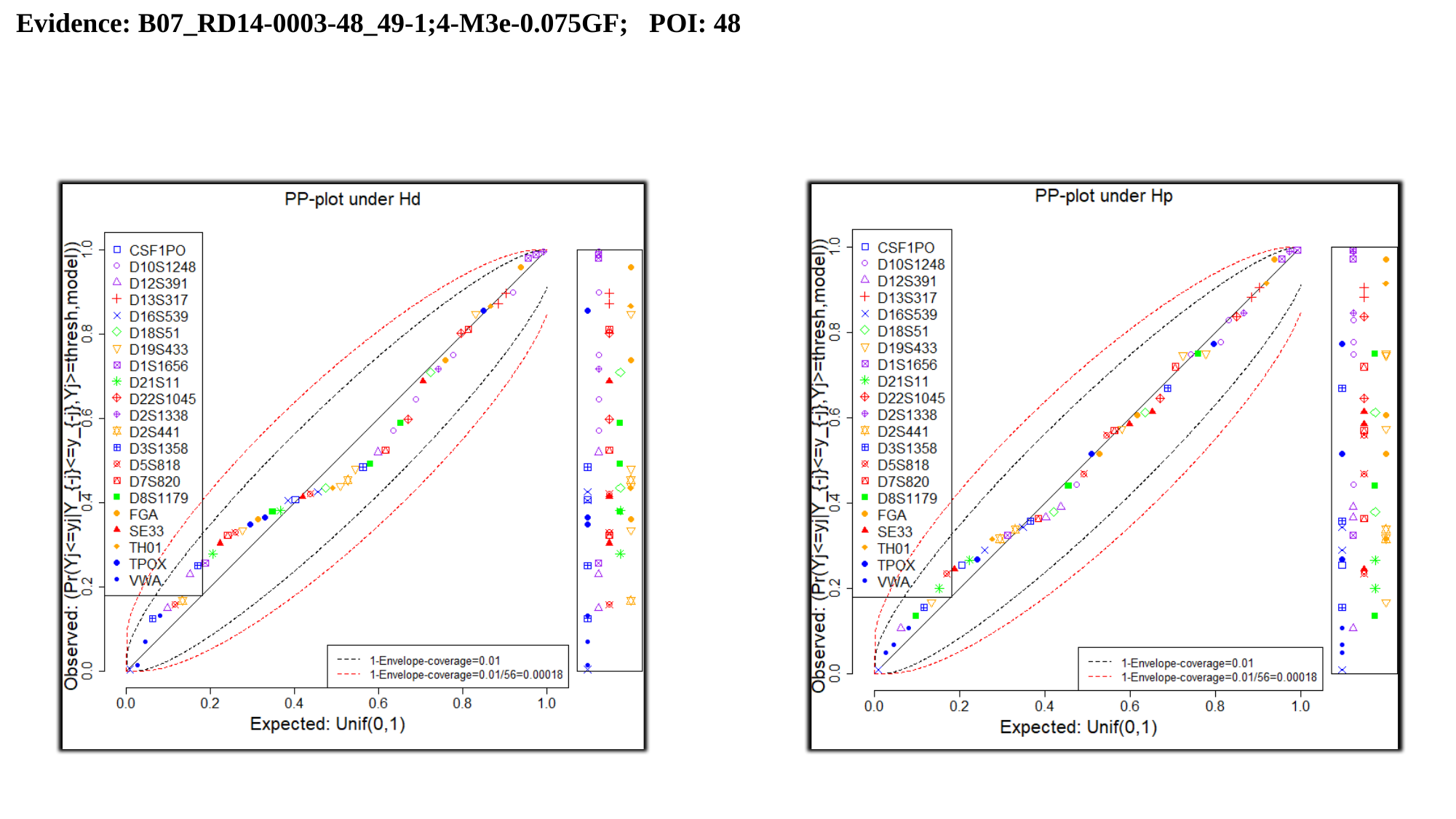

Evidence: B07_RD14-0003-48_49-1;4-M3e-0.075GF; POI: 48

### Slide 3
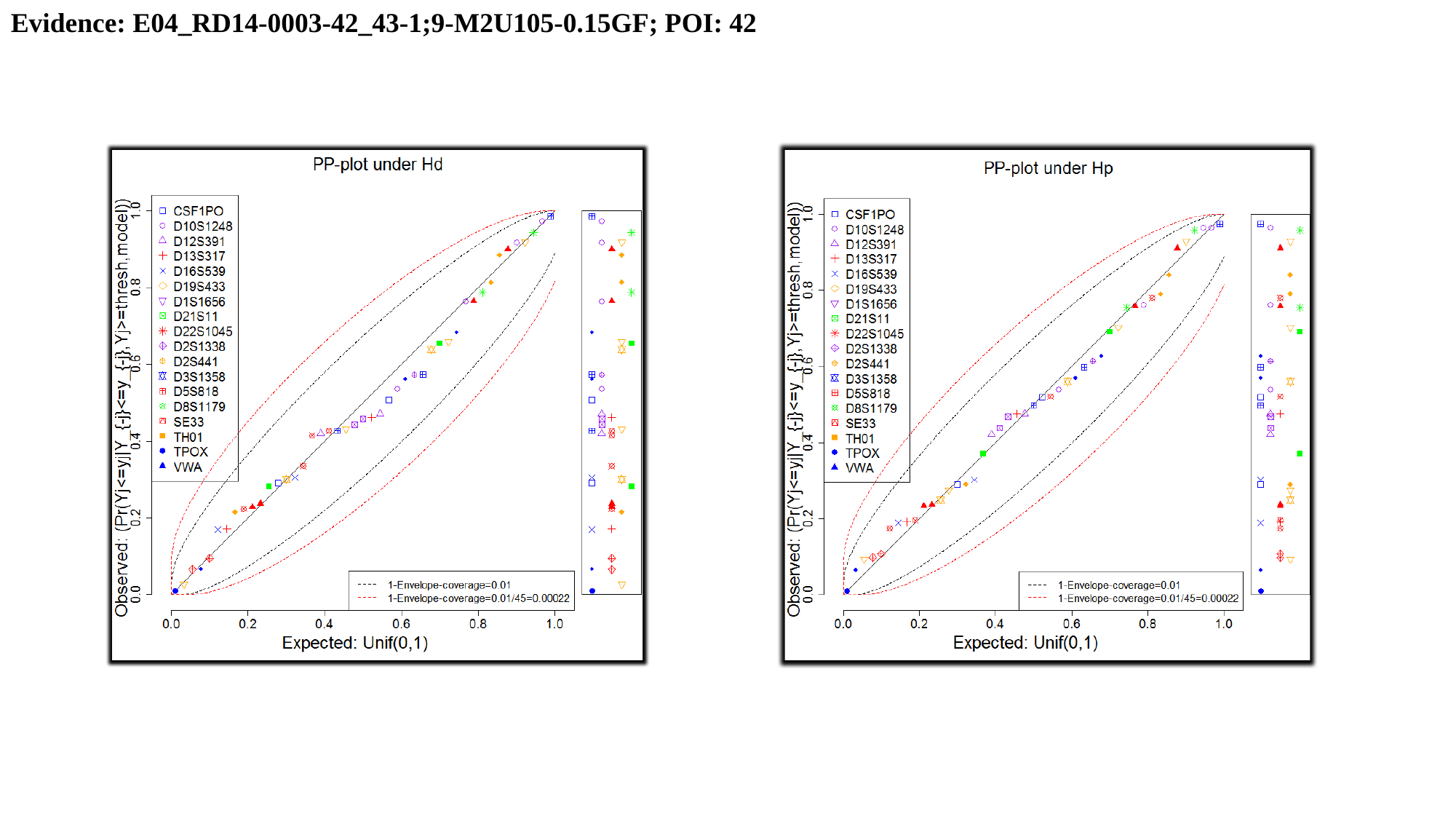

Evidence: E04_RD14-0003-42_43-1;9-M2U105-0.15GF; POI: 42

### Slide 4
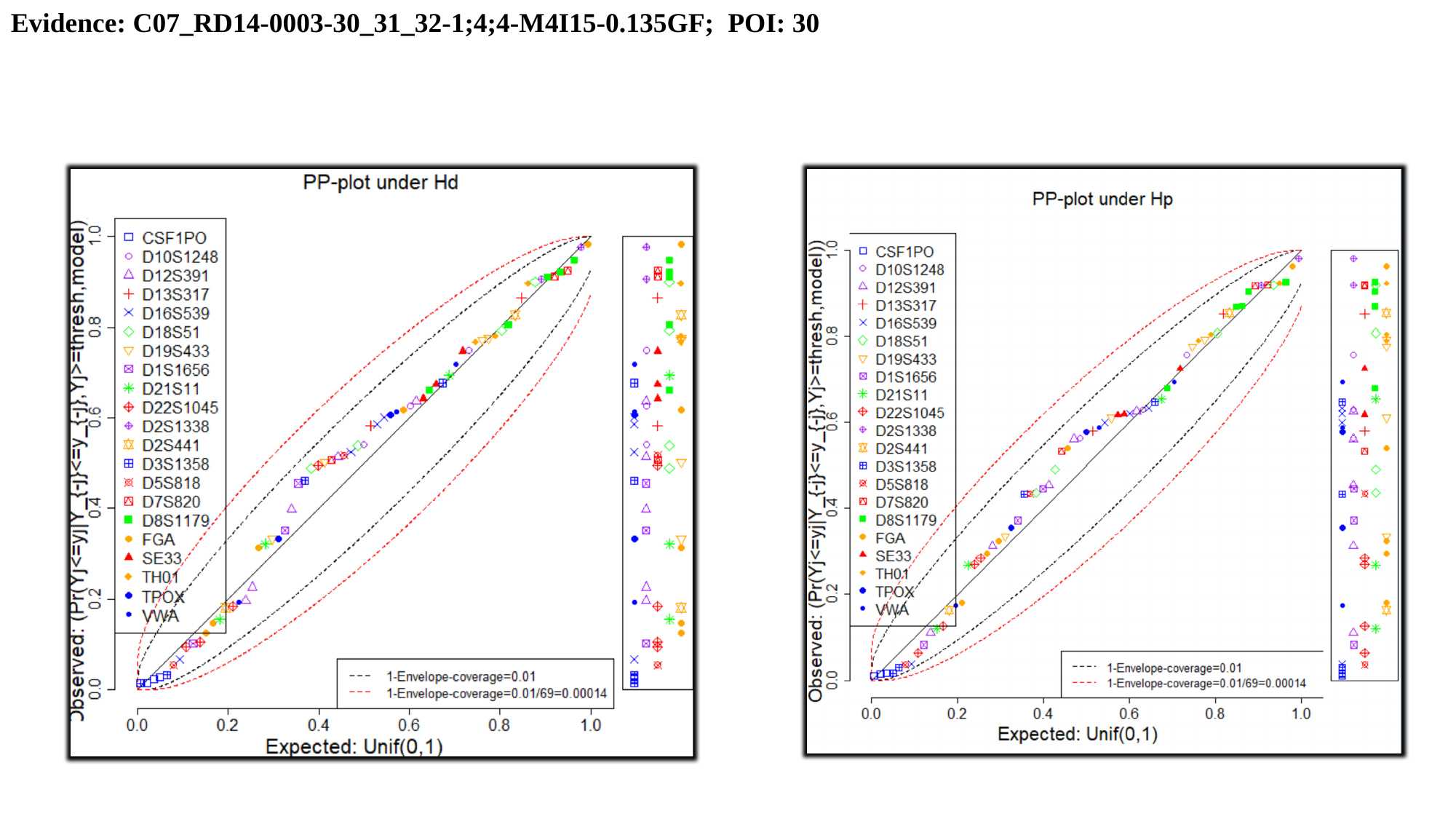

Evidence: C07_RD14-0003-30_31_32-1;4;4-M4I15-0.135GF; POI: 30

### Slide 5
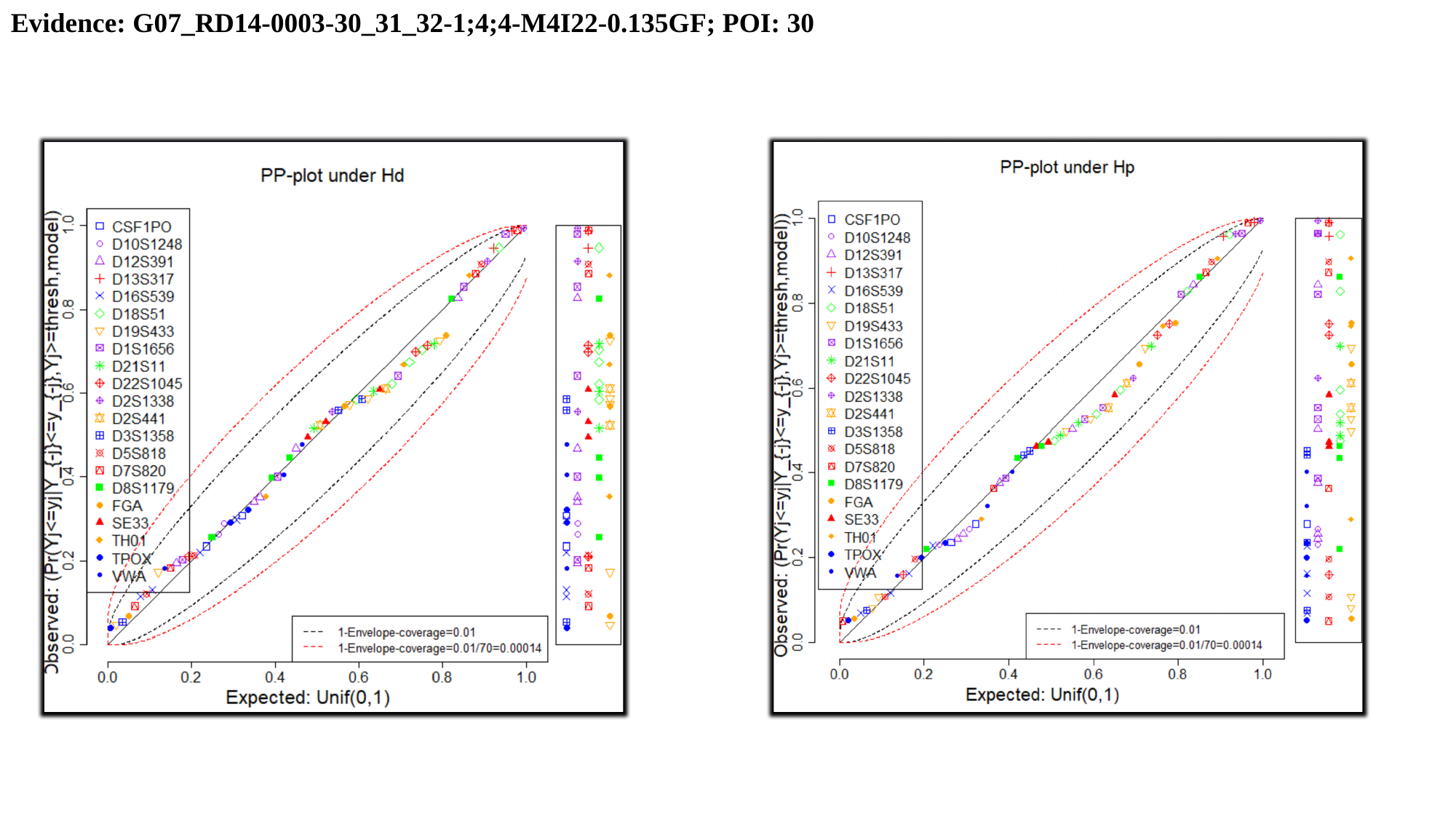

Evidence: G07_RD14-0003-30_31_32-1;4;4-M4I22-0.135GF; POI: 30
