## Additional File 4 for "Examining Discrimination Performance and Likelihood Ratio Values for Two Different Likelihood Ratio Systems Using the Provedit Dataset"

### Slide 1
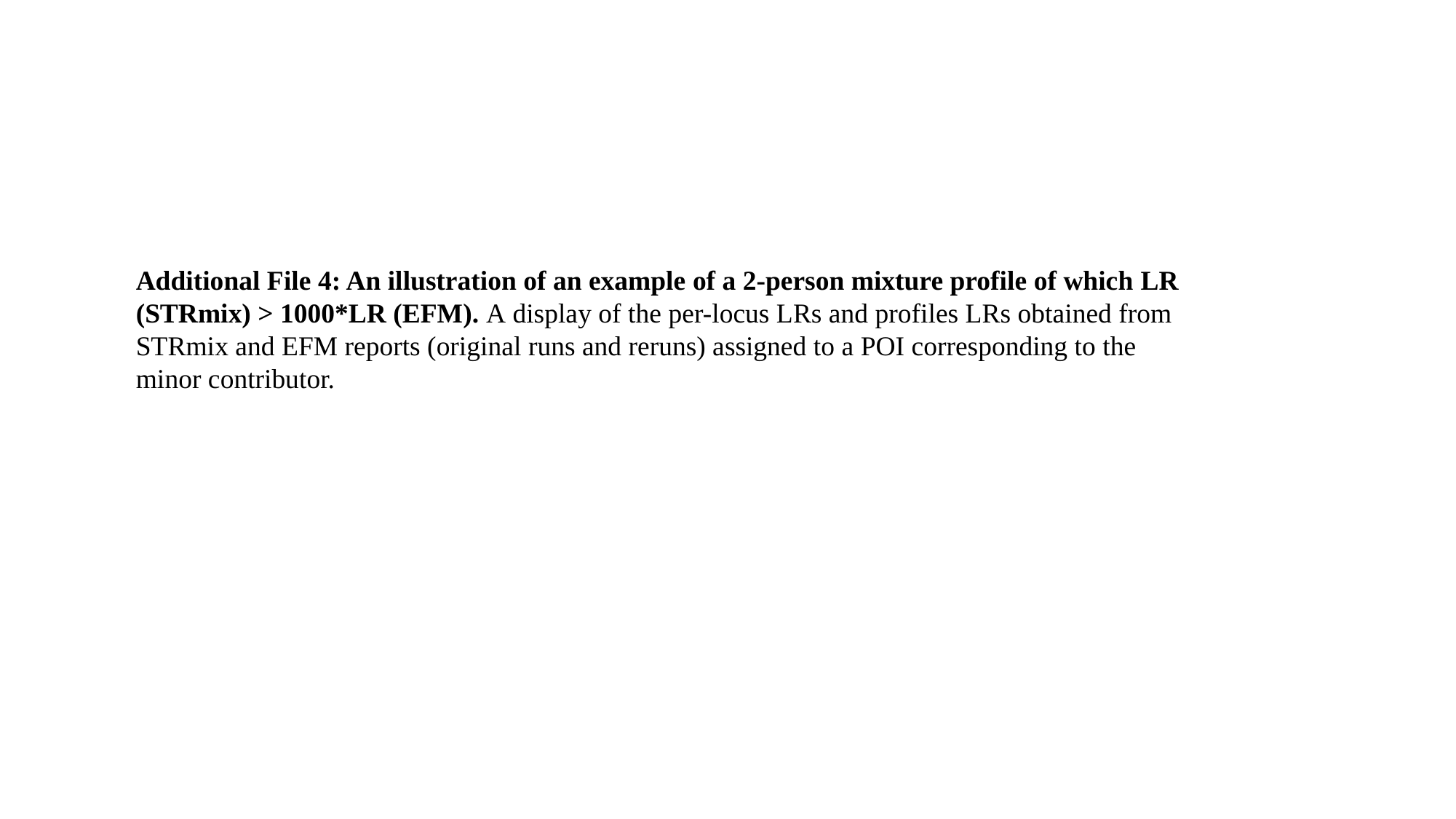

Additional File 4: An illustration of an example of a 2-person mixture profile of which LR (STRmix) > 1000*LR (EFM). A display of the per-locus LRs and profiles LRs obtained from STRmix and EFM reports (original runs and reruns) assigned to a POI corresponding to the minor contributor.

### Slide 2
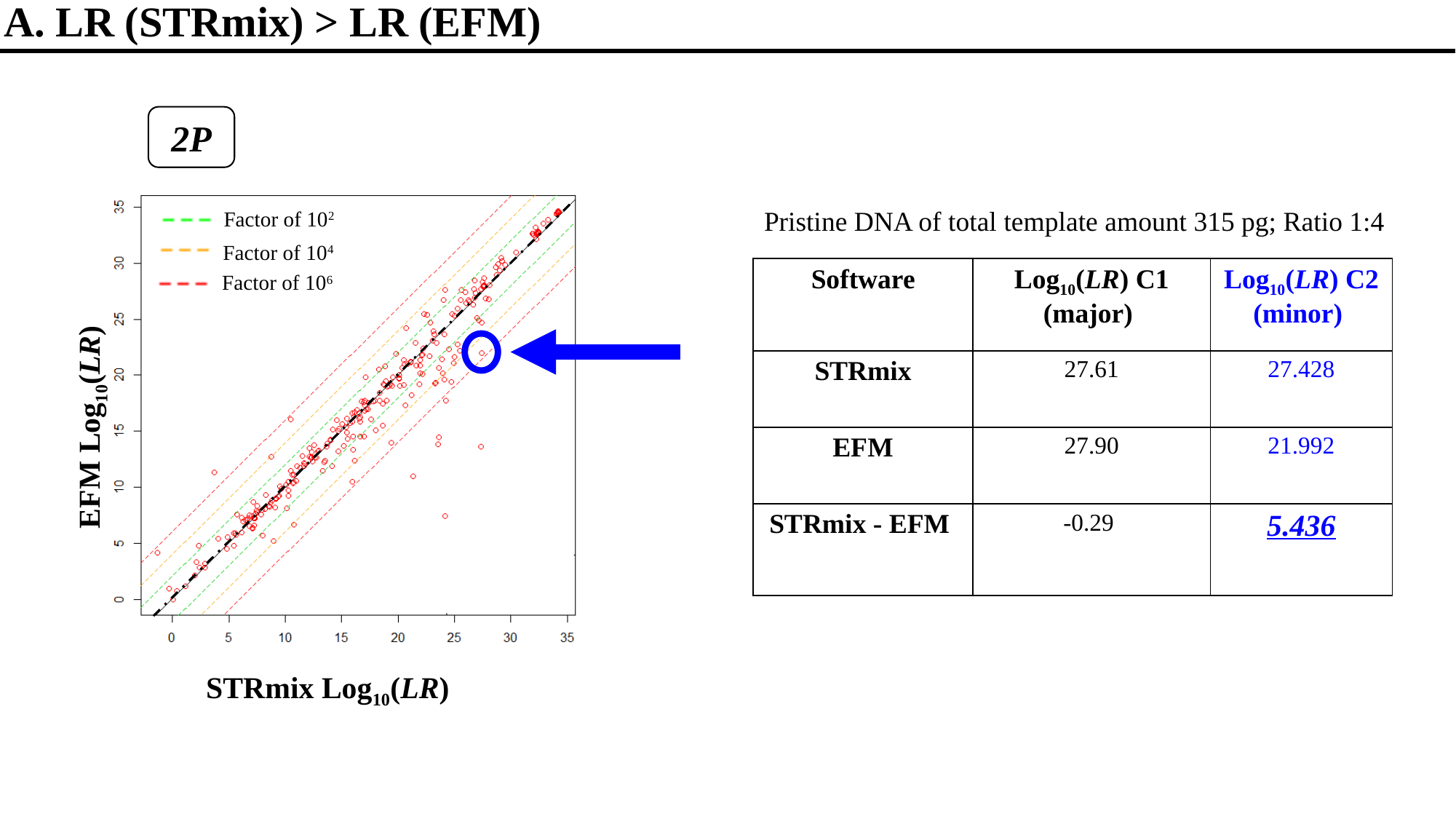

A. LR (STRmix) > LR (EFM)
2P
Pristine DNA of total template amount 315 pg; Ratio 1:4
Factor of 10^2^
Factor of 10^4^
Factor of 10^6^
| Software | Log_10_(LR) C1 (major) | Log_10_(LR) C2 (minor) |
| --- | --- | --- |
| STRmix | 27.61 | 27.428 |
| EFM | 27.90 | 21.992 |
| STRmix - EFM | -0.29 | 5.436 |
EFM Log_10_(LR)
STRmix Log_10_(LR)

### Slide 3
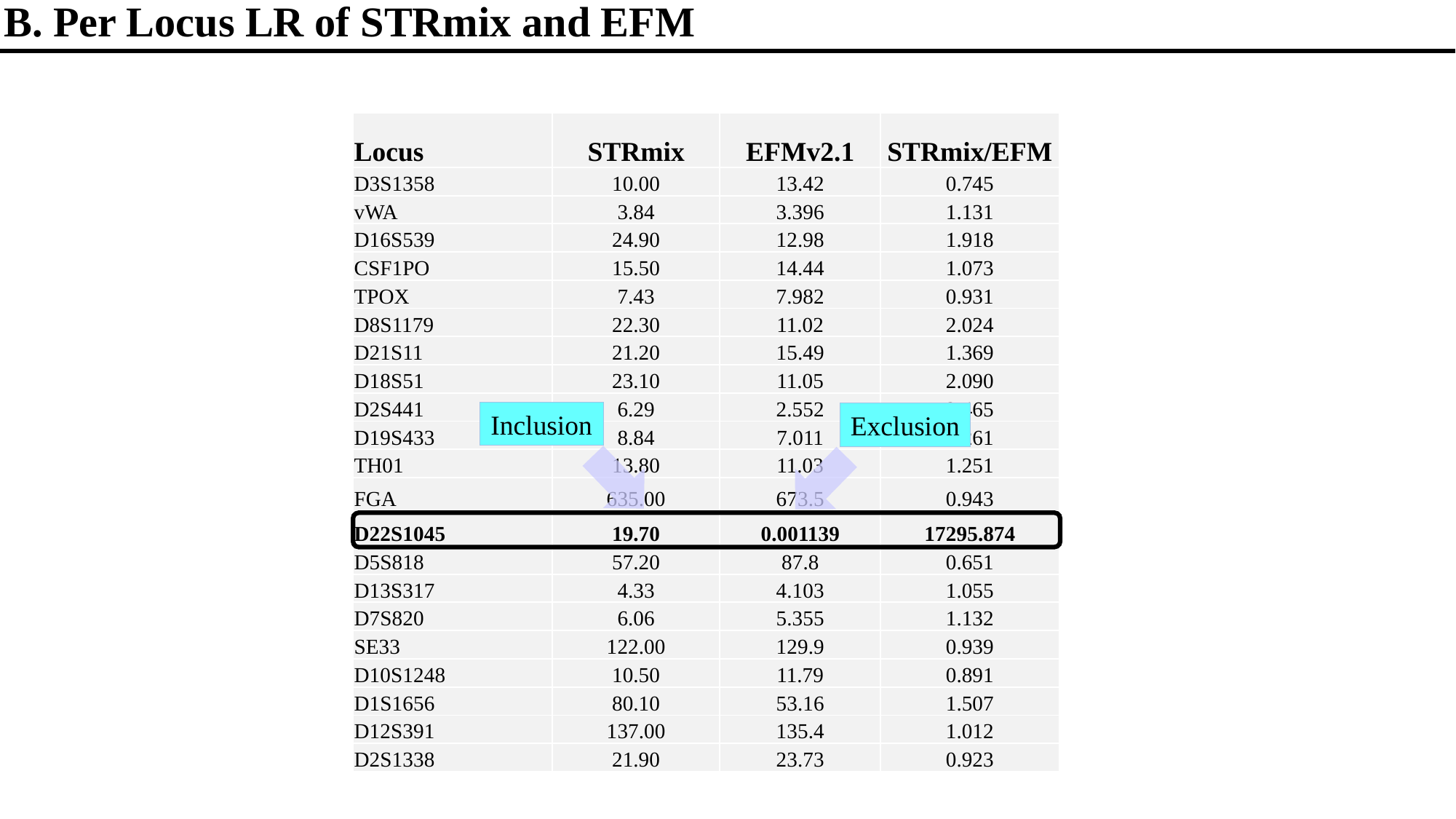

B. Per Locus LR of STRmix and EFM
| Locus | STRmix | EFMv2.1 | STRmix/EFM |
| --- | --- | --- | --- |
| D3S1358 | 10.00 | 13.42 | 0.745 |
| vWA | 3.84 | 3.396 | 1.131 |
| D16S539 | 24.90 | 12.98 | 1.918 |
| CSF1PO | 15.50 | 14.44 | 1.073 |
| TPOX | 7.43 | 7.982 | 0.931 |
| D8S1179 | 22.30 | 11.02 | 2.024 |
| D21S11 | 21.20 | 15.49 | 1.369 |
| D18S51 | 23.10 | 11.05 | 2.090 |
| D2S441 | 6.29 | 2.552 | 2.465 |
| D19S433 | 8.84 | 7.011 | 1.261 |
| TH01 | 13.80 | 11.03 | 1.251 |
| FGA | 635.00 | 673.5 | 0.943 |
| D22S1045 | 19.70 | 0.001139 | 17295.874 |
| D5S818 | 57.20 | 87.8 | 0.651 |
| D13S317 | 4.33 | 4.103 | 1.055 |
| D7S820 | 6.06 | 5.355 | 1.132 |
| SE33 | 122.00 | 129.9 | 0.939 |
| D10S1248 | 10.50 | 11.79 | 0.891 |
| D1S1656 | 80.10 | 53.16 | 1.507 |
| D12S391 | 137.00 | 135.4 | 1.012 |
| D2S1338 | 21.90 | 23.73 | 0.923 |
Inclusion
Exclusion

### Slide 4
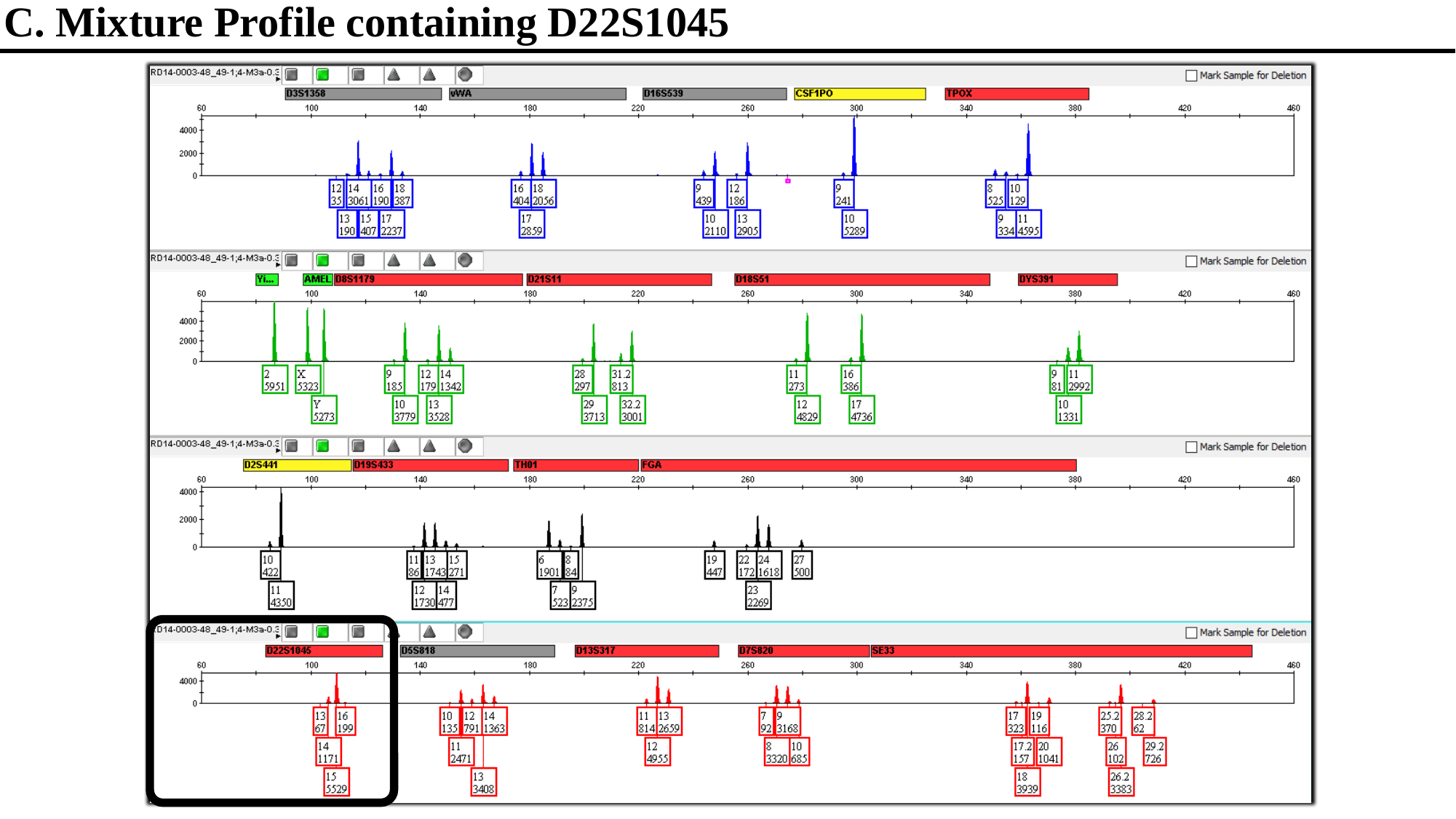

C. Mixture Profile containing D22S1045

### Slide 5
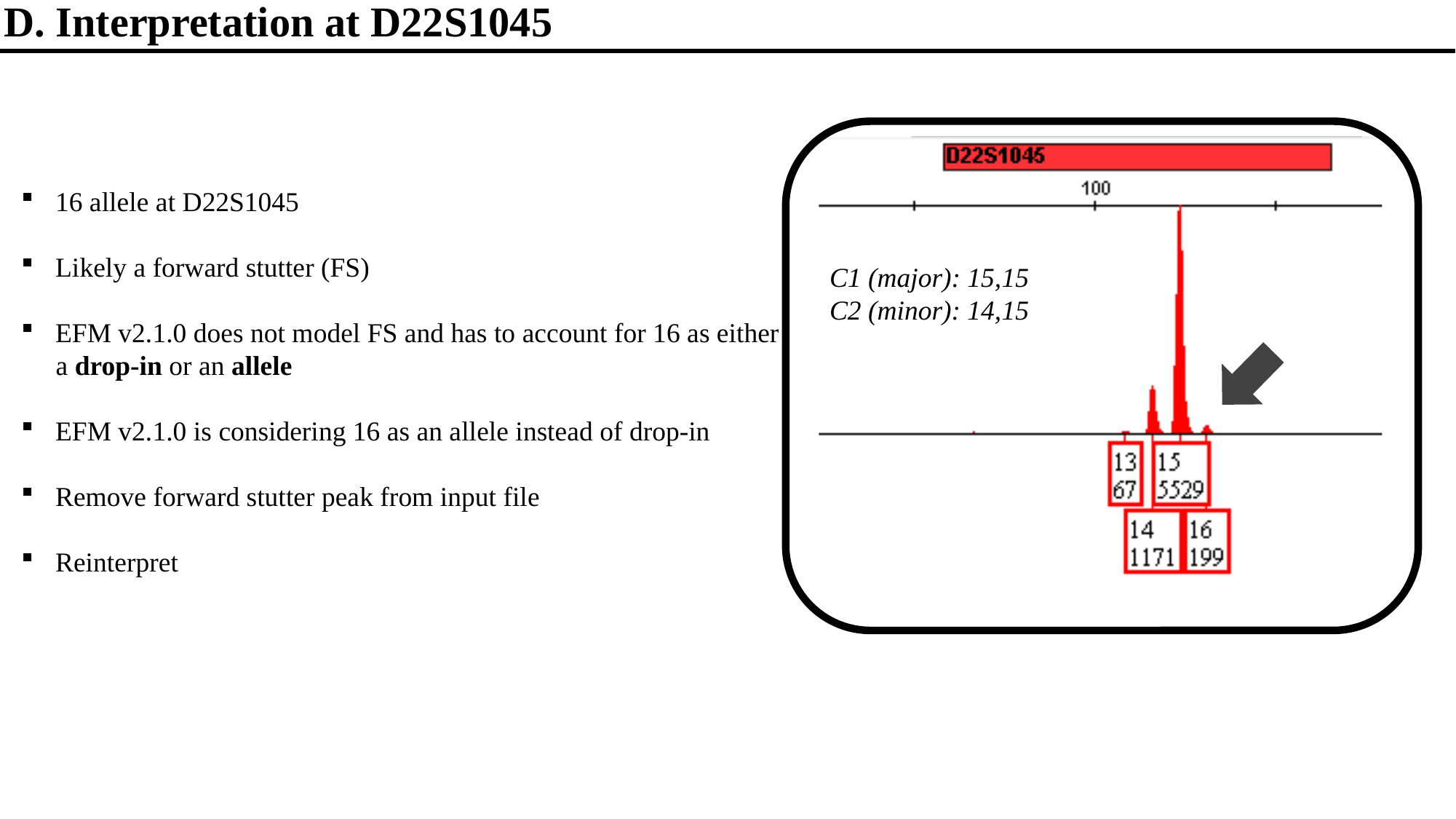

D. Interpretation at D22S1045
16 allele at D22S1045
Likely a forward stutter (FS)
EFM v2.1.0 does not model FS and has to account for 16 as either
 a drop-in or an allele
EFM v2.1.0 is considering 16 as an allele instead of drop-in
Remove forward stutter peak from input file
Reinterpret
C1 (major): 15,15
C2 (minor): 14,15

### Slide 6
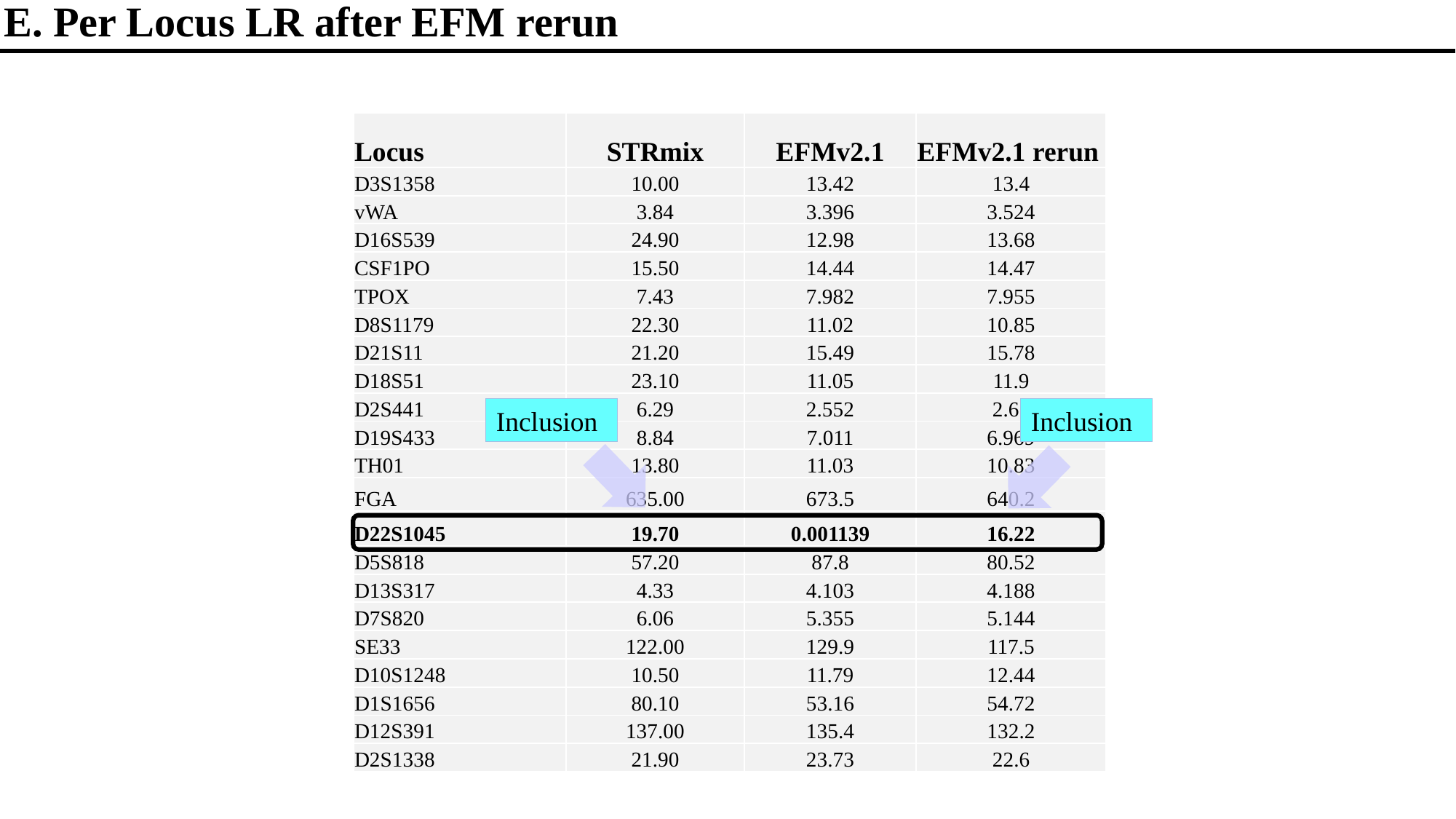

E. Per Locus LR after EFM rerun
| Locus | STRmix | EFMv2.1 | EFMv2.1 rerun |
| --- | --- | --- | --- |
| D3S1358 | 10.00 | 13.42 | 13.4 |
| vWA | 3.84 | 3.396 | 3.524 |
| D16S539 | 24.90 | 12.98 | 13.68 |
| CSF1PO | 15.50 | 14.44 | 14.47 |
| TPOX | 7.43 | 7.982 | 7.955 |
| D8S1179 | 22.30 | 11.02 | 10.85 |
| D21S11 | 21.20 | 15.49 | 15.78 |
| D18S51 | 23.10 | 11.05 | 11.9 |
| D2S441 | 6.29 | 2.552 | 2.65 |
| D19S433 | 8.84 | 7.011 | 6.969 |
| TH01 | 13.80 | 11.03 | 10.83 |
| FGA | 635.00 | 673.5 | 640.2 |
| D22S1045 | 19.70 | 0.001139 | 16.22 |
| D5S818 | 57.20 | 87.8 | 80.52 |
| D13S317 | 4.33 | 4.103 | 4.188 |
| D7S820 | 6.06 | 5.355 | 5.144 |
| SE33 | 122.00 | 129.9 | 117.5 |
| D10S1248 | 10.50 | 11.79 | 12.44 |
| D1S1656 | 80.10 | 53.16 | 54.72 |
| D12S391 | 137.00 | 135.4 | 132.2 |
| D2S1338 | 21.90 | 23.73 | 22.6 |
Inclusion
Inclusion

### Slide 7
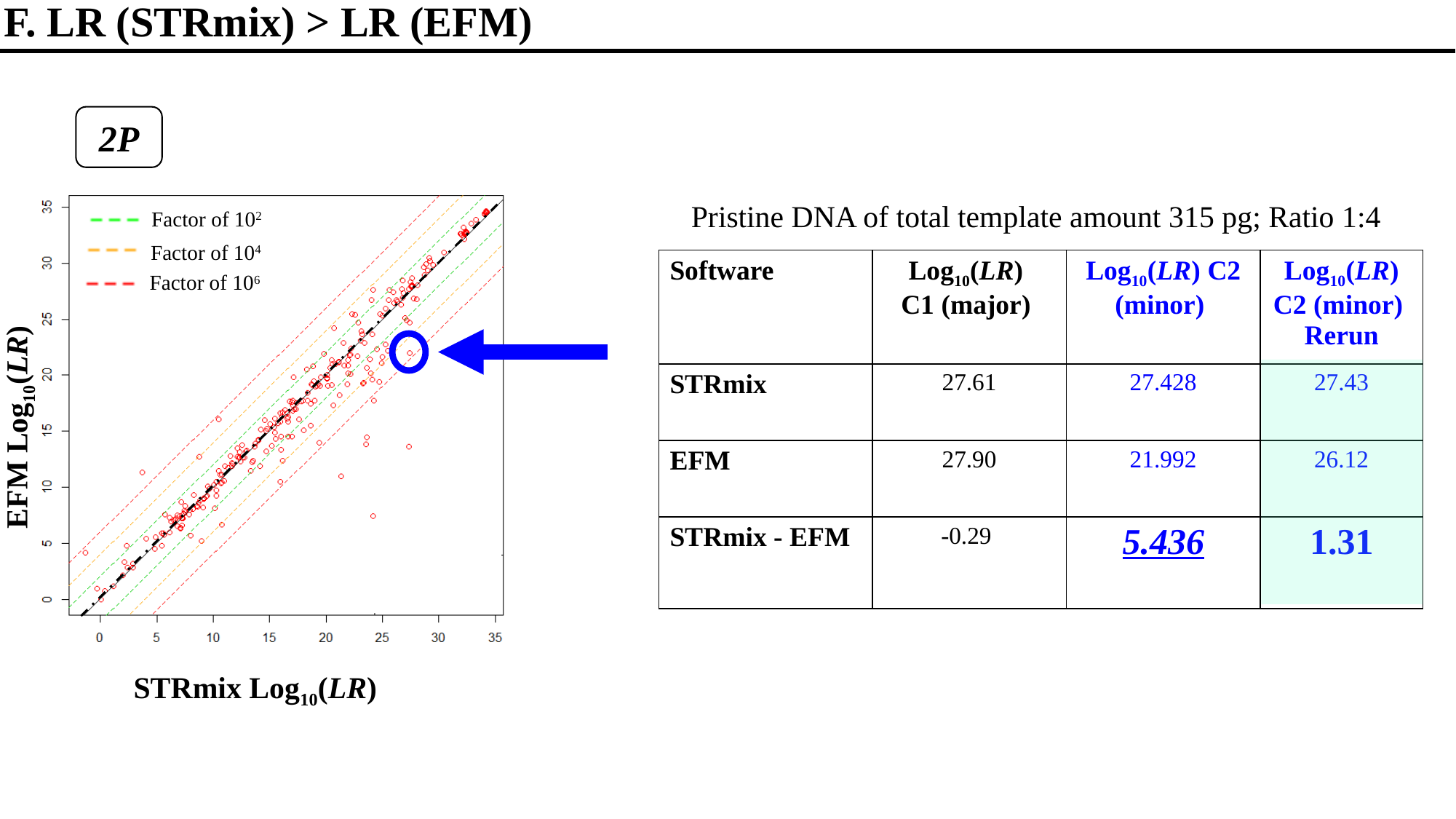

F. LR (STRmix) > LR (EFM)
2P
Pristine DNA of total template amount 315 pg; Ratio 1:4
Factor of 10^2^
Factor of 10^4^
Factor of 10^6^
2P
| Software | Log_10_(LR) C1 (major) | Log_10_(LR) C2 (minor) | Log_10_(LR) C2 (minor) Rerun |
| --- | --- | --- | --- |
| STRmix | 27.61 | 27.428 | 27.43 |
| EFM | 27.90 | 21.992 | 26.12 |
| STRmix - EFM | -0.29 | 5.436 | 1.31 |
EFM Log_10_(LR)
STRmix Log_10_(LR)
