## Additional File 5 for "Examining Discrimination Performance and Likelihood Ratio Values for Two Different Likelihood Ratio Systems Using the Provedit Dataset"

### Slide 1
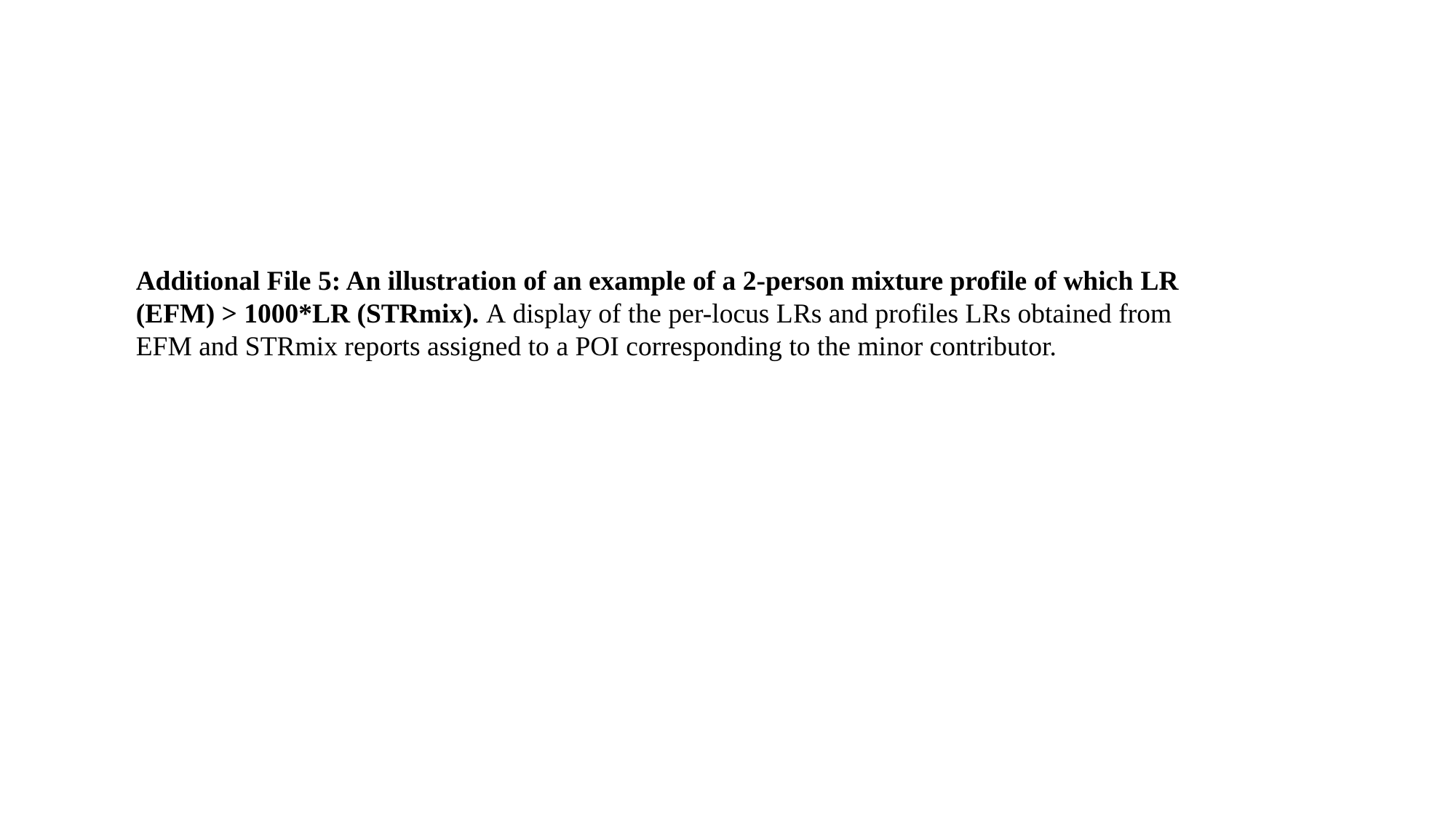

Additional File 5: An illustration of an example of a 2-person mixture profile of which LR (EFM) > 1000*LR (STRmix). A display of the per-locus LRs and profiles LRs obtained from EFM and STRmix reports assigned to a POI corresponding to the minor contributor.

### Slide 2
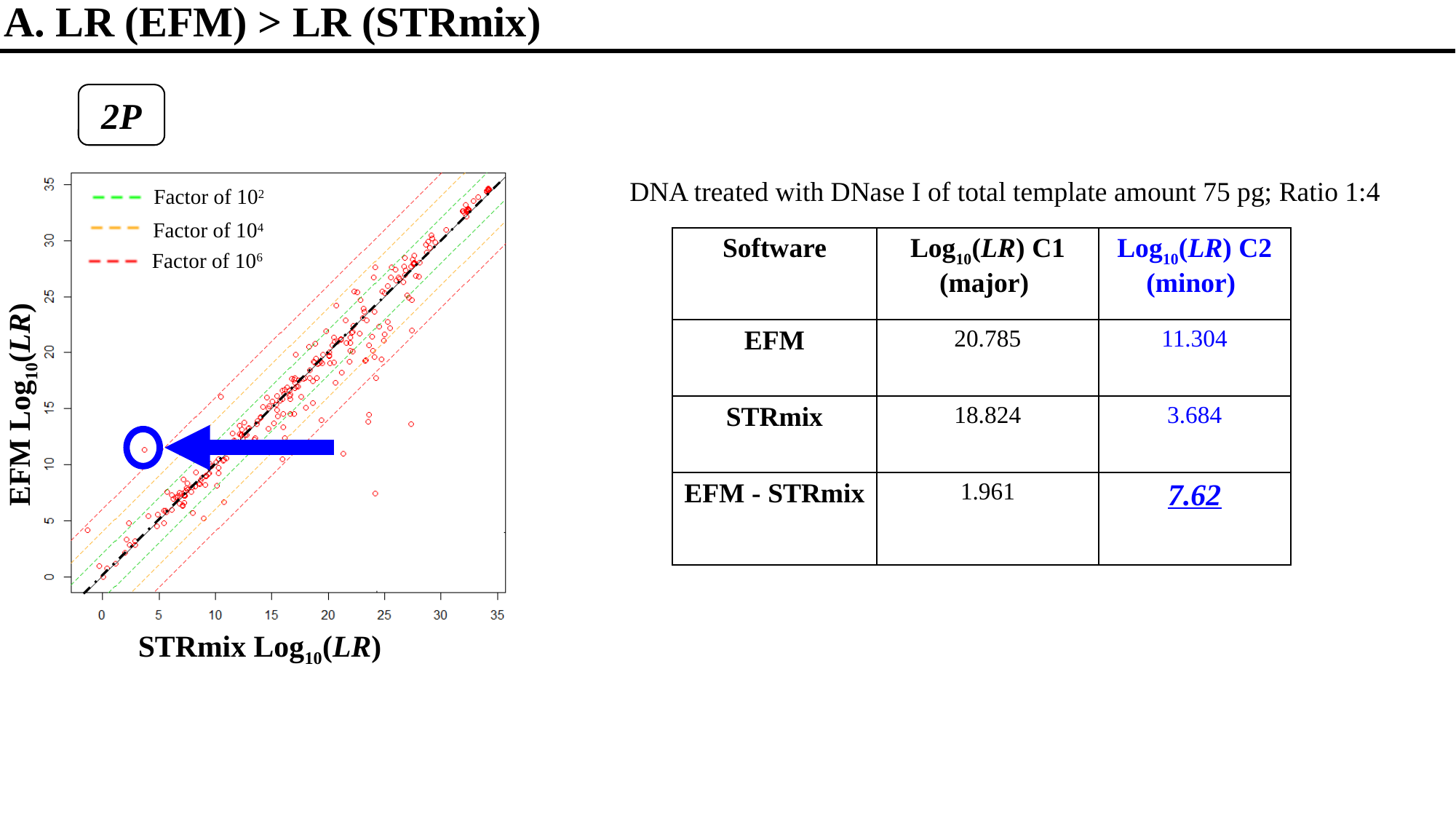

A. LR (EFM) > LR (STRmix)
2P
DNA treated with DNase I of total template amount 75 pg; Ratio 1:4
Factor of 10^2^
Factor of 10^4^
Factor of 10^6^
| Software | Log_10_(LR) C1 (major) | Log_10_(LR) C2 (minor) |
| --- | --- | --- |
| EFM | 20.785 | 11.304 |
| STRmix | 18.824 | 3.684 |
| EFM - STRmix | 1.961 | 7.62 |
EFM Log_10_(LR)
STRmix Log_10_(LR)

### Slide 3
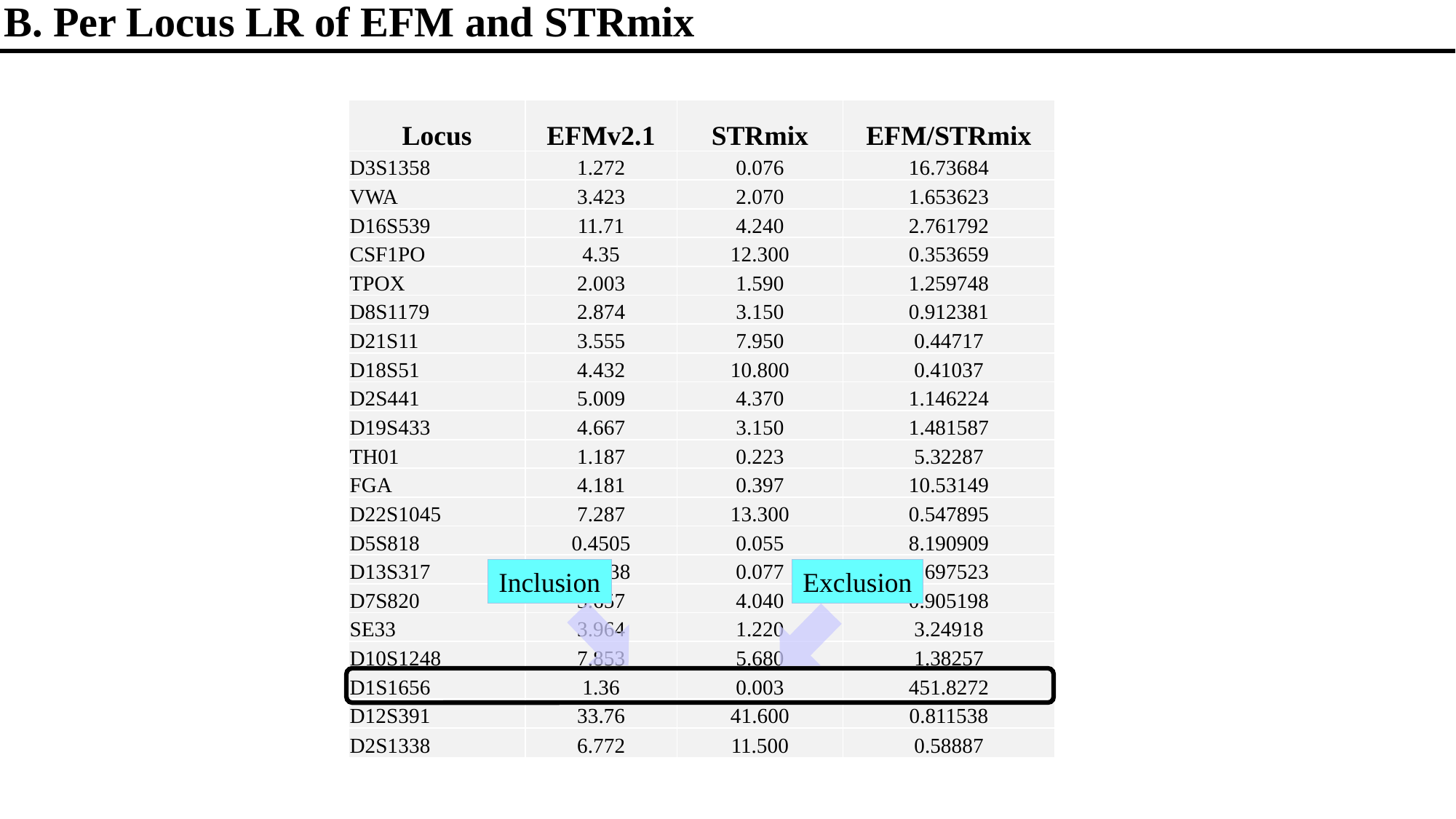

B. Per Locus LR of EFM and STRmix
| Locus | EFMv2.1 | STRmix | EFM/STRmix |
| --- | --- | --- | --- |
| D3S1358 | 1.272 | 0.076 | 16.73684 |
| VWA | 3.423 | 2.070 | 1.653623 |
| D16S539 | 11.71 | 4.240 | 2.761792 |
| CSF1PO | 4.35 | 12.300 | 0.353659 |
| TPOX | 2.003 | 1.590 | 1.259748 |
| D8S1179 | 2.874 | 3.150 | 0.912381 |
| D21S11 | 3.555 | 7.950 | 0.44717 |
| D18S51 | 4.432 | 10.800 | 0.41037 |
| D2S441 | 5.009 | 4.370 | 1.146224 |
| D19S433 | 4.667 | 3.150 | 1.481587 |
| TH01 | 1.187 | 0.223 | 5.32287 |
| FGA | 4.181 | 0.397 | 10.53149 |
| D22S1045 | 7.287 | 13.300 | 0.547895 |
| D5S818 | 0.4505 | 0.055 | 8.190909 |
| D13S317 | 0.7438 | 0.077 | 9.697523 |
| D7S820 | 3.657 | 4.040 | 0.905198 |
| SE33 | 3.964 | 1.220 | 3.24918 |
| D10S1248 | 7.853 | 5.680 | 1.38257 |
| D1S1656 | 1.36 | 0.003 | 451.8272 |
| D12S391 | 33.76 | 41.600 | 0.811538 |
| D2S1338 | 6.772 | 11.500 | 0.58887 |
Inclusion
Exclusion

### Slide 4
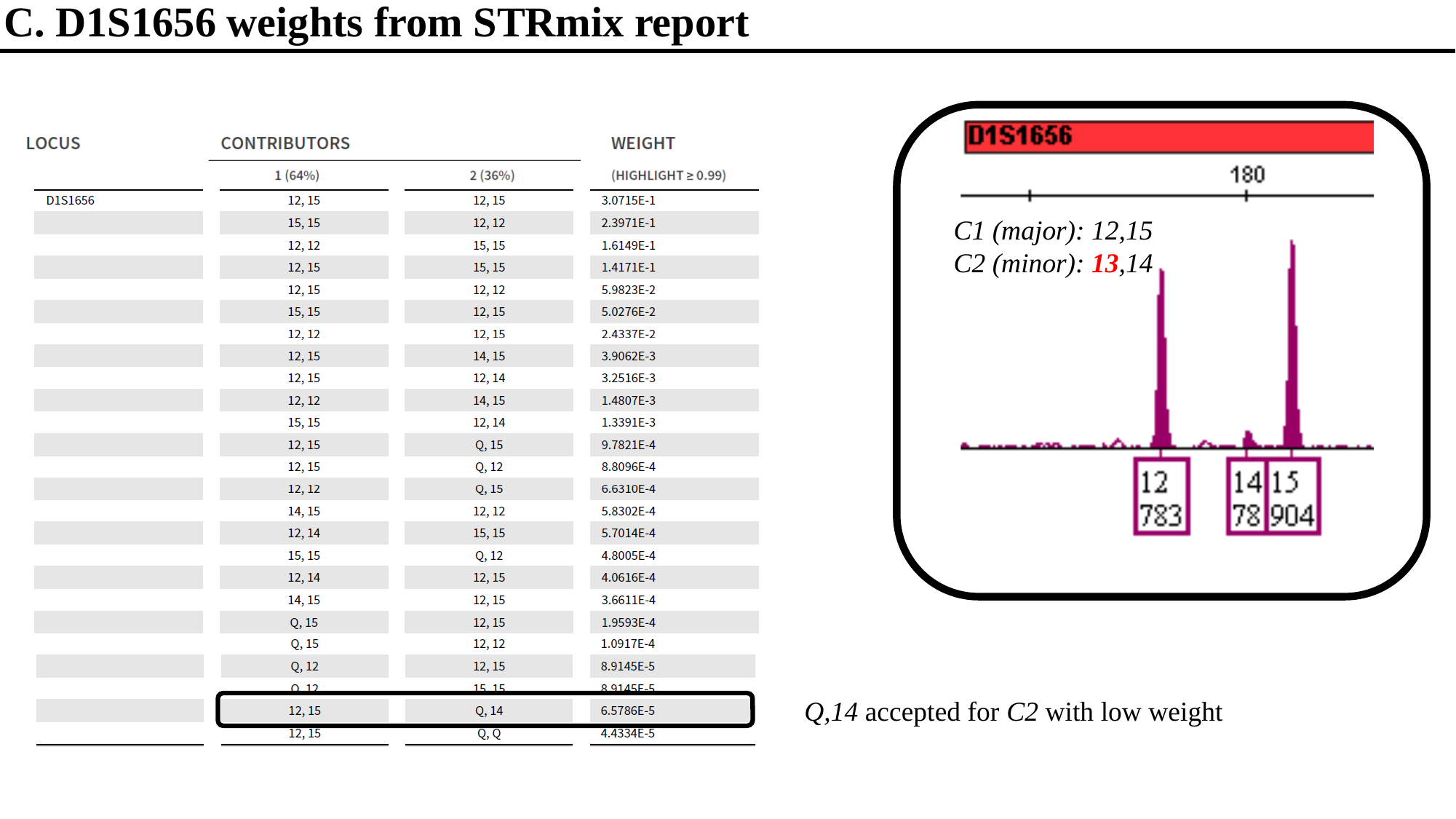

C. D1S1656 weights from STRmix report
C1 (major): 12,15
C2 (minor): 13,14
Q,14 accepted for C2 with low weight
