## Additional File 1 for "Examining Discrimination Performance and Likelihood Ratio Values for Two Different Likelihood Ratio Systems Using the Provedit Dataset"

### Slide 1
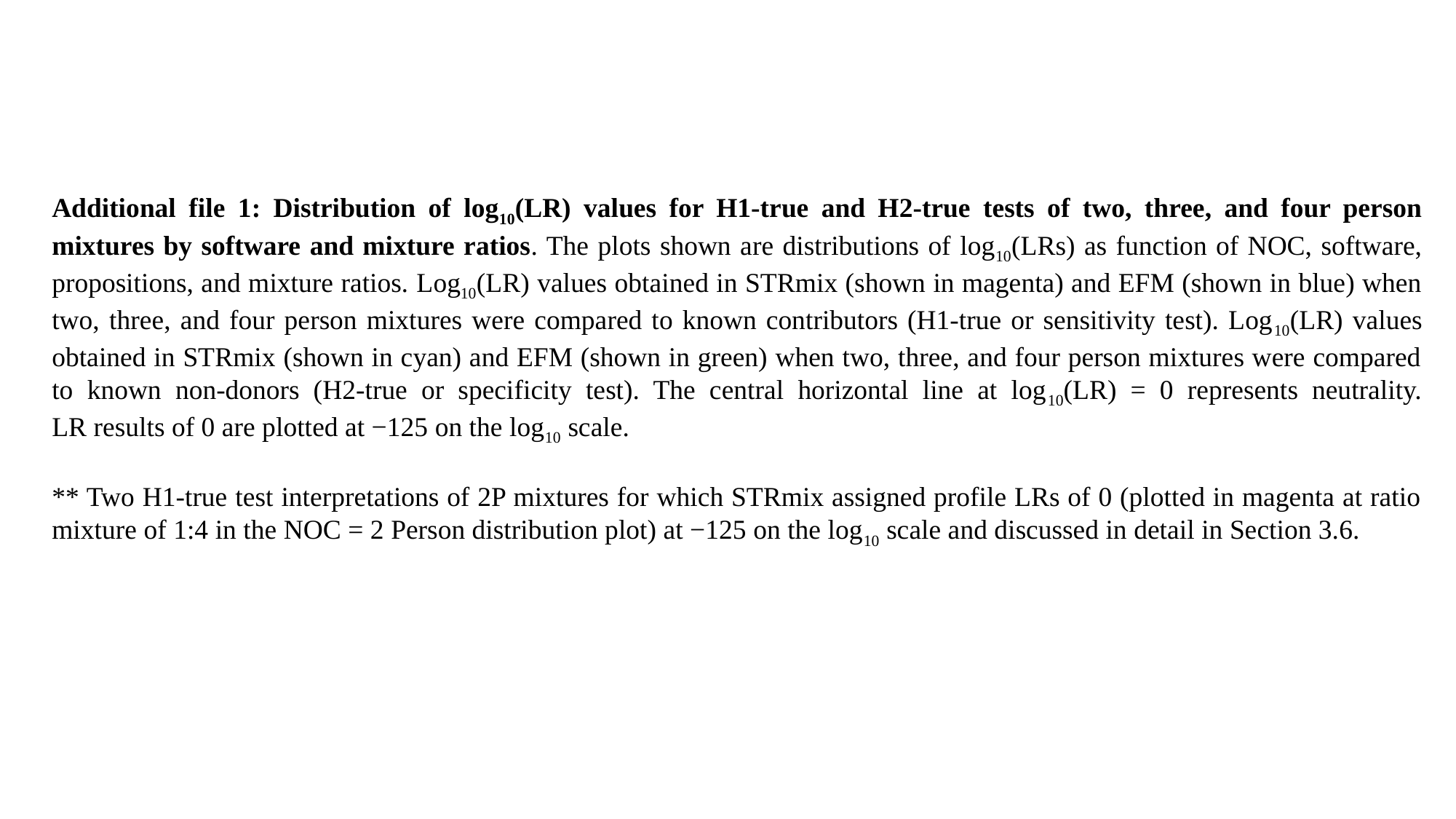

Additional file 1: Distribution of log_10_(LR) values for H1-true and H2-true tests of two, three, and four person mixtures by software and mixture ratios. The plots shown are distributions of log_10_(LRs) as function of NOC, software, propositions, and mixture ratios. Log_10_(LR) values obtained in STRmix (shown in magenta) and EFM (shown in blue) when two, three, and four person mixtures were compared to known contributors (H1-true or sensitivity test). Log_10_(LR) values obtained in STRmix (shown in cyan) and EFM (shown in green) when two, three, and four person mixtures were compared to known non-donors (H2-true or specificity test). The central horizontal line at log_10_(LR) = 0 represents neutrality. LR results of 0 are plotted at −125 on the log_10_ scale.
** Two H1-true test interpretations of 2P mixtures for which STRmix assigned profile LRs of 0 (plotted in magenta at ratio mixture of 1:4 in the NOC = 2 Person distribution plot) at −125 on the log_10_ scale and discussed in detail in Section 3.6.

### Slide 2
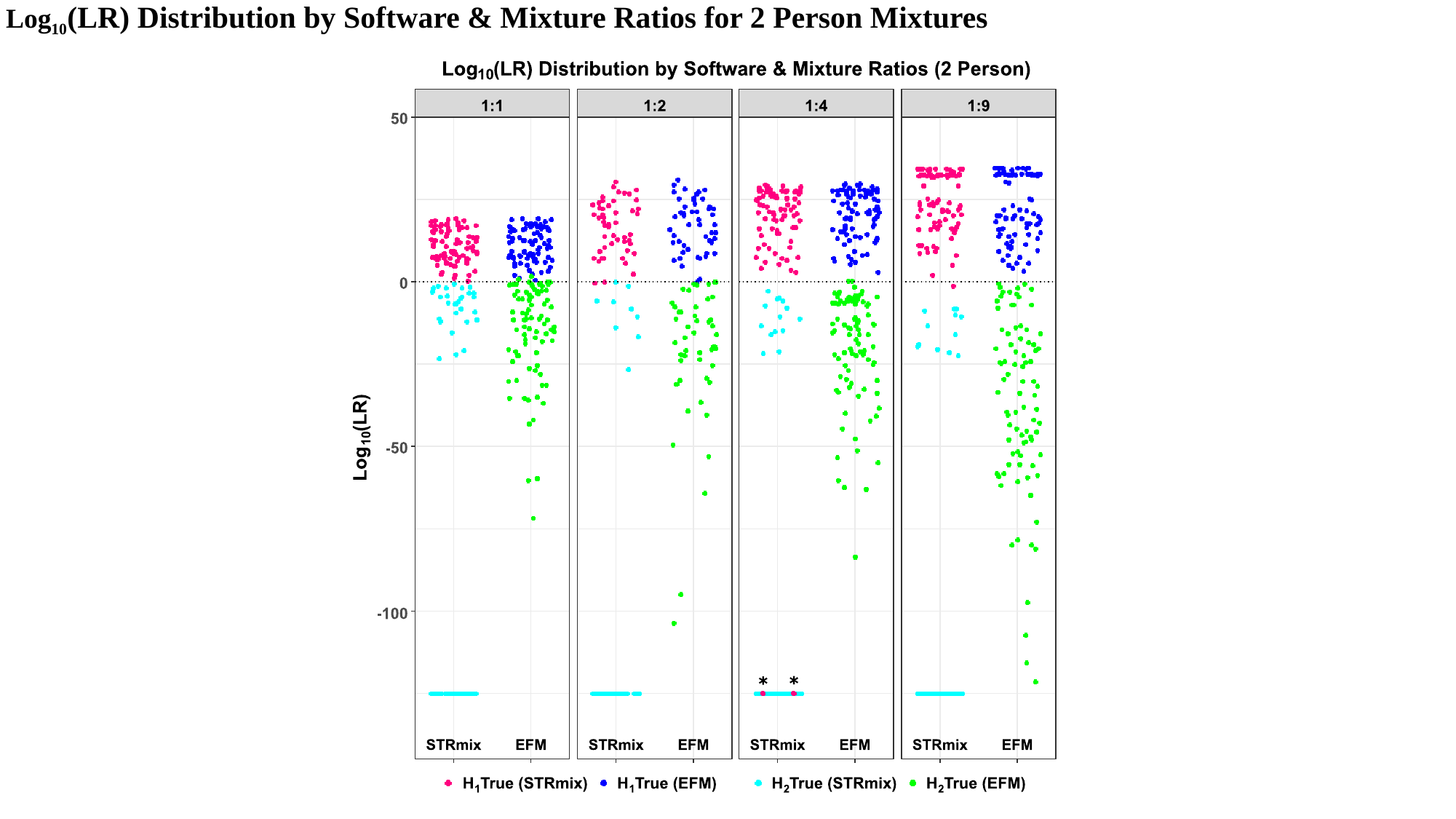

Log_10_(LR) Distribution by Software & Mixture Ratios for 2 Person Mixtures

### Slide 3
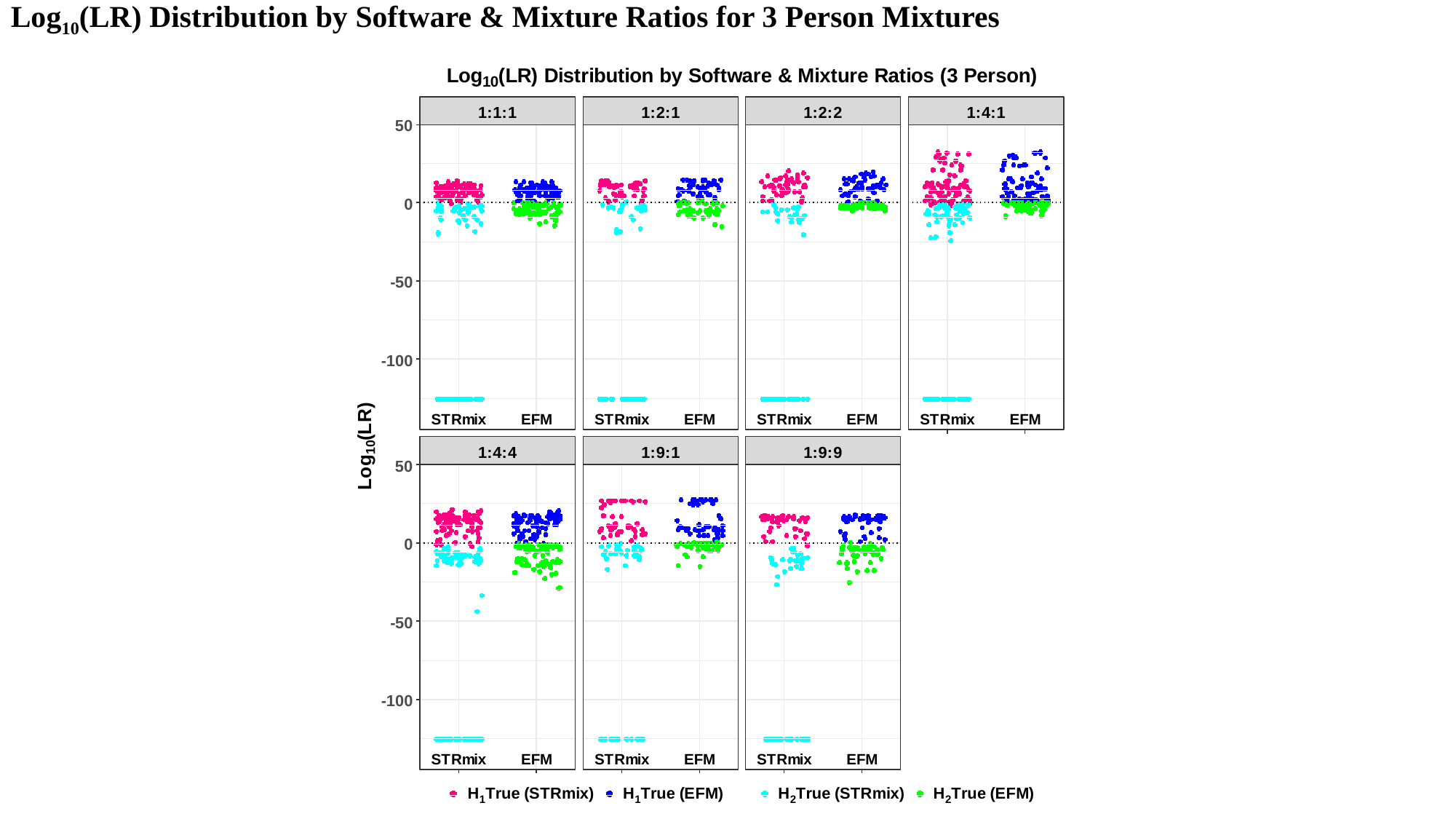

Log_10_(LR) Distribution by Software & Mixture Ratios for 3 Person Mixtures

### Slide 4
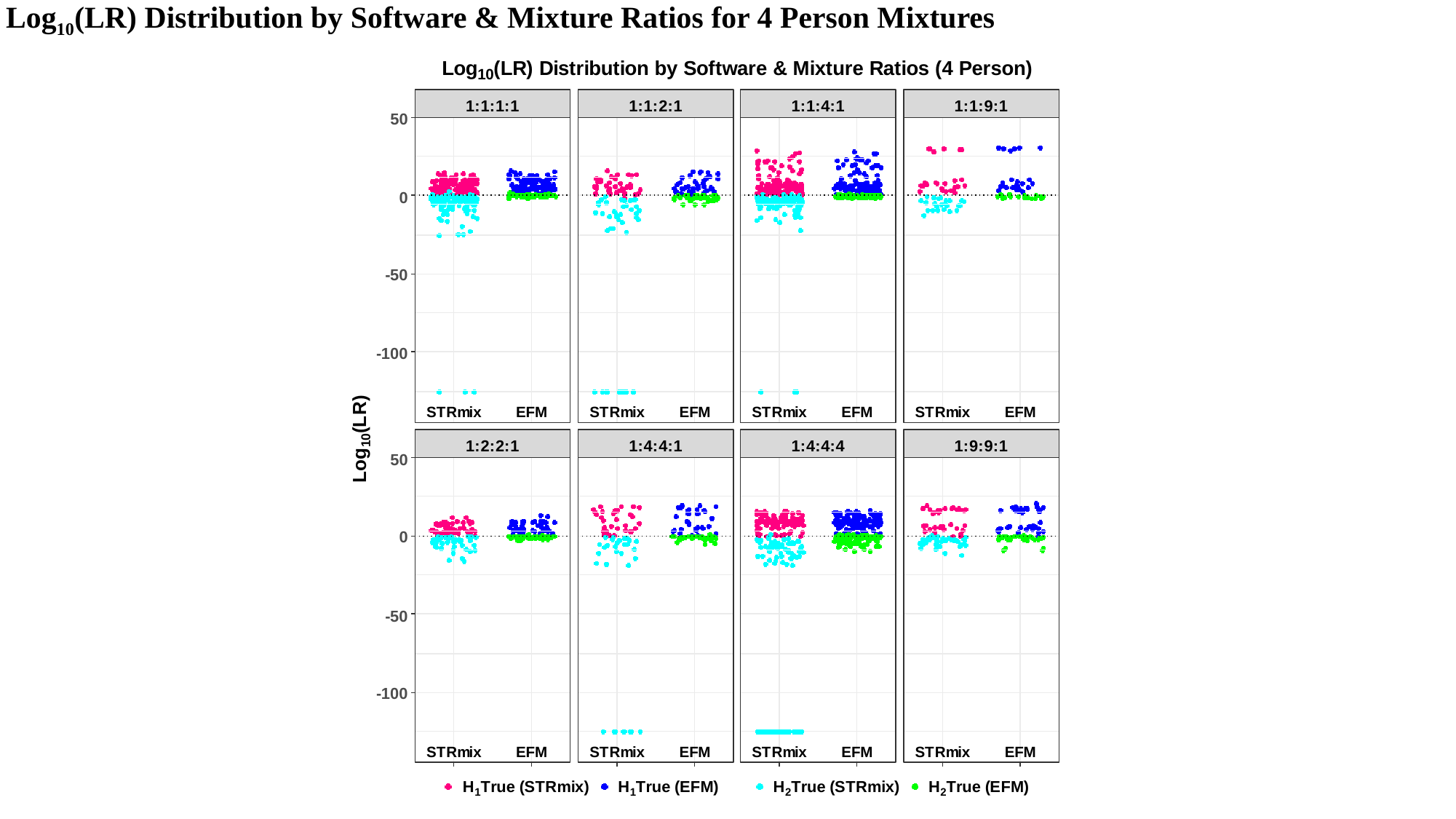

Log_10_(LR) Distribution by Software & Mixture Ratios for 4 Person Mixtures
