## Additional File 2 for "Examining Discrimination Performance and Likelihood Ratio Values for Two Different Likelihood Ratio Systems Using the Provedit Dataset"

### Slide 1
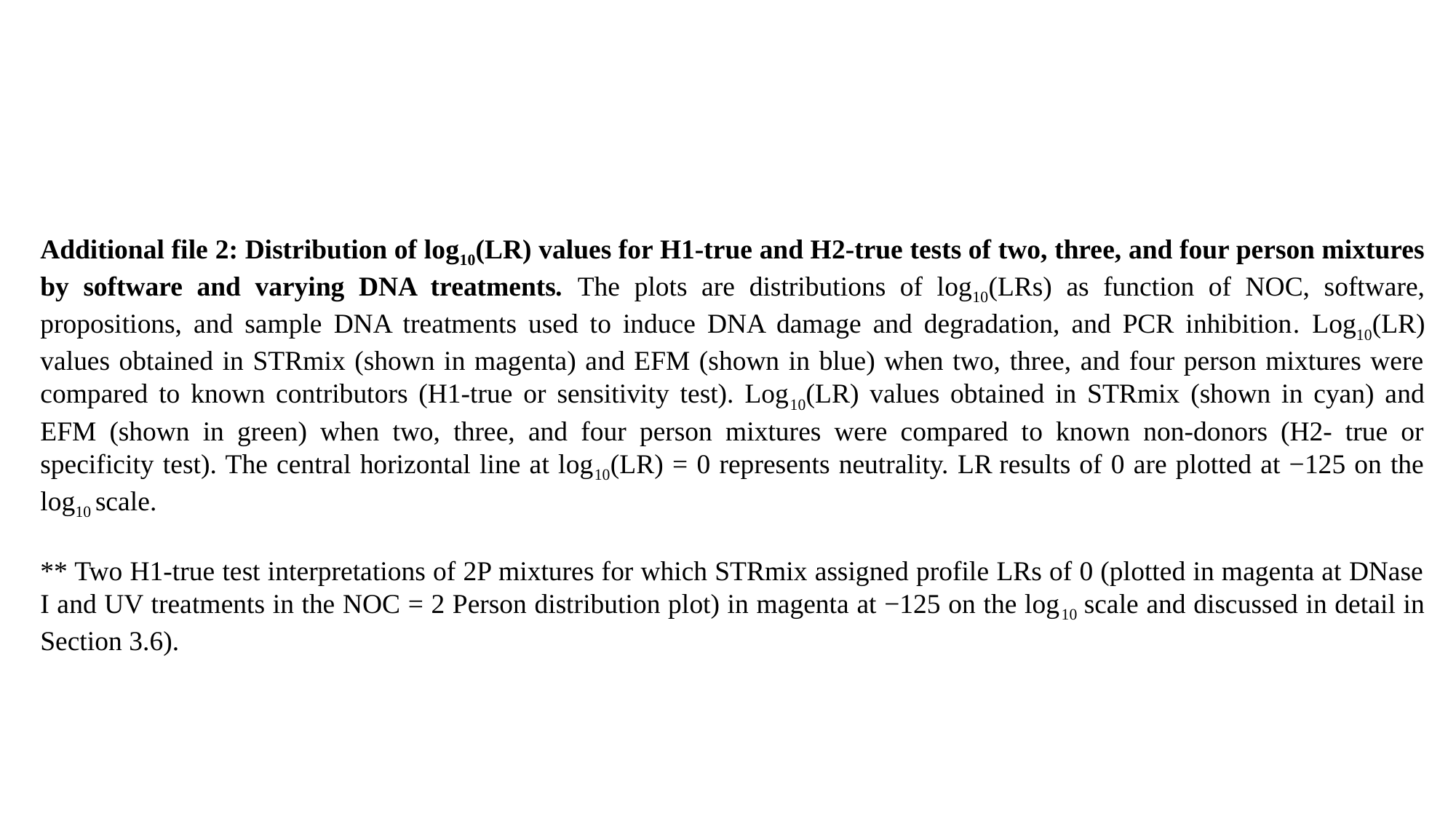

Additional file 2: Distribution of log_10_(LR) values for H1-true and H2-true tests of two, three, and four person mixtures by software and varying DNA treatments. The plots are distributions of log_10_(LRs) as function of NOC, software, propositions, and sample DNA treatments used to induce DNA damage and degradation, and PCR inhibition. Log_10_(LR) values obtained in STRmix (shown in magenta) and EFM (shown in blue) when two, three, and four person mixtures were compared to known contributors (H1-true or sensitivity test). Log_10_(LR) values obtained in STRmix (shown in cyan) and EFM (shown in green) when two, three, and four person mixtures were compared to known non-donors (H2- true or specificity test). The central horizontal line at log_10_(LR) = 0 represents neutrality. LR results of 0 are plotted at −125 on the log_10_ scale.
** Two H1-true test interpretations of 2P mixtures for which STRmix assigned profile LRs of 0 (plotted in magenta at DNase I and UV treatments in the NOC = 2 Person distribution plot) in magenta at −125 on the log_10_ scale and discussed in detail in Section 3.6).

### Slide 2
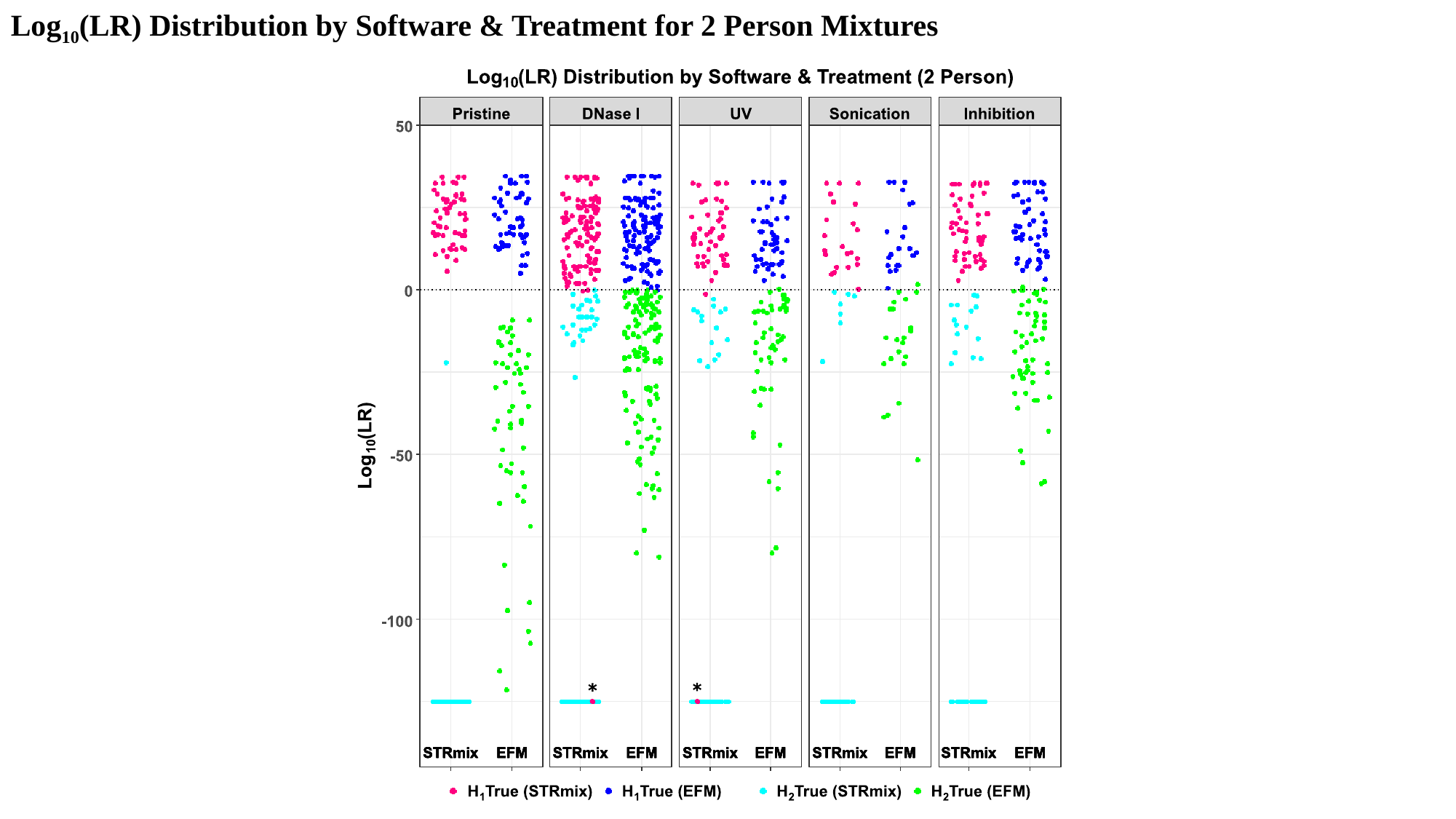

Log_10_(LR) Distribution by Software & Treatment for 2 Person Mixtures

### Slide 3
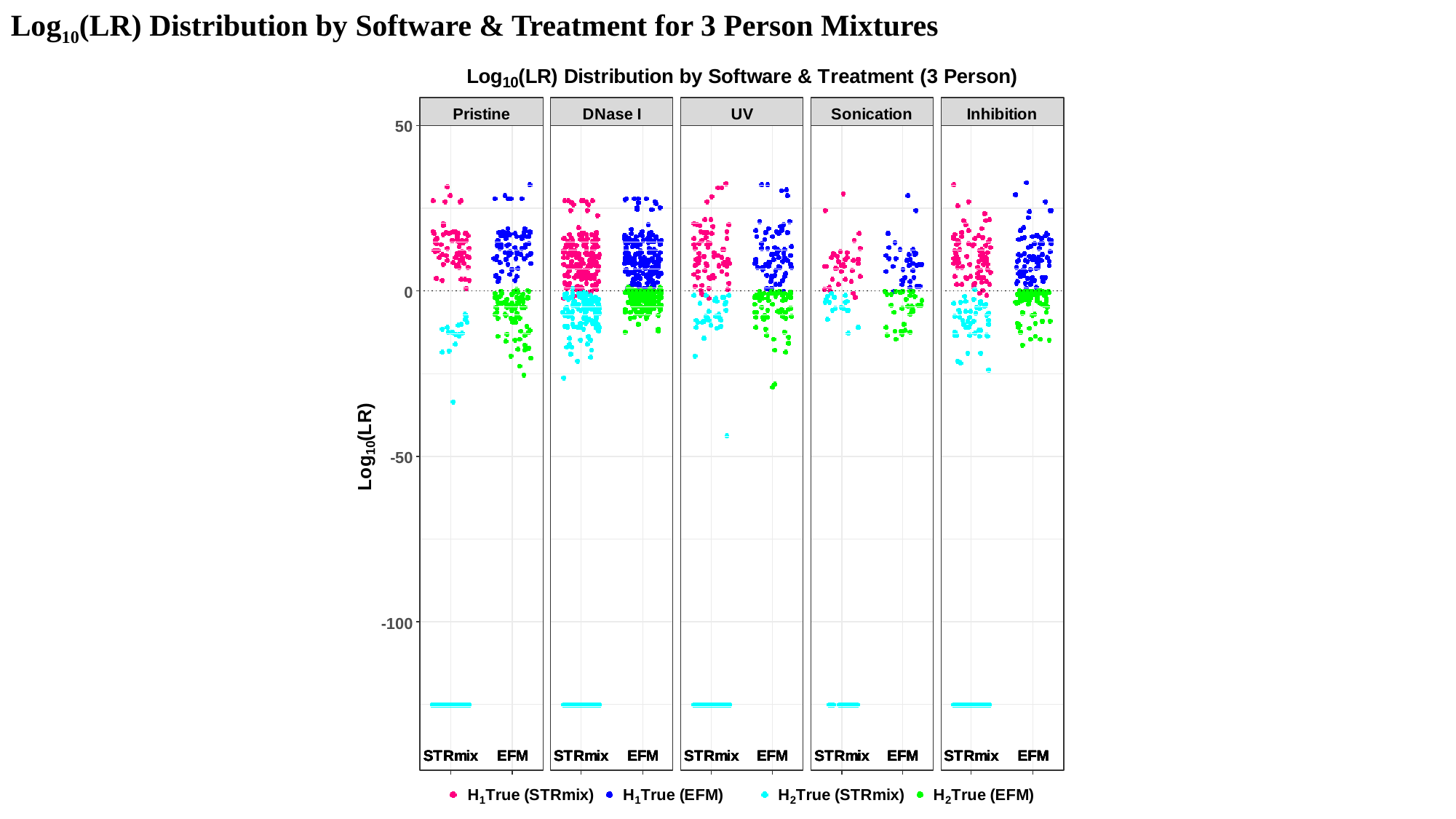

Log_10_(LR) Distribution by Software & Treatment for 3 Person Mixtures

### Slide 4
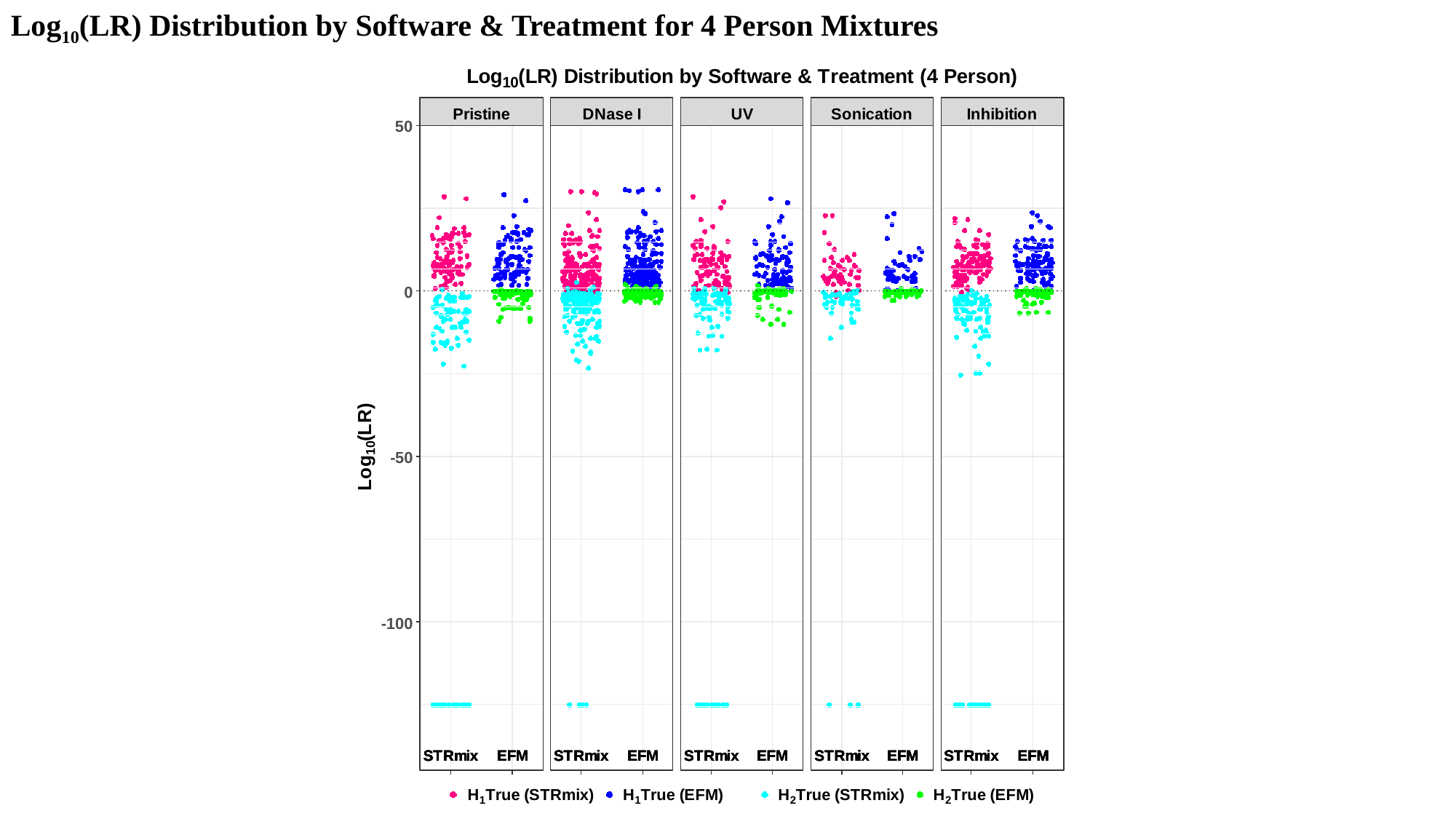

Log_10_(LR) Distribution by Software & Treatment for 4 Person Mixtures
